# Phylogeny structures species’ interactions in experimental ecological communities

**DOI:** 10.1101/2023.09.04.556236

**Authors:** Paula Lemos-Costa, Zachary R. Miller, Stefano Allesina

**Affiliations:** Department of Ecology & Evolution, University of Chicago and Department of Plant Biology, University of Illinois

## Abstract

The advent of molecular phylogenetics provided a new perspective on the structure and function of ecological communities. In particular, the hypothesis that traits responsible for species’ interactions are largely determined by shared evolutionary history has suggested the possibility of connecting the phylogeny of ecological communities to their functioning. However, statistical tests of this link have yielded mixed results. Here we propose a novel framework to test whether phylogeny influences the patterns of coexistence and abundance of species assemblages, and apply it to analyze data from large biodiversity-ecosystem functioning experiments. In our approach, phylogenetic trees are used to parameterize species’ interactions, which in turn determine the abundance of species in a specified assemblage. We use a maximum likelihood-based approach to score models parameterized with a given phylogenetic tree. To test whether evolutionary history structures interactions, we fit and score ensembles of randomized trees, allowing us to determine if phylogenetic information helps to predict species’ abundances. Moreover, we can determine the contribution of each branch of the tree to the likelihood, revealing particular clades in which interaction strengths are closely tied to phylogeny. We find strong evidence of phylogenetic signal across a range of published experiments and a variety of models. The flexibility of our framework permits incorporation of ecological information beyond phylogeny, such as functional groups or traits, and provides a principled way to test hypotheses about which factors shape the structure and function of ecological communities.

## Introduction

In 1837, Charles Darwin jotted *“I think”* and sketched a phylogenetic tree in his notebook, setting in motion a revolution in biology. Almost two centuries later, the notions of phylogeny and shared ancestry have a central place in biological research, from biochemistry to ecology. In community ecology, phylogeny was recognized early on as a powerful tool to organize our thinking about ecological communities, and the integration of phylogenetic information has promised to shed light on community assembly and patterns of species co-occurrence. Early studies relied on taxonomic information, focusing for example on species-to-genus ratios (Elton 1946; Simberloff 1970) as a proxy for competition; the idea that competition might leave a fingerprint in the taxonomic structure of ecological communities can be traced all the way back to Darwin: “As species of the same genus have usually, though by no means invariably, some similarity in habits and constitution, and always in structure, the struggle will generally be more severe between species of the same genus, when they come into competition with each other, than between species of distinct genera” (Darwin 2004).

Decades later, the availability of molecular-based phylogenies has allowed for more thorough, quantitative testing of this hypothesis. Under the umbrella of “community phylogenetics”, many studies have sought to detect the signatures of community assembly processes in the phylogenetic structure of natural communities. For example, in an influential study, Webb (2000) investigated the role of shared evolutionary history in structuring rain forest tree communities. By comparing the phylogenetic diversity of local communities to regional species pools, Webb showed that closely-related species were more likely to co-occur locally than expected when sampling randomly from the regional pool. These results are consistent with the “environmental filtering” hypothesis (Webb et al. 2002; Cavender-Bares et al. 2009; Keck and Kahlert 2019): particular environmental conditions select for species possessing specific traits, and because species with similar traits are likely to share a common evolutionary history, local phylogenetic diversity is lower than expected by chance (“phylogenetic under-dispersion”). The opposite pattern, phylogenetic over-dispersion, is typically interpreted as a signature of competitive exclusion, as articulated by Darwin. In this scenario, niche differences between species, rather than the matching of species and the environment, are the driving force behind community assembly (Webb et al. 2002; Cavender-Bares et al. 2009; Graham et al. 2009). The key assumption underlying both hypotheses is that the traits determining species’ interactions – and consequently coexistence – have a strong phylogenetic signal. However, this simple picture has been challenged in numerous studies, which have shown that the interpretation of phylogenetic under/over-dispersion can depend on the spatial scale, the number of clades included in the study, and various statistical issues inherent to observing only the endpoints of evolution and community assembly (Cavender-Bares, Keen, and Miles 2006; Mayfield and Levine 2010; Mazel et al. 2018; Davies 2021). These issues make it difficult to assess the relevance of phylogeny for understanding ecological processes and predicting ecological patterns. To overcome these obstacles, a combination of manipulative experiments and principled statistical approaches are called for.

Under the assumption that species’ interactions are tied to their phylogenetic relatedness, phylogeny can be further linked to ecosystem functioning. A basic consequence of this assumption is that phylogenetically diverse communities should harbor functionally diverse species, with less niche overlap and weaker competition, leading to increased ecosystem functioning (e.g., higher biomass, Venail et al. 2015; Davies et al. 2016). Thus, an important line of evidence for understanding the connection between phylogeny and community structure comes from studies of biodiversity and ecosystem functioning (BEF). Manipulative BEF experiments have shown not only that community biomass increases with species richness, but also that the phylogenetic diversity of experimental communities mediates their functioning (Maherali and Klironomos 2007; Connolly et al. 2011; Flynn et al. 2011; Cadotte 2013; Venail et al. 2015; Huang et al. 2020). Broadly, the results of BEF experiments are consistent with patterns found in agriculture and prairie grasslands (Jochum et al. 2020), making inferences from these experimental systems relevant for natural communities. However, careful statistical analyses of these general trends have led to conflicting conclusions regarding phylogeny and community functioning (Díaz and Cabido 2001; Petchey and Gaston 2006; Cadotte, Cardinale, and Oakley 2008; Cadotte 2013, 2015; Venail et al. 2015; Cardinale et al. 2015). Overlooked in earlier studies is the fact that species richness and phylogenetic diversity are positively correlated, and thus it is necessary to control for richness in evaluating the effect of phylogeny (Venail et al. 2015; Davies 2021). Additionally, many analyses have relied on relating community functioning to pairwise or averaged phylogenetic dissimilarity between species, ignoring the complex relationship between evolutionary tree topology and realized community structure (Mayfield and Levine 2010; Serván et al. 2020), as well as the processes that drive trait (co)evolution (Mazel et al. 2017; Swenson 2019). Despite considerable evidence that trait similarity, and therefore niche overlap, frequently increases with shared evolutionary history, simply correlating phylogenetic diversity with measured ecological properties fails to provide a clear-cut test of the connection between community function and phylogeny.

Previous attempts to incorporate phylogeny in the analysis of BEF data share two overaching features: a) they typically consider a single measurement for the function of each community (e.g., total biomass for each experimental plot, or the log-ratio of species abundances in multispecies communities to their abundances in monoculture), and b) they employ a statistical approach based on regressing these measurements against phylogenetic diversity or dissimilarity (Cadotte, Cardinale, and Oakley 2008; Flynn et al. 2011; Venail et al. 2015). Here we show that by abandoning these two features we can perform a strong and principled test for phylogenetic signal in these data, and we find that indeed phylogeny structures species’ interactions and thus patterns of coexistence and abundance.

The basic goal of this work is to build a framework in which, first, phylogenetic information is used to parameterize a model of interactions among a pool of *n* species; second, species’ interactions are used to predict coexistence and abundances for different assemblages that can be formed from the pool (Maynard, Miller, and Allesina 2020); and third, predictions are scored against experimental data from BEF experiments (Tilman et al. 2001; Van Ruijven and Berendse 2010; Cadotte 2013). To test whether phylogeny structures species’ interactions, we repeat this procedure many times, using both the “true” (molecular) phylogeny for a given community, as well as a large number of randomized trees. By comparing the resulting likelihoods, we assess whether the true phylogeny allows superior prediction of the ecological data, which would indicate that interactions are structured by this phylogenetic topology.

Our method departs considerably from previous approaches to testing for phylogenetic effects in the same datasets (Connolly et al. 2011, 2013; Cadotte 2013; Venail et al. 2015). Specifically, we make use of the biomasses recorded for individual species, rather than the total biomass for each plot, thereby gaining statistical power and resolution; we use the full tree topology in our statistical model, rather than summary statistics (e.g., phylogenetic diversity Cadotte, Cardinale, and Oakley (2008); Flynn et al. (2011)), allowing us to distinguish between similar trees; and, finally, we relate the tree structure directly to species’ interactions, and then to abundances via a model consistent with population dynamics, rather than using phylogenetic diversity as a covariate in a regression model.

Overall, our approach produces strong evidence for phylogenetic effects: the probability that a model parameterized with a random tree fits the empirical data better than a model based the actual phylogenetic tree is typically small, an outcome that is consistent across three different experimental settings, comprising a total of 19 distinct data sets. Additionally, we are able to identify which clades are most important for accurately predicting community function. Finally, we show that our framework can be extended in a number of ways and used to test other hypotheses about factors that structure ecological communities.

## Results

As detailed in the Methods section, we build a statistical model to test the effect of phylogeny on species interactions. We use this model to analyze data from three BEF experiments (Tilman et al. 2001; Van Ruijven and Berendse 2010; Cadotte 2013) in which plants selected from a pool of *n* species are grown together in a large number of different combinations. For these data sets, we have access to the recorded biomass of each species *i* across different plots in which it was grown along with other species. We use 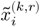 to denote the biomass of species *i* when growing in the assemblage *k* (corresponding to the set of extant species in the plot) and replicate *r* (as the same assemblage may be found in different plots). Based on the topology of the phylogenetic tree, we build a matrix of interactions for each data set. We then use this matrix of interactions to compute the expected biomass of species *i* in assemblage *k*, 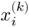, and by relating the expected and observed values via a likelihood function, we can find the maximum likelihood estimates of the free parameters in the model.

These operations are accomplished in three steps. First, the topology of the phylogenetic tree is combined with tunable parameters to produce a matrix of competitive interactions; the parameterization of the matrix is guided by models for consumer-resource dynamics (Methods and Supplementary Information; see also Serván et al. (2020)), and we consider four nested model formulations (model 1, the simplest, has 2*n* − 1 free parameters, models 2 and 3 have 3*n* − 2, and model 4 has 4*n* − 3). In all of these models, each branch *i* of the tree is associated with a parameter *λ_i_* ≥ 0, and larger values of *λ_i_* result in stronger competition between members of the clade subtended by branch *i*. Models 2 and 3 each include an additional vector of parameters (*α* and *γ*) allowing variation in consumption and uptake rates, respectively, and model 4 incorporates both parameter vectors (see Supplementary Information for details). Second, we use the matrix of interactions to compute the expected abundance of each species in every possible assemblage, following the approach of Maynard *et al*. (Maynard, Miller, and Allesina 2020, summarized in the Methods section). Thus, for each species in each assemblage, we obtain a prediction, influenced by both the structure of the tree and the tunable parameters. Finally, we specify a statistical model describing how experimental data are distributed around these predicted means (requiring an extra tunable parameter to control the shape of this distribution). This allows us to compute an overall likelihood for the free parameters given a specific tree, data set, and model formulation. We perform numerical optimization to search for the maximum likelihood estimates for the free parameters (Methods), ultimately allowing us to associate a (maximum) likelihood with each tree topology (and model formulation). Figure 1 represents these steps graphically, and the Methods section and Supplementary Information contain the mathematical and statistical details.

**Figure 1:**
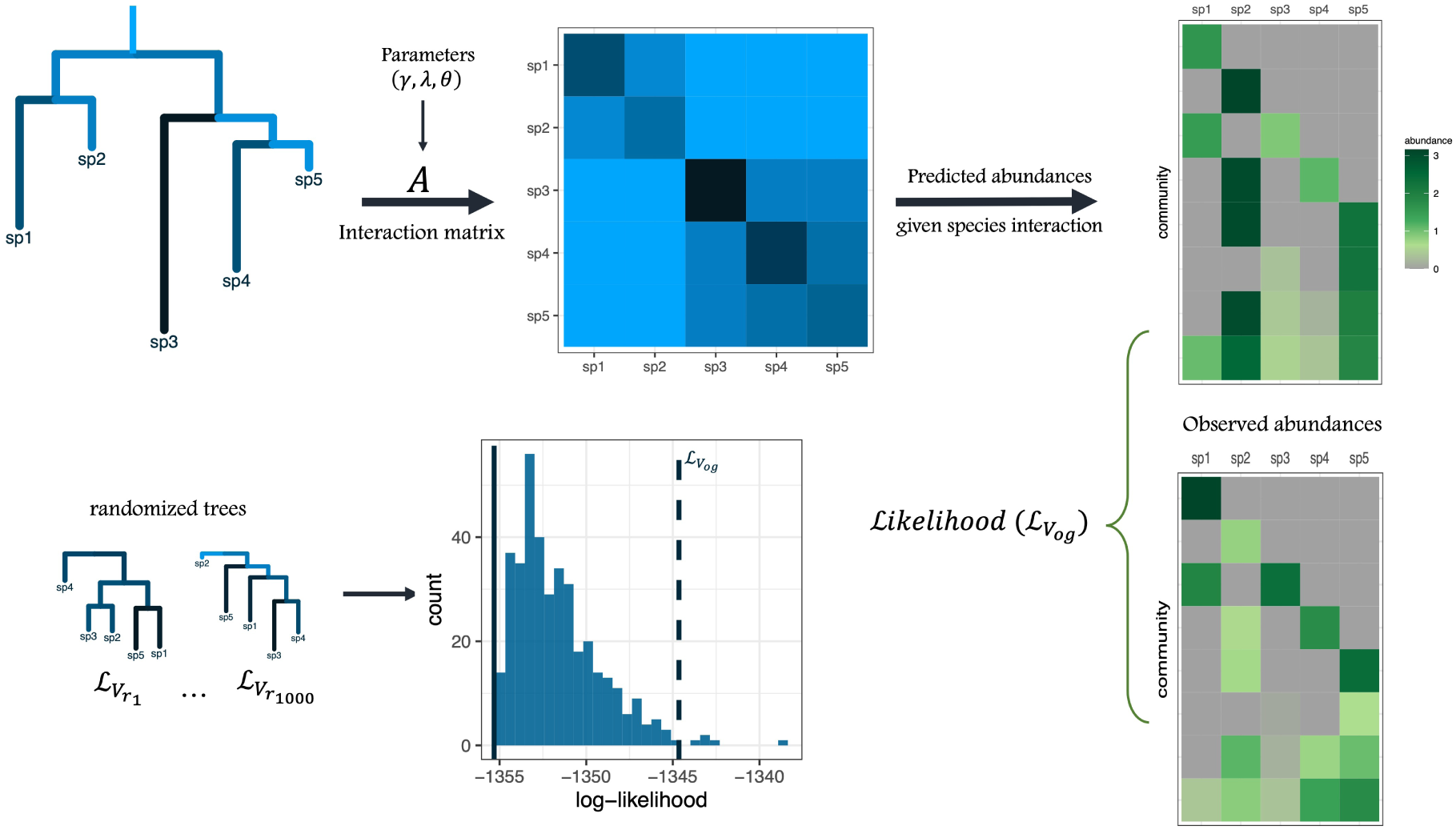
Diagram illustrating the steps of our approach. We combine a set of parameters and a phylogenetic tree to parameterize a matrix of interactions between all the species in a pool. This parameterization is inspired by consumer-resource models (Supplementary Information). Using the matrix of interactions, we predict whether a given assemblage of species will coexist, and at what abundances. These predicted abundances are thus determined by the structure of the phylogenetic tree as well as the tunable parameters. Abundances observed in experimental communities are assumed to follow a statistical distribution determined by these expectations, allowing us to compute the likelihood of the parameters given the tree, data and model. We maximize the likelihood of the parameters given the phylogenetic tree, obtaining a unique likelihood score for the tree topology, and then constrast it with model fits using randomized trees. By repeating this procedure many times, we build a distribution of likelihoods over the set of random trees. Finally, we compute a p-value which measures the probability of obtaining a better likelihood when using a random tree in place of the true phylogenetic tree.

Results for the data set published by Cadotte (2013) are shown in Figure 2 a, with one panel per model formulation. For this data set, even the simplest model, with a total of 16 free parameters (15 used to parameterize the interactions, one for the distribution), is able to recapitulate the observed abundances with good fidelity (8 species, measured in 27 distinct communities, for a total of 87 observations, including replicated assemblages). We find a correlation between the logarithm of the observed biomasses and the logarithm of the predicted biomasses close to 88%. When we consider more complex models, the log-likelihood improves marginally (from −348.37 to −342.55, see Table 1 for details; the correlation also grows to about 89%) at the cost of a substantial increase in the number of free parameters (30 parameters for model 4). In this case, and in all cases we consider, almost any metric for model selection (e.g., AIC, BIC) would favor the simplest model (model 1).

**Figure 2:**
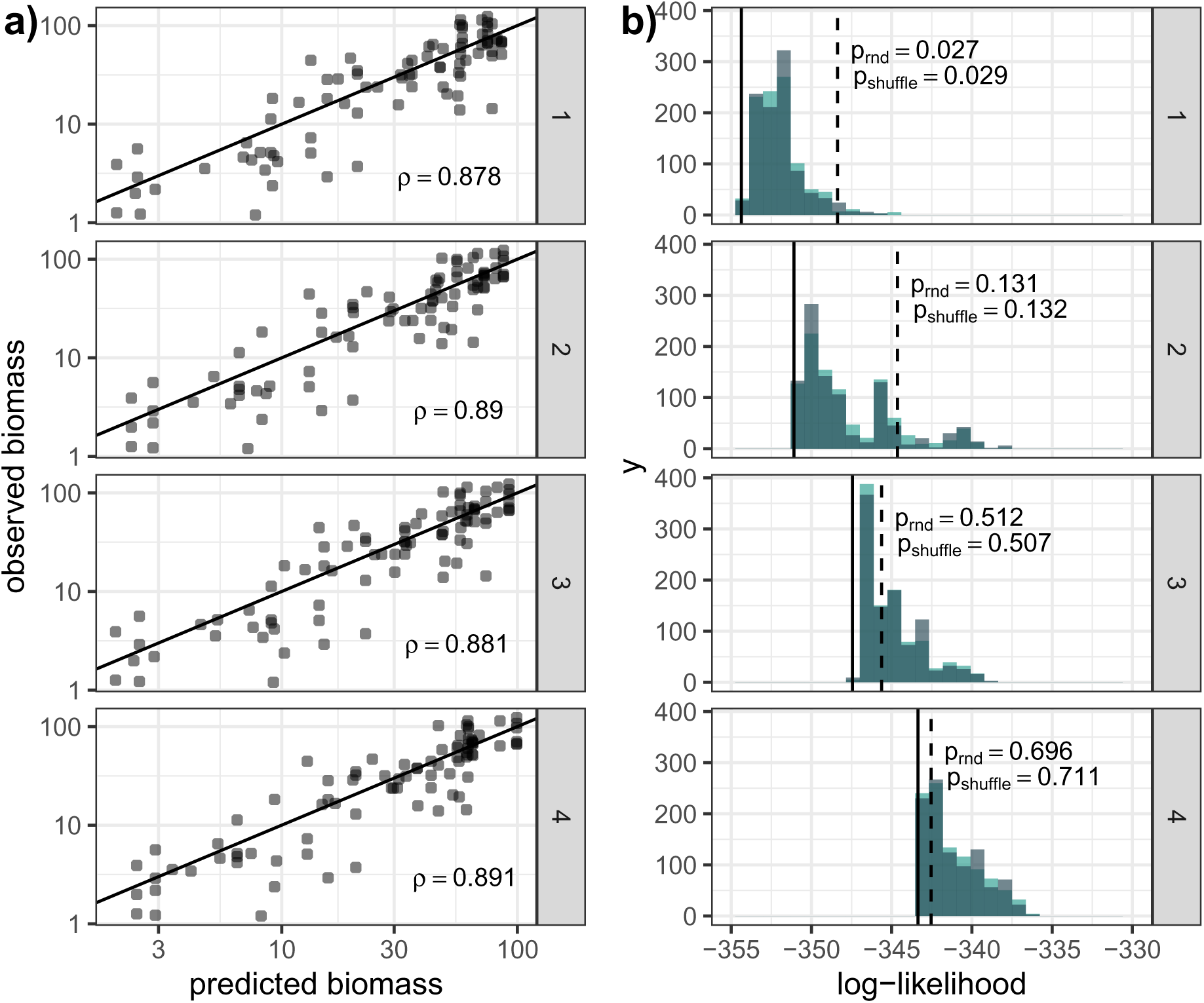
a) Predicted vs. observed biomasses for the data published by Cadotte (2013). Each point corresponds to the biomass measured for a given species in a given plot. The predicted value is necessarily the same across replicates for a given species/assemblage combination. The correlation between the logarithm of the predicted and observed biomasses is reported in each panel. b) Distribution of the likelihoods for the four models, fit using many randomized trees. The colors indicate whether the random trees were obtained from the Yule process (teal) or by shuffling the leaves of the original tree (grey). The vertical dashed line represents the likelihood using the original tree. The solid line represents the likelihood using a “star” tree, which provides a lower bound for the distribution (see text). Approximate p-values obtained for one thousand randomizations are reported in the panels.

**Table 1:**
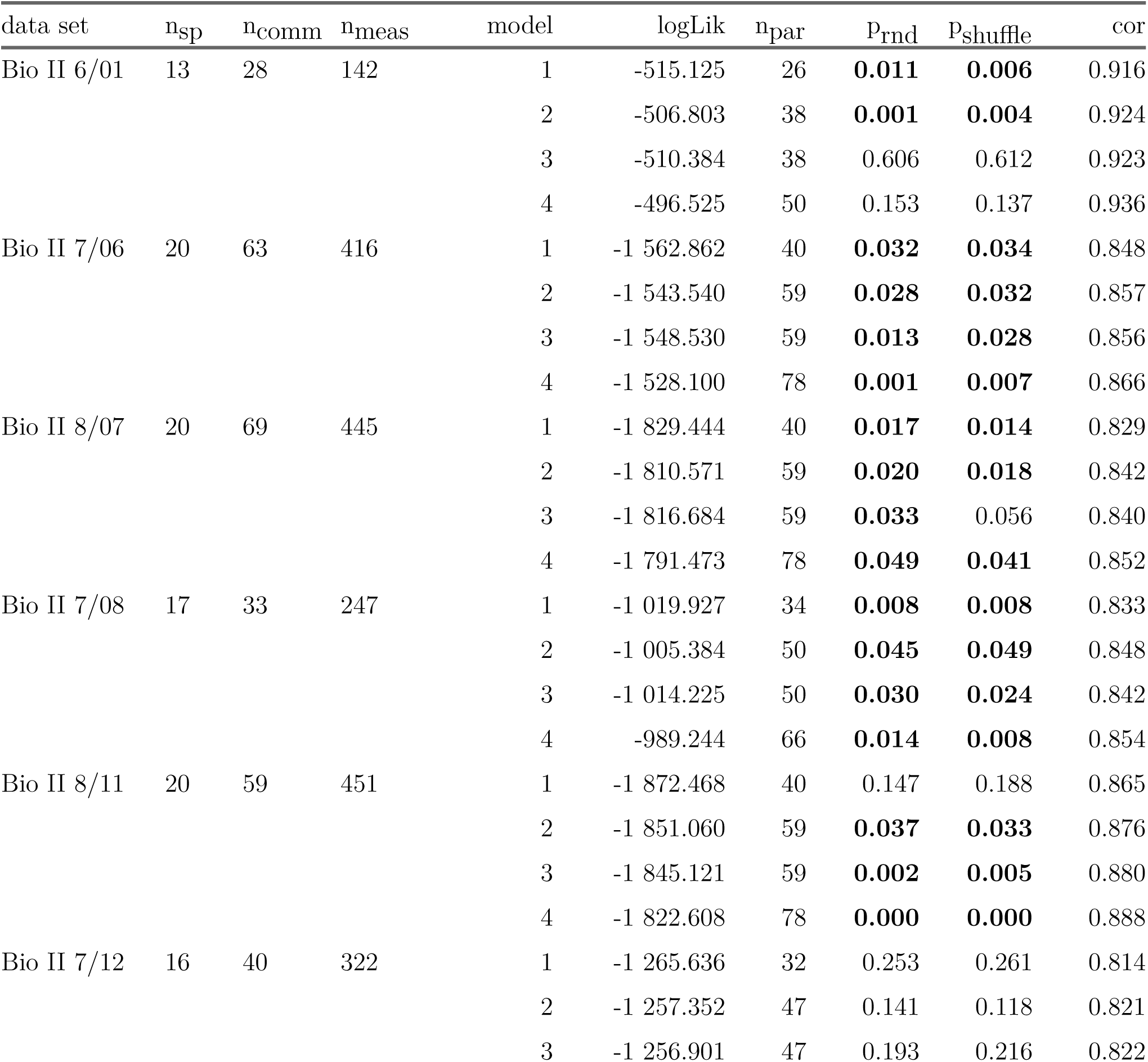
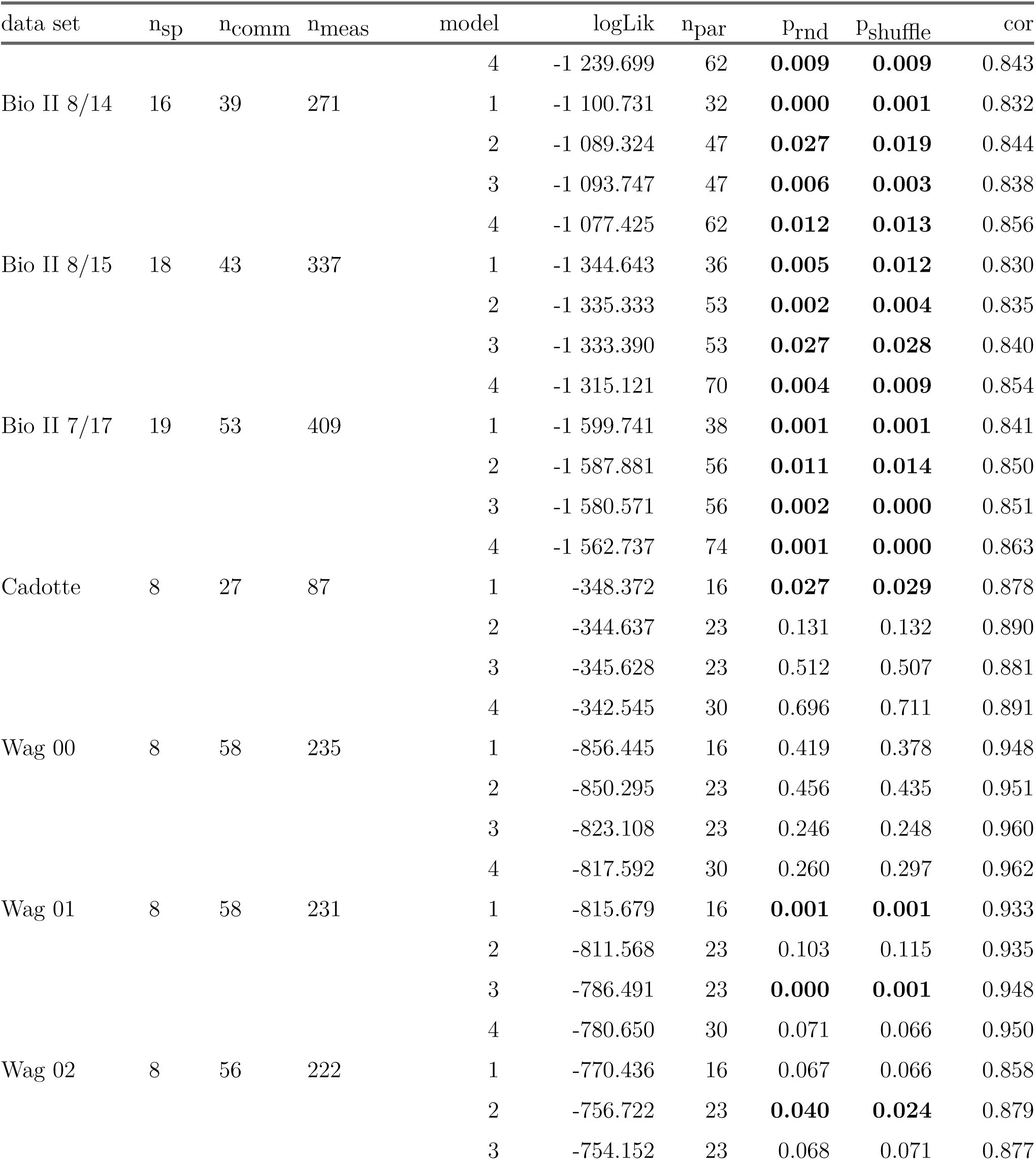
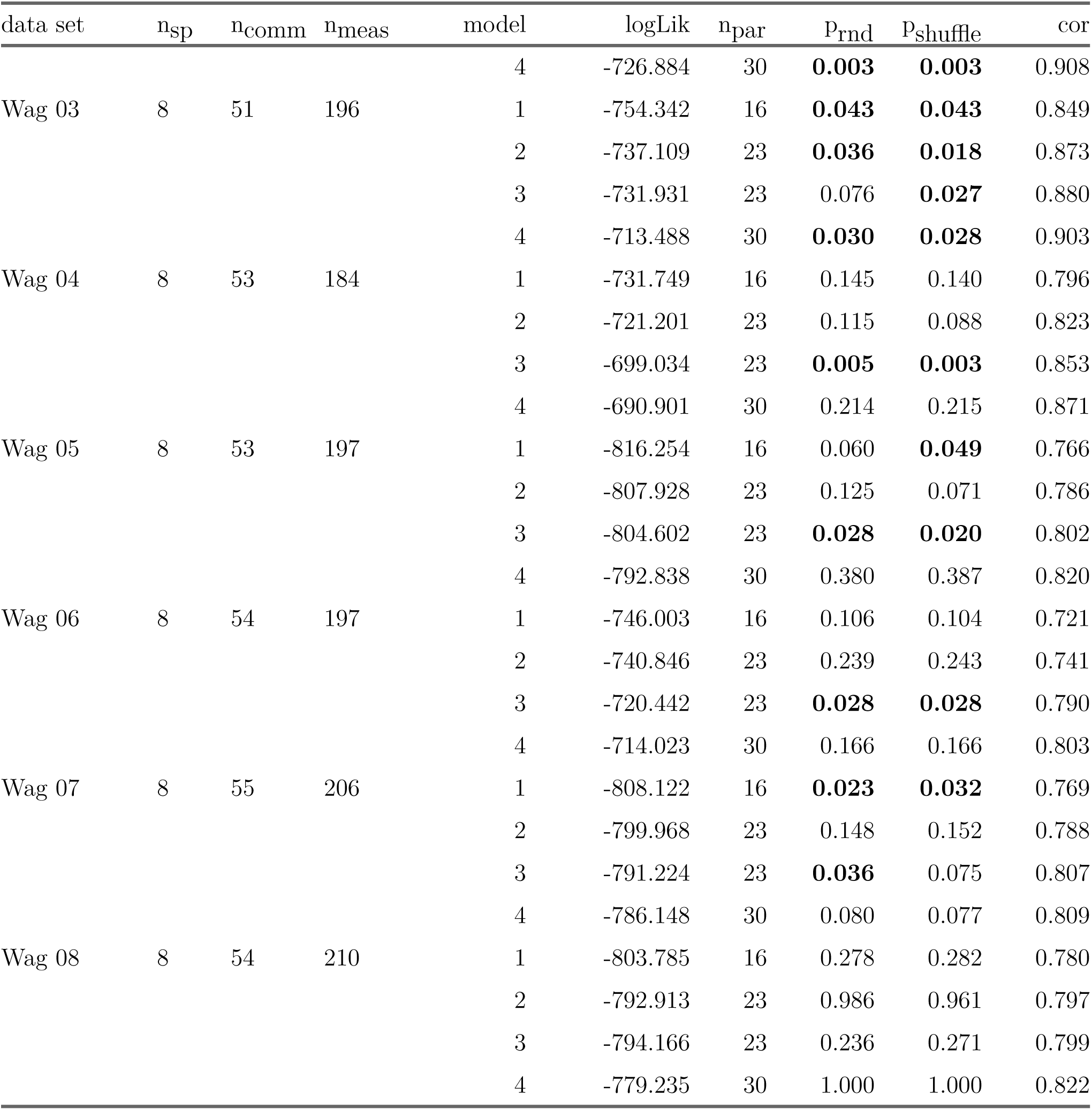
For each data set (Bio II, from Tilman et al. (2001); Cadotte, from Cadotte (2013); Wag, from Van Ruijven and Berendse (2010)), we report the number of species (nsp); the number of unique assemblages (ncomm); the total number of measurements (nmeas); the model used to fit the data; the corresponding log-likelihood (logLik); the total number of parameters (npar); the p-values for randomized trees either produced by the Yule model (prnd) or by shuffling the species’ identities (pshuffle); and the correlation between the logarithms of the observed and predicted biomasses (cor). P-values below 0.05 are bolded.

While these results indicate that models structured by phylogeny can describe the observed data quite well, they do not tell us if the particular tree topology is responsible for the quality of fit, or if other models of similar complexity (i.e., with a similar number of free parameters) would fit just as well. To answer this question, we compare the quality of fit, measured by its likelihood, against alternative models. First, as a natural reference point, we consider a “star tree”, in which all of the species descend directly from the root of the tree (Tucker et al. 2018). This configuration can be obtained from any tree topology by setting the parameters *λ_i_* corresponding to the internal branches to zero; therefore, the likelihood for the star tree provides a lower bound for the distribution of possible trees (continuous line in Figure 2 b) Next, we fit the model many times, each time using a different random tree to structure interactions. We consider two possible ways to build random trees: generated via the Yule model (teal in Figure 2 b) or randomized by shuffling species’ identities but maintaining the true tree topology (grey in Figure 2 b). Both approaches maintain the same number of free parameters, while the second approach maintains additional structure, such as the overall asymmetry of the tree. We build one thousand distinct random trees for each type of randomization and maximize the likelihood for each tree and model. From this ensemble of randomizations, we compute a p-value, quantifying the fraction of random trees that yield a better likelihood than the true phylogeny. Intuitively, 1 − *p* is the probability that using the true phylogenetic relationships for the community will improve the model fit.

For the Cadotte data set, Figure 2 shows that the star tree sets a lower bound for the likelihood, and that only a few a random trees perform better than the true tree (dashed black line) when using model 1. The p-values are approximately 0.029 and 0.028 for the shuffled and Yule trees, respectively. As we increase the number of free parameters, the flexiblity of the models increases, reducing the effect of the tree topology on the quality of fit. Accordingly, p-values increase for the more complex models (both close to 0.13 for model 2, and higher for models 3 and 4).

When we repeat this procedure for systems with a larger number of species, as in the Biodiversity II experiment (Tilman et al. 2001), we find even stronger evidence for phylogenetic effects. Additionally, when we analyze another system with eight species but more observations, from the Wageningen biodiversity experiment (Van Ruijven and Berendse 2010), we also find substantial evidence for phylogenetic effects. The p-values for every data set and model-randomization combination are reported in Table 1. In Figure 3 we summarize these results by plotting the distribution of p-values. This distribution is greatly enriched in low values. Out of a total of 152 tests (19 data sets, four models, two randomizations), we find that 86 p-values are ≤ 0.05, while we would expect less than 8 by chance alone. Another 14 are below 0.1, and 52 are above 0.1 (we would expect more than 136 by chance). This pattern is evident across the three experimental settings and across model formulations, even though we expect the more complex models to dilute phylogenetic signal. If we reduce the number of tests by pooling the results of the two randomizations (having found that the likelihood distributions for the two randomizations are very similar), we find 42 p-values ≤ 0.05 out of 76 tests (19 data sets, 4 models; 11 p-values ≤ 0.05 for model 1 and 3, 10 for models 2 and 4). Many p-values remain below this threshold when they are adjusted according to the Bonferroni (22 adjusted p-values ≤ 0.05) or Hommel (33 adjusted p-values ≤ 0.05) corrections. We conclude that these data sets bear a strong signature of phylogenetic structure.

**Figure 3:**
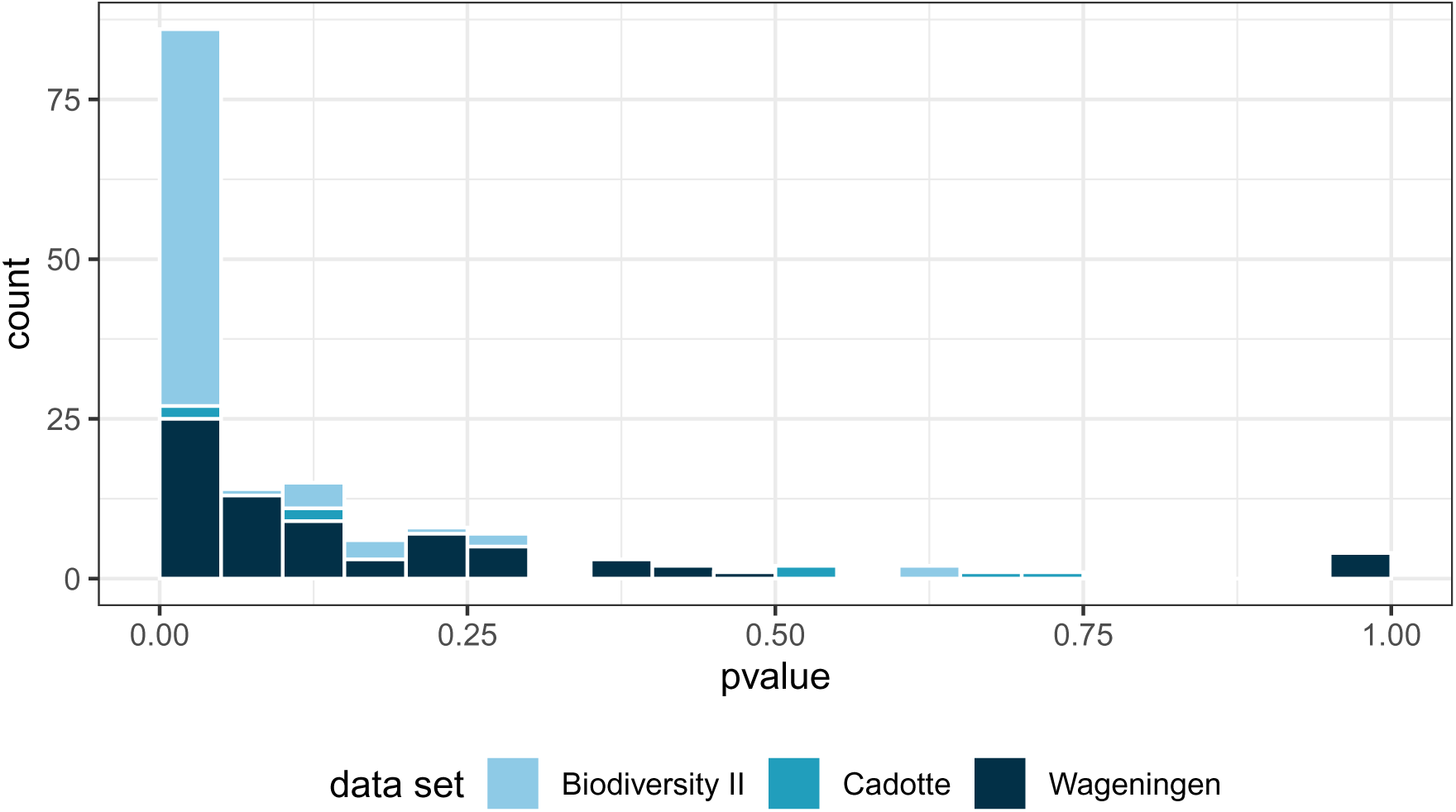
Pooled p-values obtained by testing for phylogenetic effects using four models, two randomizations, and nineteen data sets (152 p-values in total). We count the p-values falling in each bin of size 5% to build the histogram; colors indicate which of the three experiments the p-value corresponds to. The distribution of p-values is greatly enriched in low values, as expected when the null hypothesis of no phylogenetic effects (no difference between the true and random trees) can be rejected.

Our method also allows us to identify which features of the phylogenetic tree are the most important to capture the patterns in the empirical data. Take the simplest model, in which each branch of the tree (defining a clade) is associated with an increase in the strength of competition between the species in the corresponding clade. When species belonging to a certain clade do not experience increased competition with the other members of the clade, the maximum likelihood estimate of the corresponding branch length parameter will shrink to near zero. Figure 4 represents this feature visually: branches that do not strongly affect the pattern of interaction strengths are small, while those that help explain the data are associated with a positive value. As noted above, if the interactions had no structure, or structure not corresponding to the “input” tree toplogy, all internal branches would shrink to zero, giving rise to the star tree, in which only the branches associated with the root (encoding “mean-field” interspecific competition) and the tips (intraspecific competition) would have positive lengths. Thus, branches with positive values correspond to clades that help explain the structure of the empirical data.

**Figure 4:**
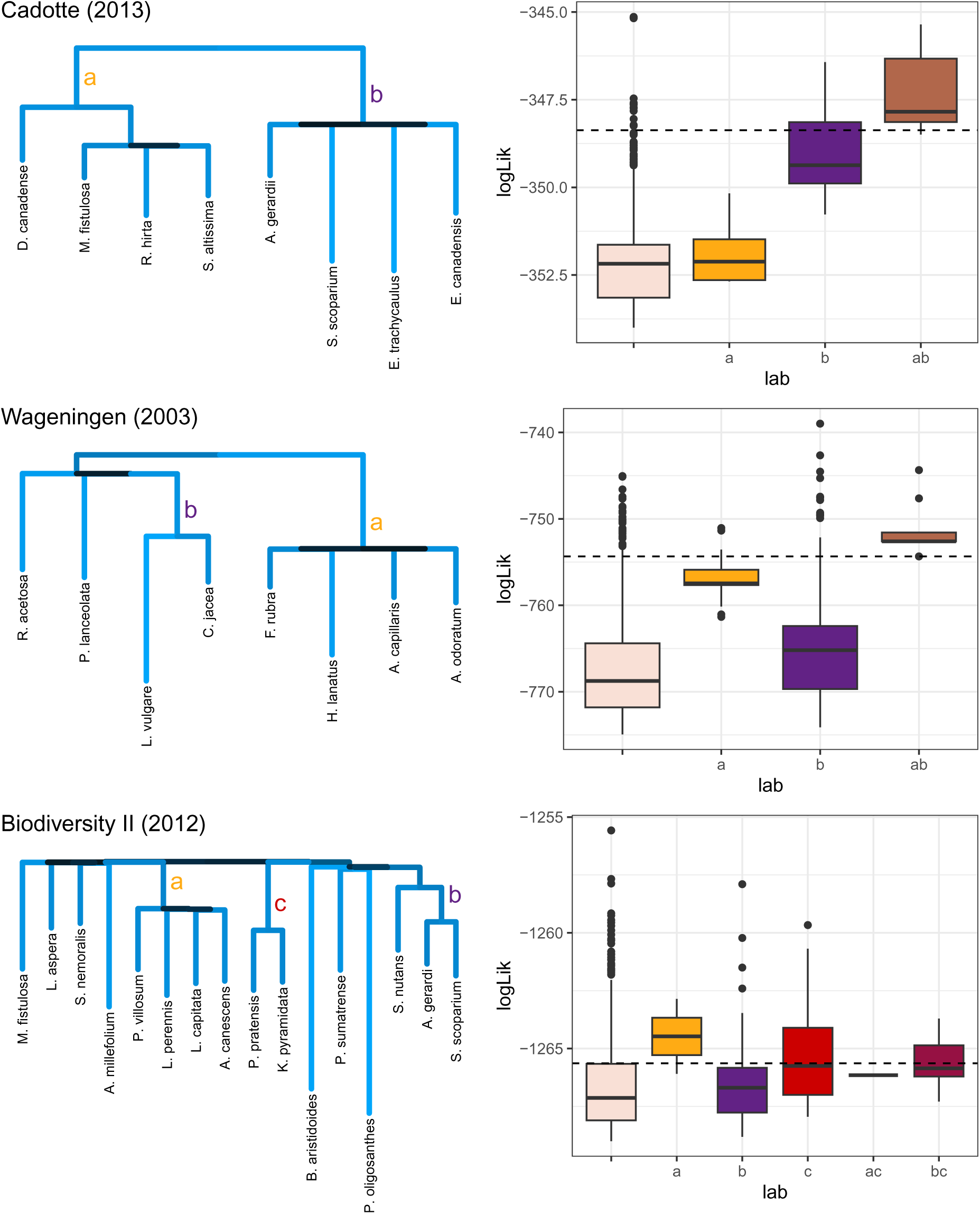
Maximum likelihood trees using model 1 for three selected data sets. The length and color of the branches correspond to increased strength of competition between the species subtended by the branch. Some branch lengths are close to zero, indicating that the species in these clades do not experience elevated competition with one another. For example, in the data from Cadotte (2013) the model does not support an increase in the strength of competition between *R. hirta* and *S. altissima*, compared to their interactions with *M. fistulosa*, despite the additional evolutionary history shared by these two species. As a result, this clade collapses into a polytomy with three species. Box-plots show the distribution of log-likelihoods for randomizations that retain the clades highlighted in the phylogenies (indicated by letters; boxes without labels correspond to randomizations where no labeled clades were retained). The dashed line indicates the

For the data from Cadotte (2013), the likelihood maximization singles out the clade of grasses (Poaceae, clade *b* in the phylogenetic tree in Figure 4), and separates the only legume (*D. canadense*) from the rest of the species (clade *a* in the figure). Likewise, the analysis of the data by Van Ruijven and Berendse (2010) (year 2003 shown) highlights the competition between the four grasses (Poaceae, clade *a* in Figure 4), as well as the two members of Asteraceae (clade *b*). We can confirm this interpretation by stratifying the random trees according to the presence of these highlighted clades. For example, when fitting the data from Cadotte (2013), we find that those trees possessing clade *a* perform slightly better than other random trees, those that group the species that are not grasses or legumes (clade *b*) perform much better, and those retainng both *a* and *b* (as in the true tree) perform best than all. In fact, this last set of trees performs as well as or better than the true tree (likelihood marked by a dashed line), indicating that these two partitions are the salient components of the phylogenetic structure. Similar results are found for the Wageningen experiments trees clustering the grasses or Asteraceae perform better than those that do not, and trees clustering both clades correctly yield a much higher likelihood. For more speciose trees, we also see evidence that a few clades contribute strongly to the quality of fit, although in this case sampling enough random trees to find all possible combinations of these key clades would require an astronomical number of randomizations. For example, in the Biodiversity II experiment (year 2012 data) we see support for increased competition between the legumes (clade *a*), and all random trees clustering the legumes together yield high likelihood; the grasses are split into several distinct clades, with strong support for increased competition between two C4 grasses (clade *b*), and two C3 grasses (clade *c*). Again, randomized trees conserving these clades perform better on average than those that do not.

We have focused here on interpreting model parameters for the simplest model (model 1), but similar conclusions apply to the more complex models, in which parameters can be interpreted in the same way. All four models point in the same direction of these data sets being phylogenetically structured, as evidenced by the overall results presented in Table 1 and the distribution of p-values. Even as we add more flexibility to the model through the incorporation of species-specific parameters related to consumption and uptake rates, the presence of phylogenetic signal remains quite robust (see Supplementary Information for the fit of models 2-4).

## Discussion

In this study, we leveraged the statistical framework of Maynard, Miller, and Allesina (2020) to fit species interaction models where interaction strengths are systematically tied to evolutionary history, summarized by phylogeny. By fitting models using both the true phylogenetic tree for a set of species as well as a randomized ensemble, we have devised a simple and direct test to determine whether phylogeny structures species’ interactions, and thus patterns of coexistence and abundance in empirical data. Analyzing nineteen data sets from three different biodiversity-ecosystem functioning experiments, we found substantial support for phylogenetic structure in species’ interactions. In this section, we briefly discuss the advantages and limitations of this approach, as well as possible extensions.

### Advantages of the approach

Our methodology uses biomasses recorded for all species in a plot, as opposed to the total biomass in the plot. While previous studies (Tilman et al. 2001; Connolly et al. 2011; Cadotte 2013; Huang et al. 2020) focused only on total biomass in order to test BEF hypotheses, examining these data at the level of species yields a larger number of measurements, giving greater statistical power to detect phylogenetic structure, while also reducing the risk of model overfitting. Because BEF and other community data sets often contain species-level biomasses (or abundances), our approach makes fuller use of the painstaking empirical work that went into building these large data sets.

This approach is grounded in a simple framework to infer interactions and predict biomasses from community “endpoints” — a framework that is gaining traction in the ecological literature (Xiao et al. 2017; Fort 2018; Maynard, Miller, and Allesina 2020; Ansari et al. 2021; Skwara et al. 2023). One of the main advantages of this framework is that it makes predictions that are compatible with models of population dynamics: if we were to simulate communities using the fitted interaction strengths as parameters of the Generalized Lotka-Volterra model, we would find the same predicted abundances as equilibria of the dynamical system. This is in contrast with models based on a regression framework, which are inherently incompatible with population dynamics (see Supplementary Information). To include phylogeny in this framework, we took a principled approach, formalizing the intuitive expectation (already clear to Darwin) that shared ancestry translates into stronger competition. This translation between phylogenetic structure and interaction strengths was guided by consumer-resource models (Methods, Supplementary Information), permitting a clear ecological interpretation of the parameters.

This basic approach is quite modular and flexible; for example, here we used four related models—each structured by phylogeny but including different levels of additional complexity—and employed two different randomizations to test for phylogenetic effects. Other model formulations or randomizations could be used to test for phylogenetic structure within the same framework, and one could additionally alter the assumptions of the statistical model (e.g., we assumed that empirical values were Gamma distributed, but one could substitute any other distribution with positive support) or otherwise tailor the pipeline for a specific data set and ecological question.

The models we have employed use the full topology of the phylogenetic tree to structure interactions. This means that, in principle, we can use likelihoods to discriminate between trees with similar coarse structure (e.g. sum of branch lengths, a commonly used summary statistic). We believe this is a substantial step forward compared to approaches that rely on a single metric for trees induced by each assemblage, as the detailed structure of a phylogeny might mediate important differences in community patterns. Additionally, we depart from the simplistic assumption that interaction strength should increase linearly (or according to another simple function) with shared ancestry. Instead, we associate a free parameter with each branch, thereby allowing some branches to take on greater importance, and other to shrink to zero, guided by the data. Essentially, we assume that interaction strengths do increase with shared ancestry, but make no other assumption about the form of this relationship. This allows us to capture phylogenetic effects even if the underlying processes of trait evolution are highly variable in time and across the phylogeny (Hansen and Martins 1996; Mazel et al. 2017; Tucker et al. 2018).

This model flexibility also allows us to identify which branches play a key role in structuring interactions. Previous efforts to move beyond straightforward phylogenetic distances used network tools to identify important branches in phylogenies for experimental communities (Davies et al. 2016). Analyzing the Biodiversity II data, as we did here, Davies et al. (2016) used this approach to detect an important effect of legumes on community function. In agreement with their conclusions, we also find that the clade including the legumes plays a central role in structuring these experimental communities (clade *a* in Biodiversity II, Figure 4). For the Wageningen experiment (Ruijven and Berendse 2005; Van Ruijven and Berendse 2010), an examination of inferred branch lengths reveals temporal patterns. This experiment showed increasing complementary effects over time, suggesting differentiation of resource use or other traits, or increasing facilitation. These adaptive processes at the community level take time and were not detected in the first year of the Wageningen experiment; however, in the experiment’s subsequent years there was a strong signal of complementary effects that increased in time and with species richness (Van Ruijven and Berendse 2010). Our approach recovers these temporal effects: in the first years of the experiment (2000, 2001 in Supplementary Information and 2003 in Figure 4) we find evidence for a strong phylogenetic signal in the clades containing the grass species and most of the dicots. From 2004 on (Supplementary Information), the fit of our model shows that the branch lengths of the dicot clade are reduced, with these species mostly collapsing into a polytomy and only the grasses retaining a strong phylogenetic signal. The decrease in the fitted branch lengths over time in the experimental data suggests reduced competition, which is consistent with the reported increase in complementary effects (Ruijven and Berendse 2005; Van Ruijven and Berendse 2010) and points to the phylogenetic origins of these effects.

### Limitations of the approach

There are important limitations, both practical and theoretical, to this methodology. Practically, our approach is computationally very intensive, especially due to the large number of randomizations needed to obtain accurate p-values. This issue is exacerbated for data sets with many species because, for each parameterization, it is necessary to invert an interaction sub-matrix for each observed assemblage. The computational cost of this operation scales approximately with the cube of the size of the matrix (number of species). This problem can be alleviated by fitting randomizations in parallel with computer clusters, but it remains a key limitation (for example, the calculations presented here required approximately 5 years of computing time). The computational challenge is made worse by the fact that we expect the likelihood surface of the parameter space to contain many local maxima. Thus, finding the global maximum likelihood is challenging, and at a minimum requires initializing the numerical search from several initial conditions.

On the theoretical side, we have considered four nested models, capturing progressively more ecological detail, and moving from the simpler to the more complex formulations adds approximately parameters at each step. Models with more parameters are more prone to over-fitting (i.e., fitting the noise as well as the signal in the data), and in fact we find that the contribution of the phylogenetic tree can sometimes be swamped by the large number of parameters in the most complex models (e.g., Figure 2). As we have seen in the Results section, some—and sometimes many—of the model parameters shrink to zero through likelihood maximization, and could therefore be removed without affecting the goodness of fit. This suggests an approach to reduce the number of parameters, by analyzing degenerate trees in which some branch have zero length (i.e., non-binary phylogenetic trees), or coarse-grained trees (e.g., based on higher taxonomic levels).

### Beyond phylogeny

Interestingly, from an ecological perspective, the collapse of many fitted branch lengths suggests that only parts of the phylogeny—key clades, possibly corresponding to key functional traits—matter for species’ interactions. Thus, we might hope to test for other, perhaps simpler, kinds of structure. This is possible through a straightforward extension of our approach. In this framework, the topology of a tree is encoded by binary matrix *V* (see Methods), in which rows correspond to ancestral species, columns to extant species, and *V_ij_* = 1 whenever the row is an ancestor of the column (and zero otherwise). The same approach could be extended to any binary matrix *V* (provided the resulting matrix of interactions and all its sub-matrices are invertible). For example, we could postulate models in which there are *k* generalized “groups” of species (with or without nested or overlapping membership). Then the rows of *V* would be groups, and *V_ij_* = 1 whenever species *j* belongs to group *i*. Groups might correspond to functional traits: in this case the rows of *V* are traits and *V_ij_* = 1 whenever species *j* possesses trait *i*. These simple yet powerful extensions allow our basic pipeline to be used to test for community structure well beyond phylogenetic signal. While we find evidence for phylogenetic structure in the data sets considered here, the strong assumption that phylogeny serves as a proxy for ecological similarity might not always hold in natural communities, or might be subject to distortion. For example, trait similarity might be a consequence of convergent evolution, as opposed to shared evolutionary history (Cavender-Bares, Keen, and Miles 2006; Losos 2008). Further, traits determining species interactions—and hence influencing ecosystem functioning—may be under strong selection, and consequently evolutionary history as inferred from neutral markers might not reflect functional trait similarity (J. P. Wright et al. 2006; Davies et al. 2016). In such cases, our approach makes it possible to explicitly test the role of traits known to be under selection or known to influence species interactions and coexistence (Fry, Power, and Manning 2014). In fact, setting aside the computational challenge of fitting a large number of trees, one could in principle attempt to find the maximum likelihood tree topology or group structure and effectively “reverse-engineer”” the traits or other factors that contribute most to community structure and functioning.

Importantly, regardless of the chosen structure, the same randomization scheme can be used to test whether the traits or groups of interest are informative about ecological outcomes. Using an appropriately chosen randomized ensemble, one can always compare a matrix of interest against randomizations to compute a p-value, which quantifies the value of using the chosen structure to constrain interactions, as compared to models that are equally complex, but biologically “uninformed”.

### Beyond competition

So far, we have not relaxed the assupmtion that species’ interactions are strictly competitive. Even when considering species at same trophic level, the view of competition as the only force structuring communities might be too limited (A. J. Wright et al. 2017; Frainer et al. 2018). One of the striking patterns observed in BEF experiments is “overyelding”, in which a species achieves a higher biomass in mixed communities than it does in monoculture (Connolly et al. 2013). This pattern has been associated with complementary effects, but might also reflect facilitation (A. J. Wright et al. 2017). Current approaches make it challenging to disentangle these two mechanisms, but the flexibility of our framework might also help resolve the role of facilitation in structuring these ecological communities.

There are at least two ways our framework could be extended to allow for other types of interactions besides competition. First, by letting the clade-specific interactions (*λ_i_*) take any sign, positive or negative, we can model facilitation or mutualism together with competition. Second, we can let the auxilary parameters modulating phylogenetic effects (*γ*, *θ*; see Methods) take arbitrary values, thereby allowing asymmetric interactions, such as predation or paratism. None of these changes increases the number of free parameters, but should always result in a better fit to the data by removing constraints on the parameters.

Naturally, this flexibility comes with costs, including increased risk of overfitting and more complex interpretation of the model parameters. To avoid these issues and retain maximal interpretability, we have concentrated on less flexible models with competition only, although these extensions represent a natural area for future inquiry.

## Conclusions

We have introduced a principled, robust statistical approach to detect phylogenetic effects on species interactions. Given the topology of a tree, we find a maximum likelihood model by optimizing over a set of free parameters. Using data from three different biodiversity-ecosystem functioning experiments, we have found that the models structured by phylogeny provide a good fit to observations, and provide strong support for the presence of phylogenetic effects. Our approach departs from existing regression-based frameworks, making it compatible with population dynamic models; moreover, it more fully exploits all of the information in the published data, both in terms of measurements (using biomasses recorded at the level of species instead of at the level of plots), and in terms of phylogeny (using the full tree structure instead of summary statistics). Even though it is computationally intensive, our approach can easily be extended to account for functional groups, explicit traits or other types of interactions beyond competition.

## Methods

### Modeling framework

As outlined in the main text, our statistical framework is composed of three fundamental steps. Each of the steps could be carried out in a number of ways.

#### a) From tree topology to interactions

We represent the topology of a binary phylogenetic tree with *n* leaves (i.e., the species in the pool) as a binary matrix *V* of size (2*n* − 1) × *n*. The rows of the matrix represent either the extant species or their ancestors (including the common ancestor to all species, i.e., the root of the tree), while the columns represent the extant species. A coefficient *V_ij_* = 1 if *i* is an ancestor of *j* (or itself *j*), and is zero otherwise. This matrix is called the “basis matrix” of the tree in the literature (Bravo et al. 2009), and is closely related to the correlation structure induced by shared evolutionary history (Hansen and Martins 1996): given a vector of 2*n* − 1 branch lengths *α*, then *V^T^D*(*α*)*V* is the variance-covariance matrix of the tree.

Inspired by classic consumer-resource models (Supplementary Information), we parameterize the matrix of interactions between the species as:

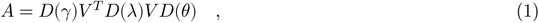

where *γ* and *θ* are positive vectors of length *n* with entries summing to one (thus representing *n* − 1 free parameters each), *λ* is a positive vector of length 2*n* − 1, *D*(*ζ*) is a diagonal matrix with the elements of *ζ* on the diagonal, and *V^T^* denotes the transpose of matrix *V*. We therefore have 4*n* − 3 free parameters in total. We also consider simplified models in which either *D*(*γ*), *D*(*θ*), or both are taken to be 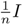, i.e., a scaled version of the identity matrix, reducing the number of parameters to 3*n* − 2, 3*n* − 2 and 2*n* − 1, respectively. Details on these parameterizations and their derivation from consumer-resource models are reported in Supplementary Information.

Provided with a tree topology describing the evolutionary history of the *n* species in the pool (defining th associated matrix *V*), and empirical data in which different sets of species taken from the pool are grown together, we aim to find the maximum likelihood estimates of the parameters *γ*, *λ*, and *θ*. To achieve this goal, we first relate interactions to predicted abundances, and then predicted abundances to experimental data.

#### b) From interactions to predictions

We want to relate species interactions, encoded in the phylogenetically-structured matrix *A* (Eq. 1) with the abundances species would attain in different assemblages. Take *k* to be the set of species in a given assemblage. We assume that the abundance of species *i* in assemblage *k*, denoted 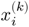, is linearly related to the abundance of the other species in the assemblage, with each effect mediated by the relevant pairwise interaction strength:

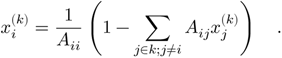

Thus, species *i* grown alone would reach 1/*A_ii_* (its carrying capacity), while growth together with *j* would yield a lower value whenever *A*_ij_ > 0, i.e., when species *i* and *j* are competitors.

Writing *A*^(*k*)^ for the sub-matrix of *A* obtained by retaining only the columns and rows of species in assemblage *k*, we can write this system of equations in more compact form as:

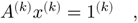

where 1^(*k*)^ is a vector of ones with as many elements as there are species in *k*. Thus, provided with a matrix *A* we can compute the value of *x*^(*k*)^, for any *k*:

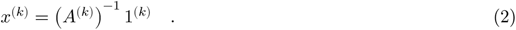

There are two possible outcomes: a) all the elements of *x*^(*k*)^ are positive, which we interpret as predicted coexistence of the species in assemblage *k*, with expected abundances specified by the vector *x*^(*k*)^; b) one or more of the elements are negative, which we interpret as the impossibility of coexistence for this set of species. Versions of this basic framework have been proposed numerous times in the ecological literature (Xiao et al. 2017; Fort 2018; Maynard, Miller, and Allesina 2020; Ansari et al. 2021), and here we closely follow the approach of Maynard *et al*. (Maynard, Miller, and Allesina 2020; Skwara et al. 2023).

Provided with a matrix *A* defined in Eq. 1, we can thus predict the expected coexistence and abundance of species in all possible assemblages using Eq. 2. Finally, we can relate these predictions to the empirical data.

#### c) From predictions to likelihoods

Each data set comprises a set of measurements 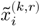, specifying the biomass of species *i* when growin in assemblage *k* (i.e., defined by the extant species in the plot) and replicate *r*. The data sets contain a large number of assemblages, possibly with replicated experiments (i.e., distinct plots in which the same assemblage is observed). We take these measurements to be noisy estimates of the “true”, expected biomass of species *i* in assemblage *k*:

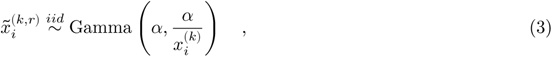

where we assume that the observations have been sampled independently from a Gamma distribution with parameters *α* (scale) and 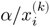 (rate), such that the expected value is 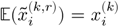 and the variance is 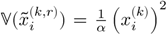. The parameter *α* is assumed to be specific to the data set. Thus, for a given data set all measurements are taken from distributions with a constant coefficient of variation, 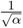. Whenever *α* > 1, the mode of the distribution is larger than zero, and the shape of the distribution displays a central tendency.

The choice of this particular Gamma distribution is convenient (e.g., it has support on the positive real line, a constant coefficient of variation, and adds a single free parameter); alternatively, one could assume a Log-Normal distribution, or any other appropriate alternative. We do not expect the choice of a particular distribution to affect our qualitative conclusions.

### Numerical search for maximum likelihood estimates

In summary, provided with the phylogenetic tree for the species pool, the parameters of the model (the vectors *γ*, *θ* and *λ*, as well as the parameter *α* controlling the shape of the distribution), and the empirical data, comprising empirical measurements of the biomass of the species across different (possibly replicated) assemblages, we can: a) use the parameters and the tree to build a matrix of interactions (Eq. 1), b) compute the predicted abundances for all assemblages (Eq. 2), and c) compute a likelihood by multiplying the probability of observing each measurement under the fitted model Eq. 3.

This bridge from parameters to likelihoods allows us to search numerically for the maximum likelihood estimates of the parameters. Briefly, a) we initialize the parameters such that *A* is the sum of the identity matrix and a random matrix with small norm (practically, ensuring coexistence in all possible assemblages); b) search for parameters that increase the likelihood via a combination of simulated annealing, followed by hill-climbing (where slightly alter parameters and accept the move if likelihood is increasing), and numerical optimization via the Nelder-Mead and BFGS algorithms provided by the *optim* function in *R* (R Core Team 2021); c) step in b) is repeated 500 times, each time improving the likelihood until (presumptive) convergence to the maximum likelihood estimates. Parameterizations that predict a lack of coexistence for assemblages that were observed to coexist experimentally are discarded. Parameterizations that yield a likelihood lower than found for the corresponding star tree are also discarded. Finding optimal parameters is computationally very difficult (there can be many local optima in the parameter space), and very expensive (because each prediction requires the inversion of a matrix, following Eq. 2). We implemented the approach using a combination of *Stan* (Stan Development Team 2023) and *R* (R Core Team 2021) code, started each parameter search five times with a random initialization, and retained the best likelihood for each data set/model combination. The code, raw and organized data, and results are available at github.com/StefanoAllesina/lemos-costa-2023.

### Testing for phylogenetic signal

In the preceding sections, we showed that, given empirical data and a matrix *V* corresponding to a phylogenetic tree, we can find the maximum likelihood estimates of the parameters in Eq. 1, and thus the maximum likelihood associated with the tree, 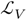. To test whether phylogeny can explain species interactions, and consequently the observed abundances in different assemblages, we repeat the same fitting procedure for a random tree *V*′, yielding 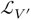. By repeating this procedure many times, we can estimate the probability that a random tree yields a higher likelihood than the “true” tree associated with *V*. As such we take 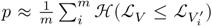, where *m* is the number of random trees, 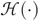 the Heaviside step function and 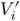 is the matrix associated with the *i*^th^ random tree.

Finally, we need to define what we mean by a random tree. We consider two possibilities: a) a random tree with the same number of leaves constructed according to the Yule model, a classic speciation model in which every branch has the same probability of speciating at every time; b) a tree with the same topology as the original, but where we shuffle species’ identities. Therefore, for each data set we present randomizations for four nested models (all variations on Eq. 1) and two distinct ways of constructing the random trees. For each randomization method, we sample tree topologies until we have found one thousand distinct trees; whenever duplicate trees are encountered in this process, we fit both and retain the best likelihood.

### Data and data organization

The methods of Maynard, Miller, and Allesina (2020) were devised for the analysis of experiments carried out in laboratory conditions with a small species pool, for which a substantial proportion of all the possible combinations of species were observed. Biodiversity-Ecosystem Functioning experiments, on the other hand, are often conducted in experimental plots, using a pool of a dozen or more species. Because the number of possible assemblages one can form with species is 2*^n^*, and only dozens or hundreds of experimental plots are typically feasible, the number of sampled assemblages in these experiments is necessarily small compared to the theoretical limit (e.g. 2^1^0 is already more than a thousand distinct assemblages for *n* = 10). We therefore attempted to select data sets in which a) only species that appear in multiple plots are considered to be part of the pool; b) unidentified biomass or non-target species do not dominate the plots; c) we have sufficient replication or diversity of assemblages to confidently fit the parameters of the model.

To this end, we consider data collected as part of the Biodiversity II experiment (Tilman et al. 2001, data from 2001 to 2018, inclusive, downloaded from the Cedar Creek website), the Wageningen diversity experiment (Van Ruijven and Berendse 2010, provided by Dr. Jasper van Ruijven), and an experiment carried out at the Toronto’s Koffler Scientific Reserve by Marc W. Cadotte (data accompanying Cadotte 2013).

The data from Cadotte (2013) contains plots in which a subset of 14 species were planted. Some did not grow and went extinct during the experiment. We consider a species that did not reach 2.5% of its maximum observed biomass to be extinct in the plot. We then considered species that appeared in at least 5 distinct assemblages; if a plot contained species that did not meet this requirement, the whole plot was discarded; the procedure was then repeated until we obtained a subset of the original data in which all species appeared in at least 5 distinct assemblages. This left us with 8 species, forming 27 distinct assemblages, for a total of 87 measurements (including replicated assemblages). Code to implement this procedure, along with the raw and elaborated data are accompanying this work.

The same general procedure was applied to the data from Van Ruijven and Berendse (2010), with the difference that we required each species to be present in 5 distinct assemblages. After the selection procedure, all eight species were retained, and we obtained nine separate data sets, one for each yearly sampling from 2001 to 2009, inclusive.

The data from Tilman et al. (2001) are the most difficult to process, because aside from the original target pool of eighteen species, many other species are found in the plots, along with a substantial amount of litter or unidentified biomass. We therefore included in the “species pool” a larger number of species, excluded plots in which the total biomass for species not in the pool surpassed 10% of the total biomass, and then applied the procedure used for the other data sets. In this way, we were able to extract nine data sets (for distinct years) containing between 13 and 20 species and a substantial number of assemblages.

Summary statistics for each data set, including number of species, number of unique assemblages and number of measurements are reported in Table 1.

To obtain a phylogeny for each species pool, we normalized species names using the World Flora Database (http://www.worldfloraonline.org/) and then constructed the phylogenetic tree associated with each set of species using the *V.PhyloMaker* package in *R* (Jin and Qian 2019).

## Ackowledgments

We thank C. Serván, P. Lechón-Alonso, M. Chakraverti-Wuerthwein, S. Kundu, and M. Loschi for fruitful comments and discussion, and to J. Van Ruijven for sharing the data from the Wageningen experiment. We acknowledge the University of Chicago’s Research Computing Center for their support of this work. This work was supported by the National Science Foundation (DEB #2022742 to SA).

## Supplementary Information

### Derivation of the model from consumer-resource dynamics

**Figure 5:**
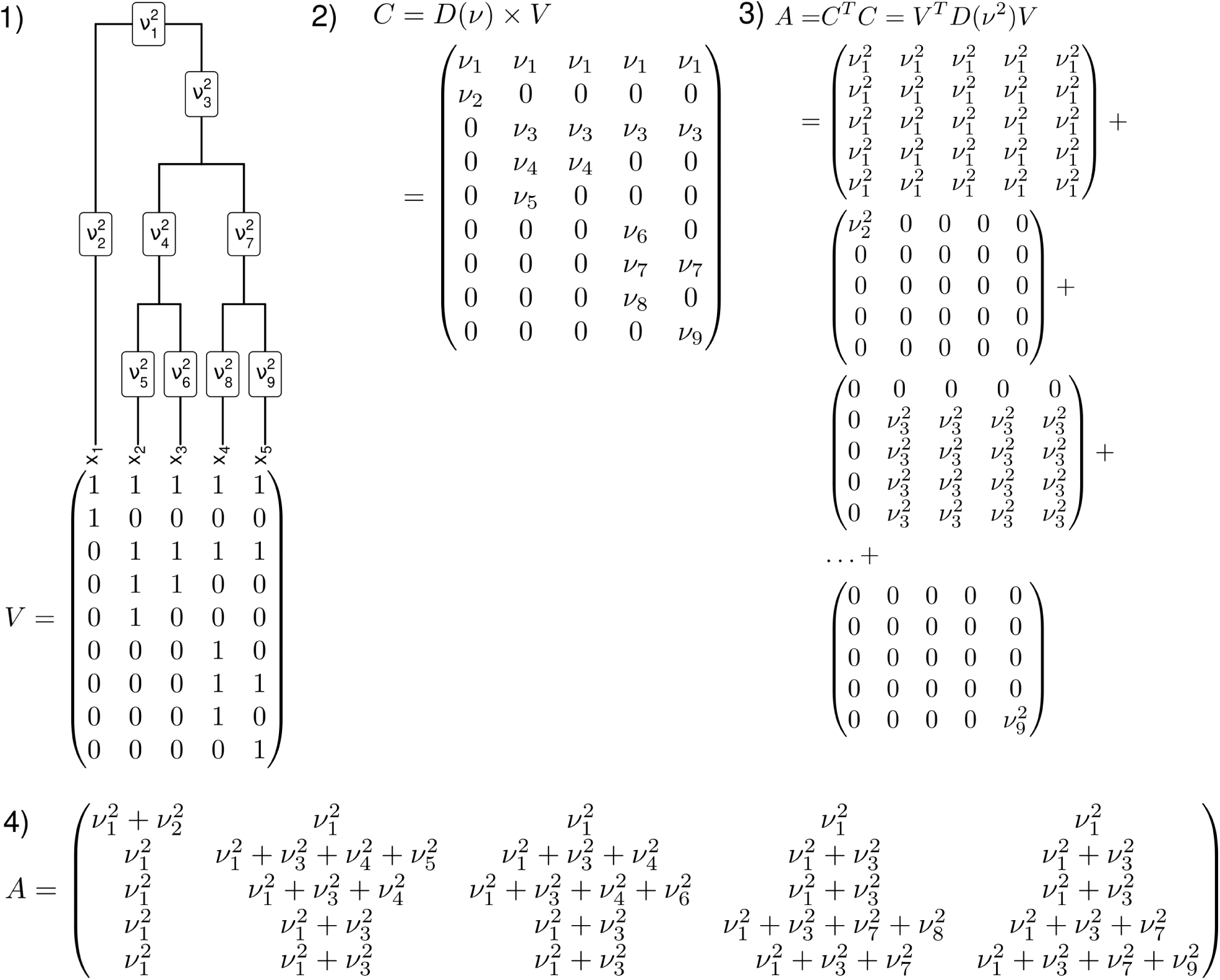
Consumer-resource framework for model 1. 1) Each clade has access to a “private” resource, as can access the resource). 2) Each resource has a value, stored in vector *ν*; in this case the consumption matrix (*ν_j_*^2^) for each clade a species belongs to. 4) The resulting matrix encoded in matrix *V* (the rows are the resources, the columns the species, and a coefficient is one if the species is simply *C* = *D*(*ν*)*V*. 3) The competition matrix *A* is obtained by summing the competition coefficients 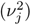 for each clade a species belongs to. 4) The resulting matrix has a nested block structure, with a block per clade.

We consider *n* consumer species with a specified phylogeny. We imagine a situation in which each clade in the tree has access to a clade-specific resource. Thus we can build a matrix *V* with 2*n*−1 rows (the resources, one per clade, including a resource accessible to all species) and *n* columns (the consumers). The matrix has a coefficient of *V_ij_* = 1 whenever consumer *j* has access to resource *i*, and zero elsewhere. Next, we model consumption, by assuming that each resource has a certain value (taken to be the same irrespective of the consumer), and that each consumer has a given uptake rate (taken to be the same for each resource):

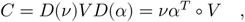

where *D*(*ν*) is a diagonal matrix with the value of each resource on the diagonal, *D*(*α*) a diagonal matrix of uptake rates and ○ is the Hadamard product. We obtain a system of 2*n* − 1 + *n* differential equations, which follow the Generalized Lotka-Volterra model (MacArthur consumer-resource model MacArthur (1970)):

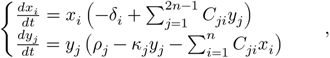

in which *x_i_* = *x_i_*(*t*) is the density of consumer *i*, *y_j_* = *y_j_*(*t*) is the density of resource *j*, and all parameters are positive. *δ_i_* is the intrinsic death rate of consumer *i*, *ρ_j_* is the growth rate of resource *j* and *κ_j_* is its self-limitation. If we assume that resource dynamics quickly equilibrate for any consumer densities (practically, if 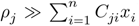), we can perform a “separation of time-scales” (as in MacArthur 1970), and solve the equation for the equilibrium of *y_j_*:

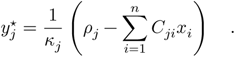

Plugging these values into the equations for the consumers yields:

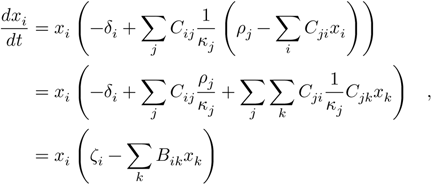

where we have defined *ζ_i_* = (∑*_j_ C_ij_ρ_j_*/*κ_j_*) − *δ_i_* and *B* = *C^T^ D*(*κ*)^−1^*C*. Finally, we factor *ζ_i_* outside the parenthesis, obtaining:

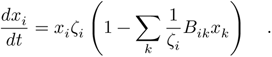

The matrix of species’ interactions is therefore:

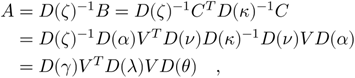

where *θ* = *α*, 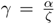 and 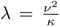. This is exactly the Eq. 1 in the main text, which we use for model 4. Further assuming that *ζ* = 1/*α* we obtain model 2; choosing *α* = 1 yields model 3 and, finally, using both simplifications we obtain model 1. Figure 5 illustrates the parameterization for the case of the simplest model: *A* = *V^T^D*(*λ*)*V*. We note that the variance-covariance matrix of a tree with branch lengths *λ* would take same form; as such, an alternative parameterization is thus to take competition coefficients to be the result of evolution via Brownian motion of many traits (Ives and Helmus 2011; Serván et al. 2020).

### Analysis of previous approaches

Traditional approaches used to fit data from Biodiversity-Ecosystem Functioning experiments rely on regression models that consider the effects of biodiversity, in terms of species richness and composition, and some measurement of ecosystem function, such as biomass, carbon fixation, and litter decomposition. These models usually consider total biomass of each plot as the response variable, and regress it against the number of species or other properties of the plots; for example, they consider a design matrix of zeros and ones detailing the presence/absence of each species in a given plot. These models can be supplemented by measures of phylogenetic distance between the species, species’ functional groups, species-specific effects, pairwise interactions, etc. Because of their underlying structure, these models do not take full advantage of the data, which usually report individual biomasses for each species in each plot—these data are aggregated when computing the total biomass per plot. These models suffer from three main limitations: (i) the use of aggregate data to fit the models when measurements are made at the species-level; (ii) lack of consistency with population dynamical models (but see Parain et al. 2019); and (iii) underlying assumptions regarding the error structure of the statistical models, which are not in accordance with the structure of the data or the assumptions of the model. Following the formulation by Connolly et al. (2011), a typical regression model might have the following structure:

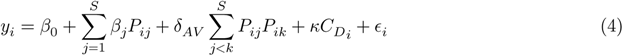

where *y_i_* is a measurement of ecosystem function for plot *i* (e.g., total biomass), *S* is the total number of species considered in the experiment (total species richness), *β*_0_ is an intercept, *β_j_* is the effect of species *j* on function in monoculture, *P_ij_* is an indicator variable for the presence (or absence) of species *j* in plot *i*, *δ_AV_* is the average interaction effect, 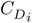 is the phylogenetic diversity in plot *i*, computed as 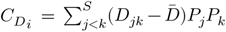, where *D_jk_* is the phylogenetic distance between species *j* and *k* (accounted as the sum of branch lengths between the two species) and 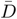 is the average distance of all species in the pool), and finally *∈_i_* is the error term.

The amount of data needed to fit such a model can be very large. To examine the performance of this kind of regression model, we therefore use simulations where data is generated in agreement with basic biological assumptions and test the model with the simulated data. Here we simulate ecological communities of competing species following the Generalized Lotka-Volterra dynamics, with the matrix of interactions given by the variance-covariance matrix corresponding to a phylogenetic tree. This should be an ideal situation for fitting the model, as interactions are constant and strictly pairwise, with a clear phylogenetic structure. Starting from a pool of *n* = 10 species, we assume all species have the same intrinsic growth rates, so that dynamics are determined solely by phylogenetic relations. We draw a random tree, construct its variance-covariance matrix, assemble all 2*^n^* − 1 = 1023 communities that can be formed from the species pool and record each species’ biomass in each community. The structure of the matrix of interactions can be thought of as emerging from models of trait evolution; for example, if we follow the evolution of many trait values, each changing under Brownian motion, this process gives rise to the trait variance-covariance matrix which we associated with competition coefficients. In this way, differences in trait values are proportional to evolutionary distances (Hansen and Martins 1996; Ives and Helmus 2011). One key aspect of this parameterization is that all sub-communities that can be formed with a subset of species are feasible and dynamically stable (see Serván et al. 2020 for details).

Having produced a large synthetic data set containing more than a thousand “plots”, we can assess the quality of fit using a variety of regression models of varying complexity. Model *M*0 : *y* ∼ *P* regresses total biomass against the presence/absence of each species (*P*); model *M*1 : *y* ∼ *P* + *C_D_* regresses the total biomass against presence/absence of species (*P*) and the phylogenetic distance of species in the plot (*C_D_*); *M*2 : *y* ∼ *P* + *C_D_* + *P* : *C_D_* is the same as *M*1 but also considers the interaction between presence and phylogenetic distance; *M*3 : *y* ∼ *P* + ∑*P_i_P_j_* considers the presence absence of each species and their pairwise interaction; *M*4 : *y* ∼ *P* +∑*P_i_P_j_* +*C_D_* is the same as *M*3 adding the phylogenetic distance of each plot; and and phylogenetic finally *M*5 : *y* ∼ *P* + ∑*P_i_P_j_* + *C_D_* + *P* : *C_D_* is the same as *m*4 including the interaction between presence and phylogenetic distance. Below we show the plots for each of the models, including the predicted versus observed biomasses and diagnostic plots from linear regression.

These results show that a) none of the models can fully capture the simulated data (despite the lack of noise); b) all of the model fits violate some of the main assumptions of linear regressions: residuals are not independent and normally distributed, variances are not constant, and thus Quantile-Quantile plots deviate substantially from the 1:1 line. We conclude that these model structures are not adequate to capture models for population dynamics—generalizing beyond competition, choosing more complex models for population dynamics, or introducing stochasticity and measurement error can only worsen the fit.

The inadequacy of regression-based approaches motivates our alternative methodology for inferring phylogenetic structure in these data.

**Figure 6:**
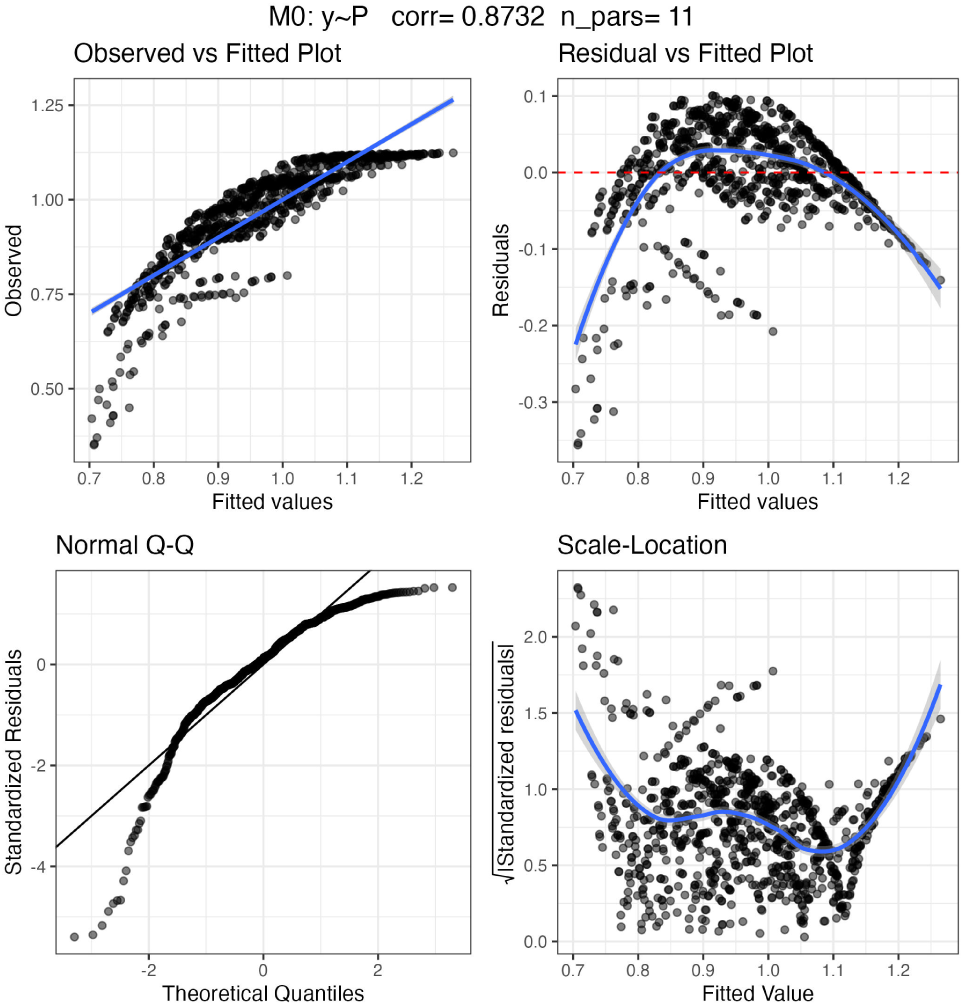
Fitting regression models to simulated data from a phylogenetically-structured Generalized Lotka-Volterra model. Each figure reports the structure of the model (top) as well as scatterplots showing the relationship between observed and fitted total biomasses per plot; corresponding distribution of standardized residuals (for linear regression, these should be centered at zero and have the same variance irrespective of the fitted values); Quantile-Quantile plots (points should fall on the 1 to 1 line); scale location plots, showing whether residuals change with the input (they should form a horizontal band of points).

**Figure 7:**
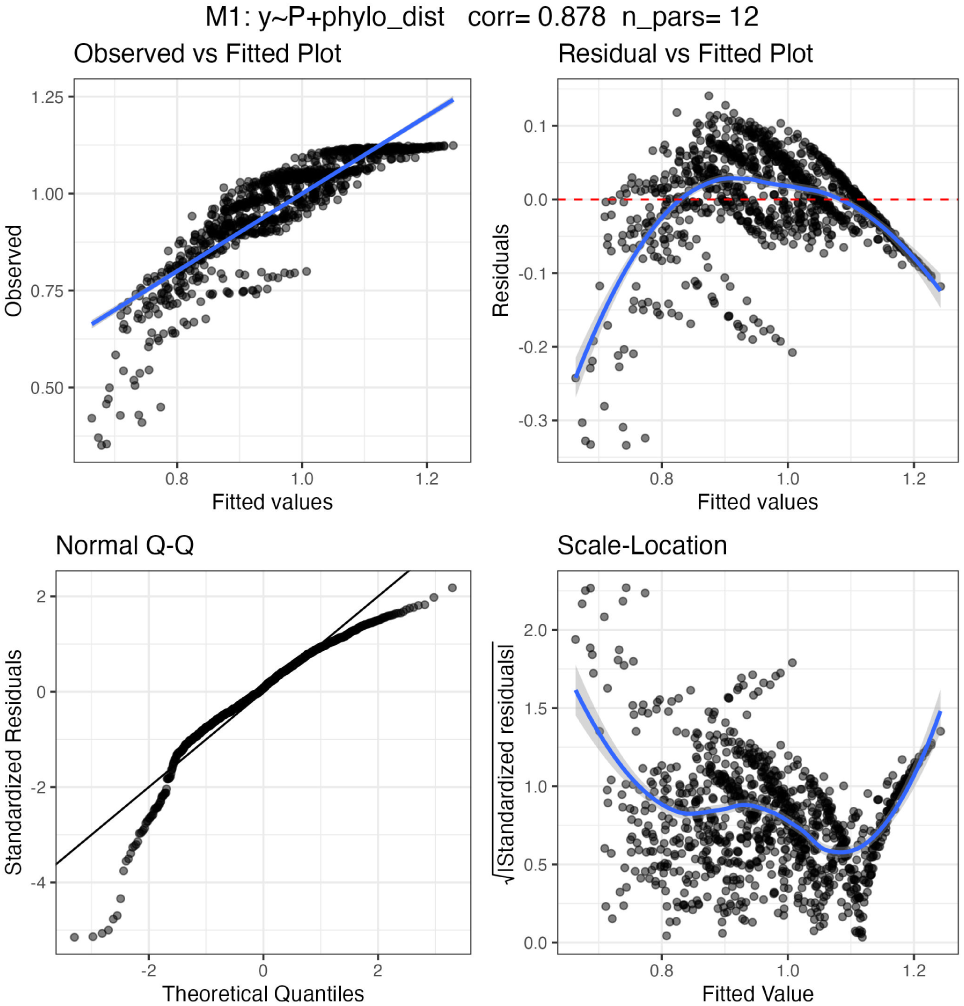
Fitting regression models to simulated data from a phylogenetically-structured Generalized Lotka-Volterra model. Each figure reports the structure of the model (top) as well as scatterplots showing the relationship between observed and fitted total biomasses per plot; corresponding distribution of standardized residuals (for linear regression, these should be centered at zero and have the same variance irrespective of the fitted values); Quantile-Quantile plots (points should fall on the 1 to 1 line); scale location plots, showing whether residuals change with the input (they should form a horizontal band of points).

**Figure 8:**
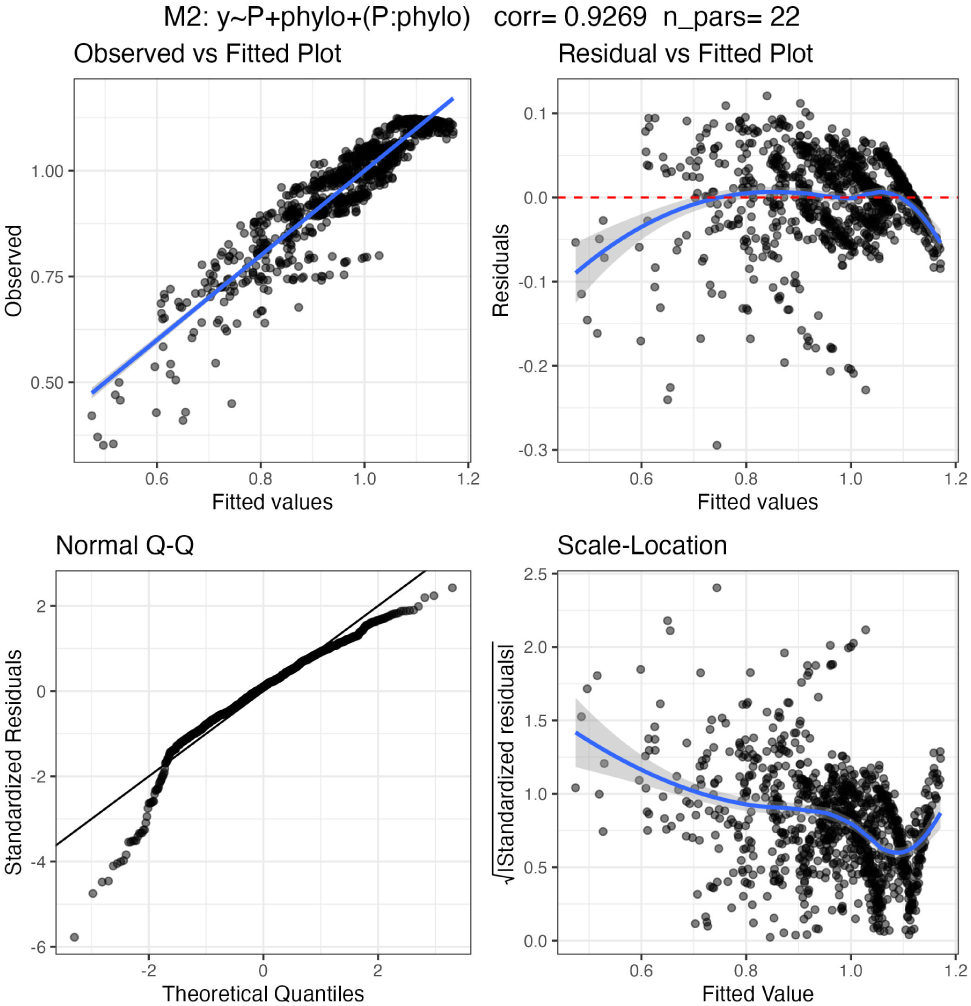
Fitting regression models to simulated data from a phylogenetically-structured Generalized Lotka-Volterra model. Each figure reports the structure of the model (top) as well as scatterplots showing the relationship between observed and fitted total biomasses per plot; corresponding distribution of standardized residuals (for linear regression, these should be centered at zero and have the same variance irrespective of the fitted values); Quantile-Quantile plots (points should fall on the 1 to 1 line); scale location plots, showing whether residuals change with the input (they should form a horizontal band of points).

**Figure 9:**
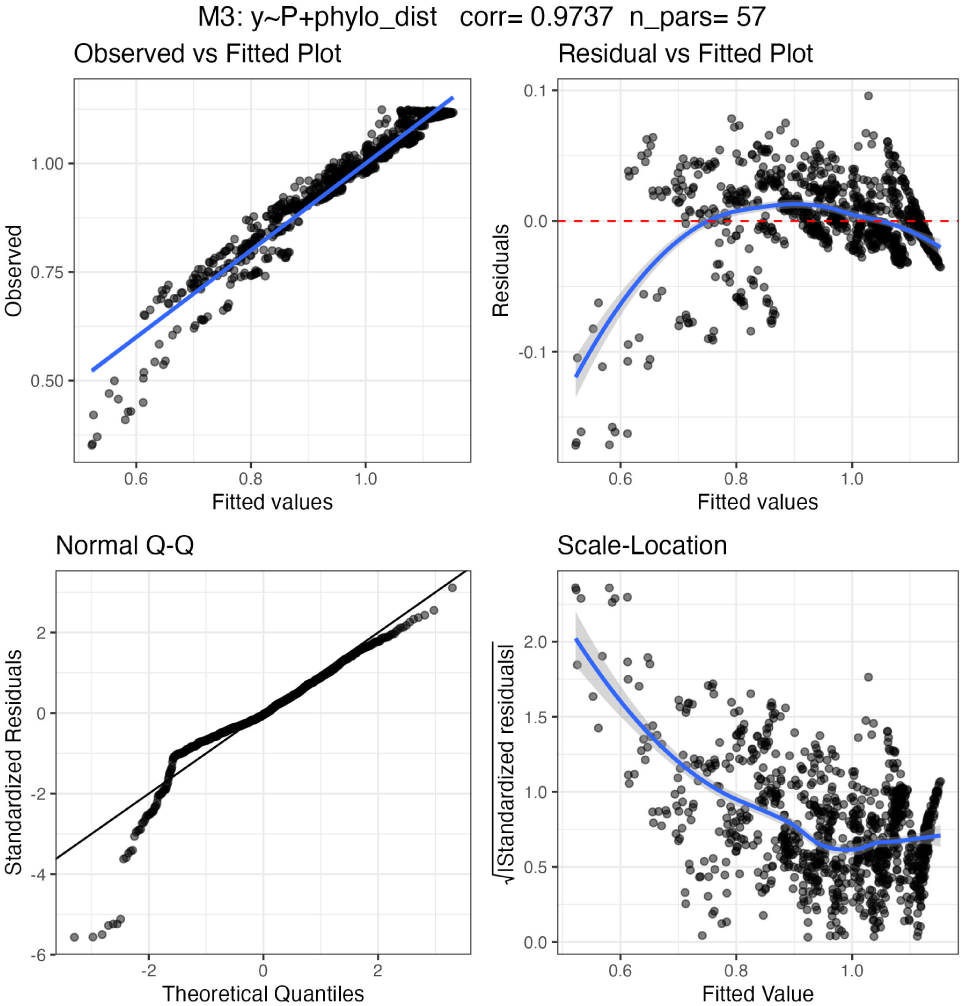
Fitting regression models to simulated data from a phylogenetically-structured Generalized Lotka-Volterra model. Each figure reports the structure of the model (top) as well as scatterplots showing the relationship between observed and fitted total biomasses per plot; corresponding distribution of standardized residuals (for linear regression, these should be centered at zero and have the same variance irrespective of the fitted values); Quantile-Quantile plots (points should fall on the 1 to 1 line); scale location plots, showing whether residuals change with the input (they should form a horizontal band of points).

**Figure 10:**
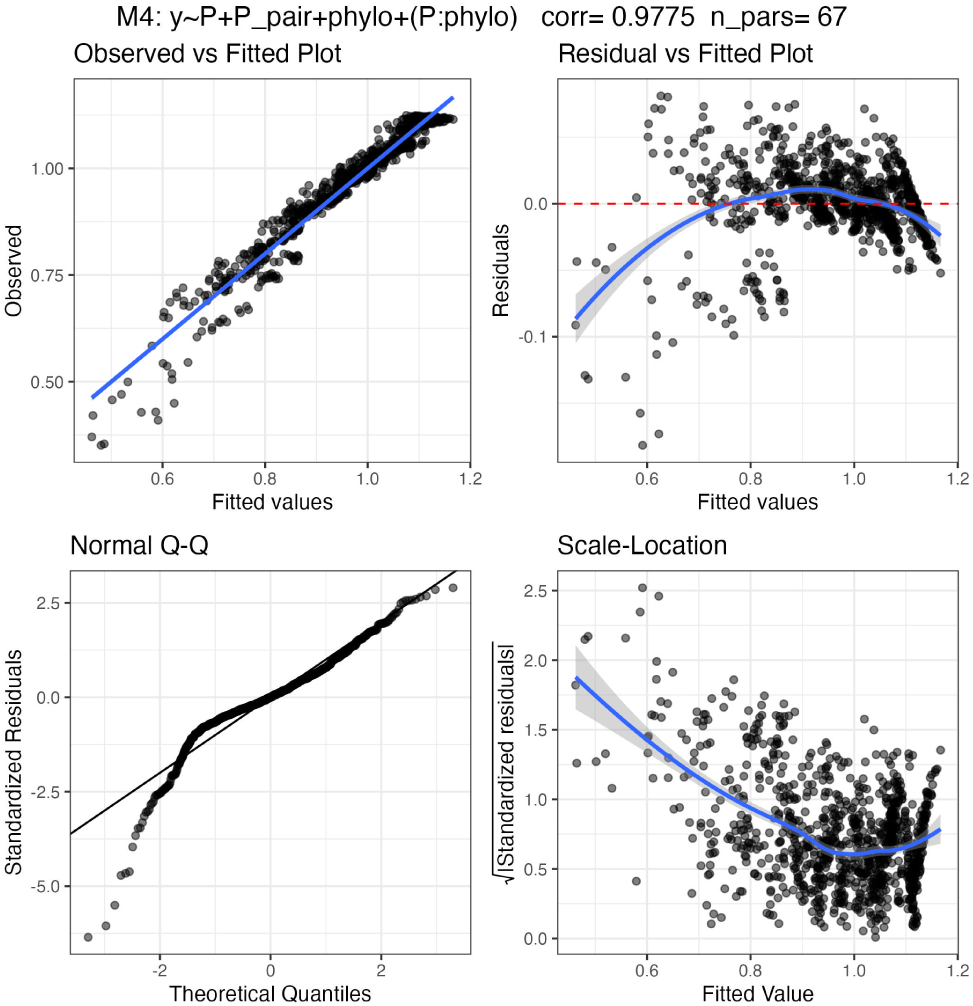
Fitting regression models to simulated data from a phylogenetically-structured Generalized Lotka-Volterra model. Each figure reports the structure of the model (top) as well as scatterplots showing the relationship between observed and fitted total biomasses per plot; corresponding distribution of standardized residuals (for linear regression, these should be centered at zero and have the same variance irrespective of the fitted values); Quantile-Quantile plots (points should fall on the 1 to 1 line); scale location plots, showing whether residuals change with the input (they should form a horizontal band of points).

### Full results

The results for each data set are reported in Table 1. Scatterplots for the data stemming from the Biodiversity II and the Wageningen experiments are reported in Figure 11 and Figure 13, respectively. Histograms of the likelihoods for the randomized trees are reported in Figure 12 and Figure 14. The fitted trees and associated likelihood boxplots for randomizations stratified by selected clades are reported in Figure 15 – Figure 22 for the Biodiversity II experiment and Figure 23 – Figure 30 for the Wageningen experiment. Finally, the fits for models 2-4 with the associated tree and vectors are also in **?@fig-models2-4**.

**Figure 11:**
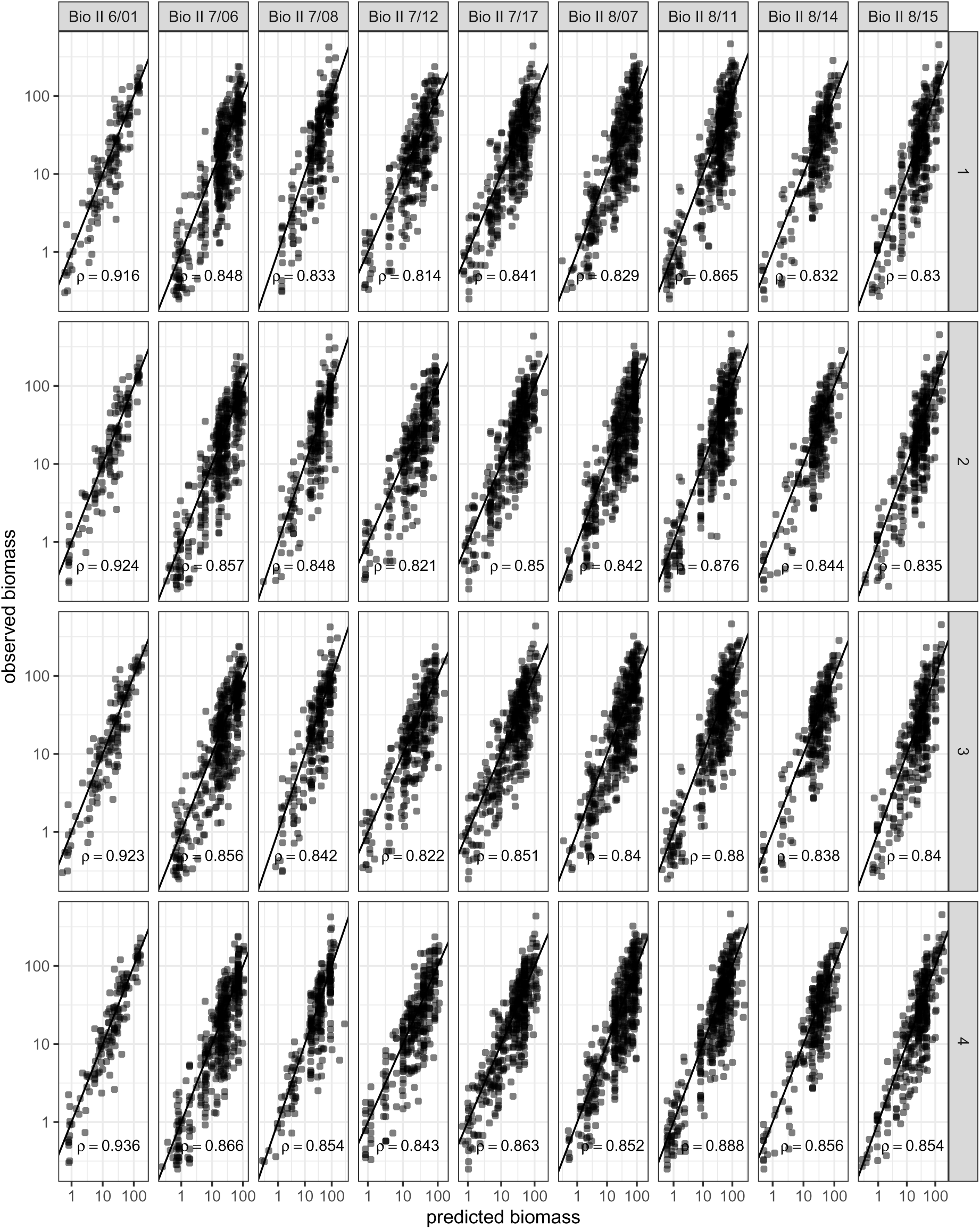
Scatterplot of predicted vs. observed biomass for the Biodiversity II data. Each panel represents a specific sampling date and model. The correlation between the logarithm of the observed biomass and the logarithm of the predicted biomass is reported in each panel.

**Figure 12:**
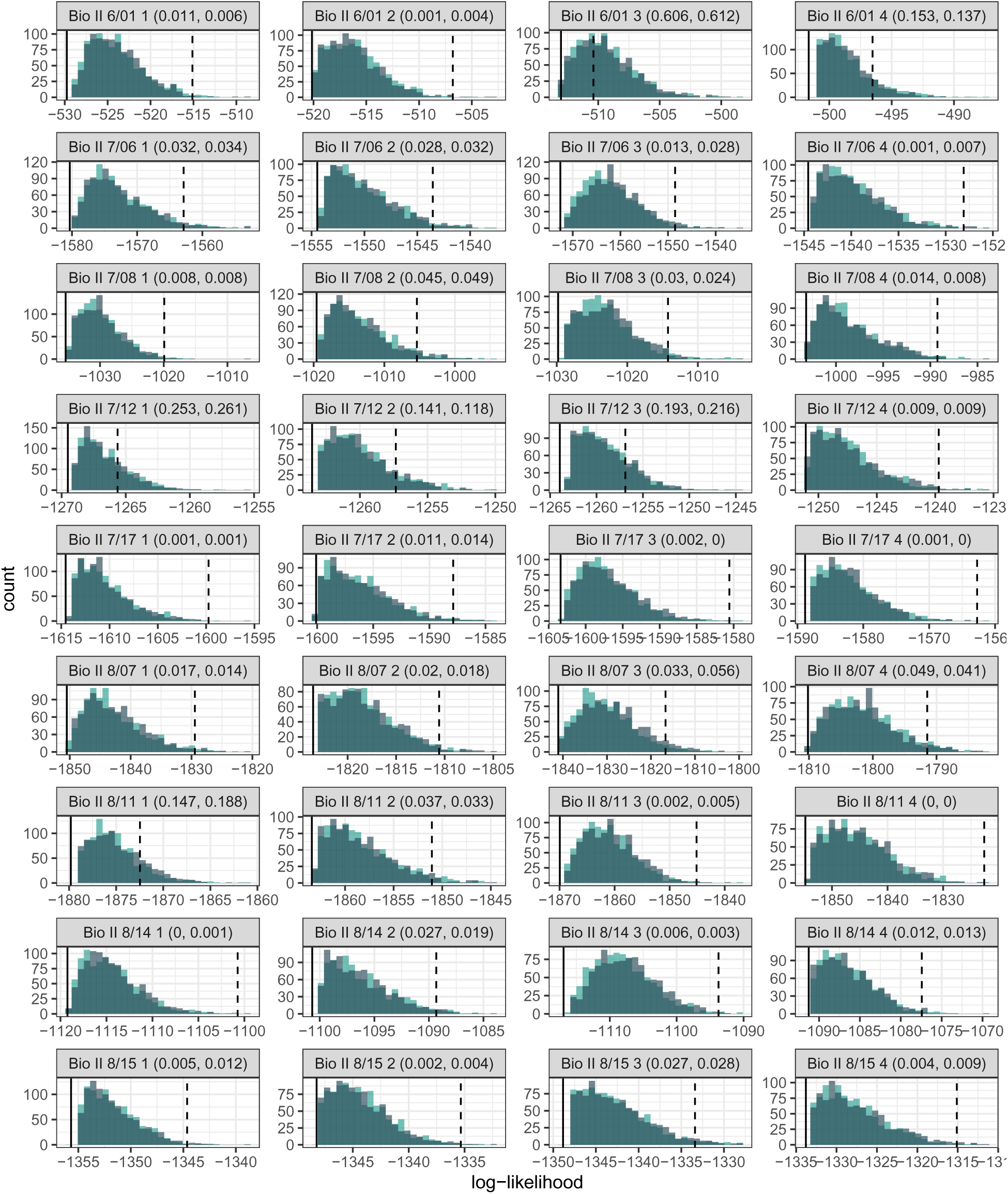
Histograms of the log-likelihoods for models parameterized using random trees and data from the Biodiversity II experiment. The distributions for the random trees generated by the Yule model are in teal, and those for the shuffled trees in grey. Each panel reports the p-values obtained for the Yule and shuffled trees, respectively.

**Figure 13:**
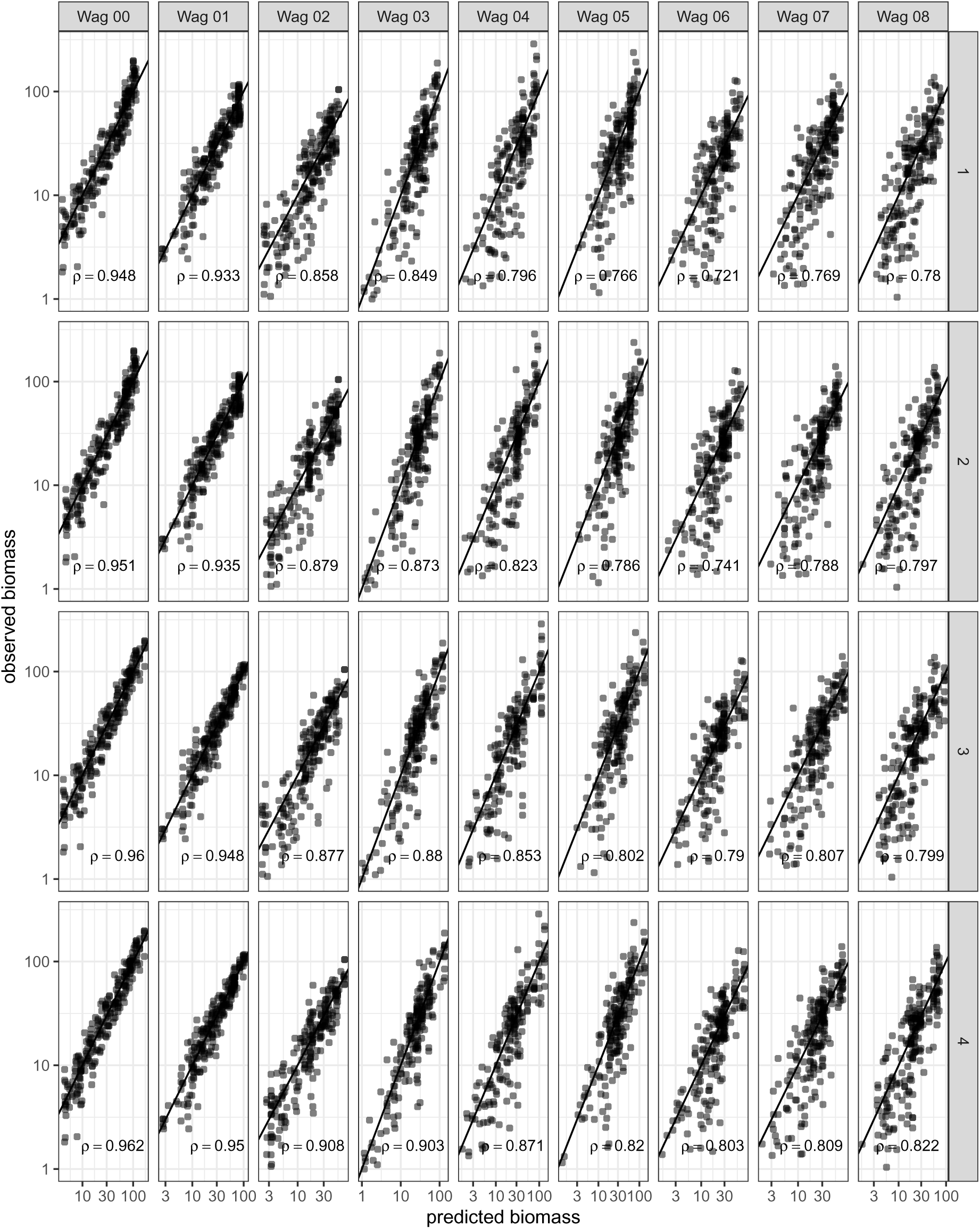
Scatterplot of predicted vs. observed biomass for the Wageningen biodiversity experiment data. Each panel represents a specific sampling date and model. The correlation between the logarithm of the observed biomass and the logarithm of the predicted biomass is reported in each panel.

**Figure 14:**
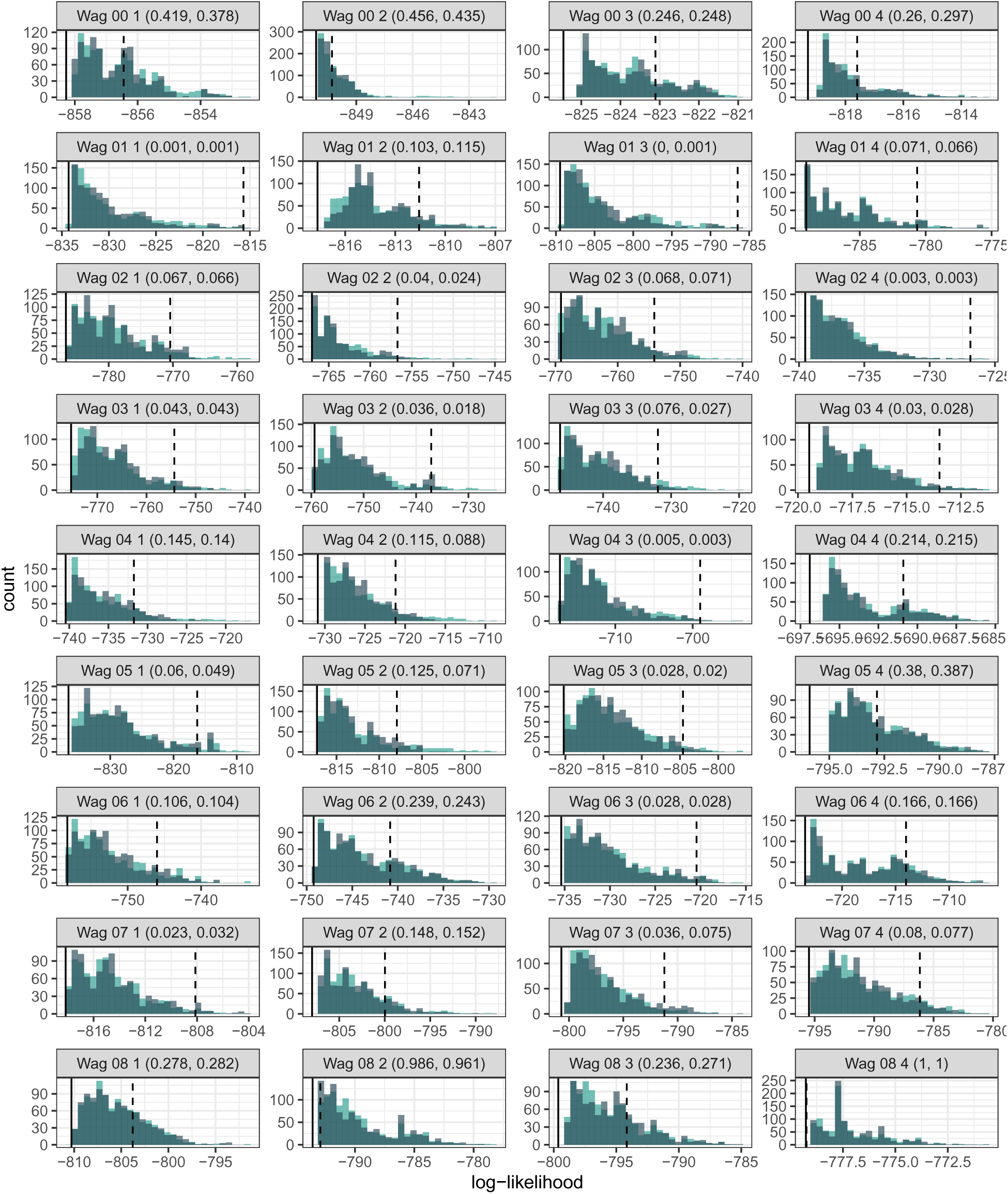
Histograms of the log-likelihoods for models parameterized using random trees and data from the Wageningen diversity experiment. The distributions for the random trees generated by the Yule model are in teal, and those for the shuffled trees in grey. Each panel reports the p-values obtained for the Yule and shuffled trees, respectively.

**Figure 15:**
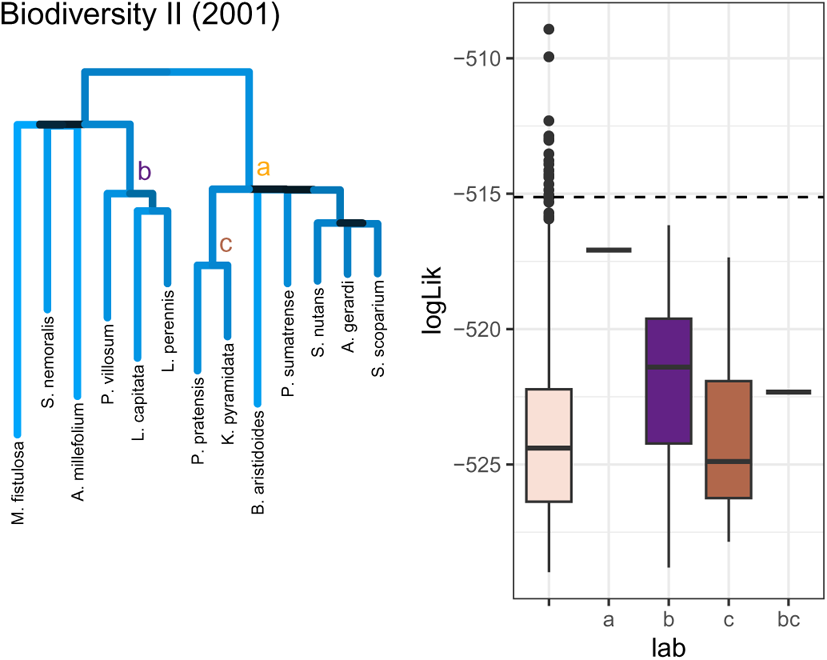
Resulting trees for the Biodiversity II experiments in Cedar Creek, fit for phylogenetic tree from model 1 and boxplots of likelihoods for randomizations that kept the selected clades highlighted in the trees. Dashed line represents likelihood of the original tree

**Figure 16:**
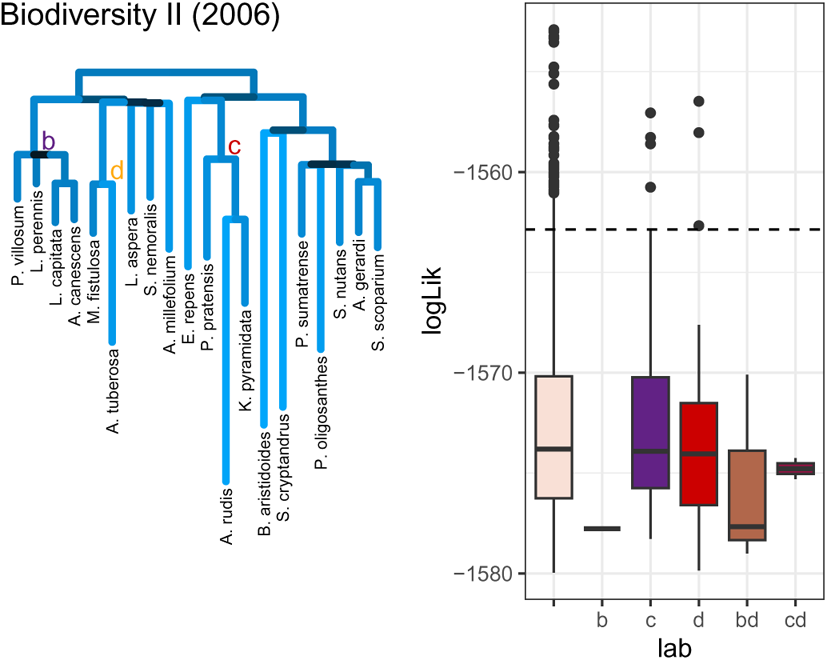
Resulting trees for the Biodiversity II experiments in Cedar Creek, fit for phylogenetic tree from model 1 and boxplots of likelihoods for randomizations that kept the selected clades highlighted in the trees. Dashed line represents likelihood of the original tree

**Figure 17:**
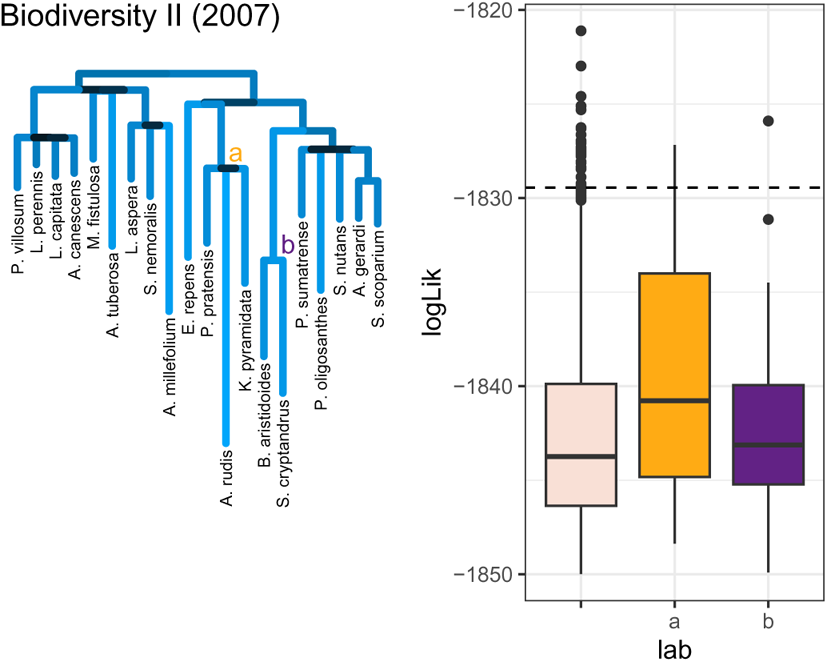
Resulting trees for the Biodiversity II experiments in Cedar Creek, fit for phylogenetic tree from model 1 and boxplots of likelihoods for randomizations that kept the selected clades highlighted in the trees. Dashed line represents likelihood of the original tree

**Figure 18:**
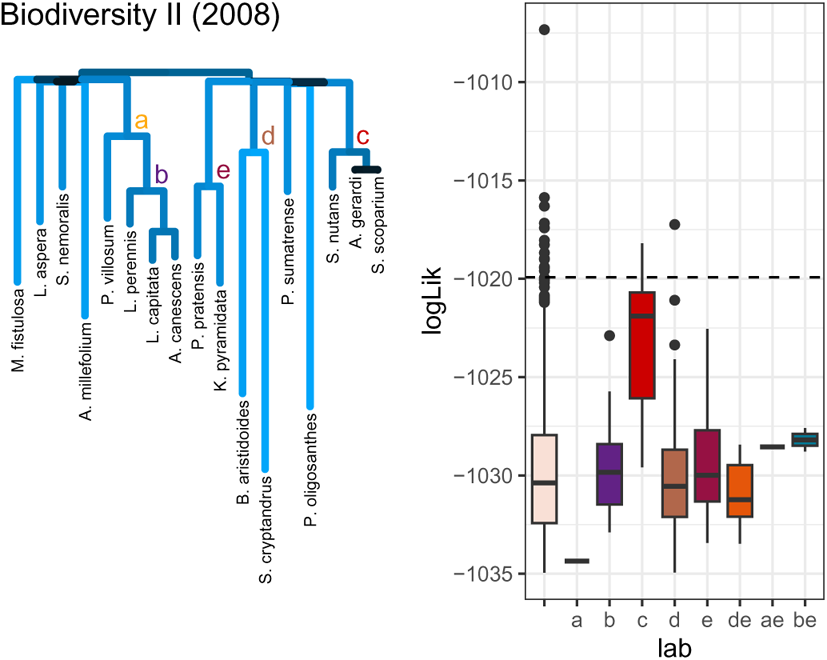
Resulting trees for the Biodiversity II experiments in Cedar Creek, fit for phylogenetic tree from model 1 and boxplots of likelihoods for randomizations that kept the selected clades highlighted in the trees. Dashed line represents likelihood of the original tree

**Figure 19:**
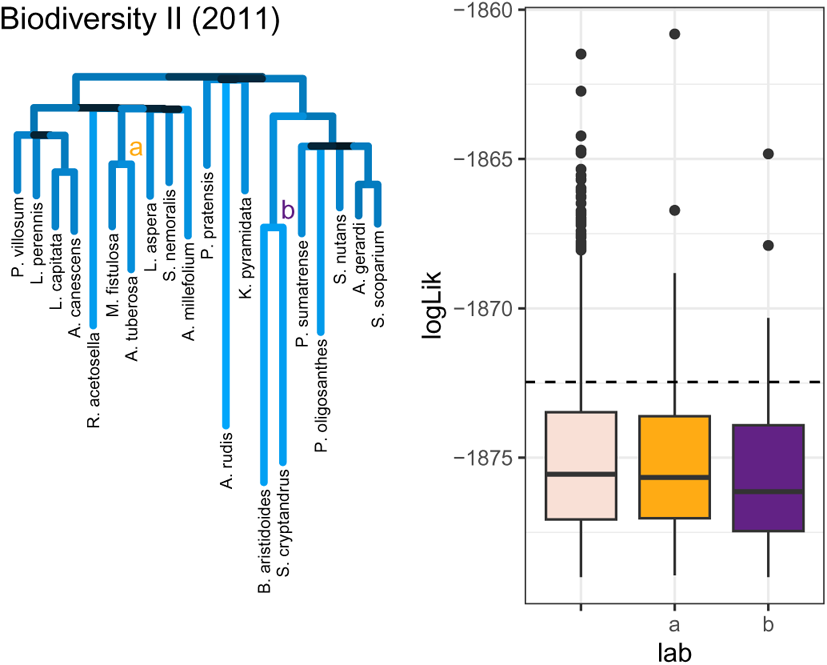
Resulting trees for the Biodiversity II experiments in Cedar Creek, fit for phylogenetic tree from model 1 and boxplots of likelihoods for randomizations that kept the selected clades highlighted in the trees. Dashed line represents likelihood of the original tree

**Figure 20:**
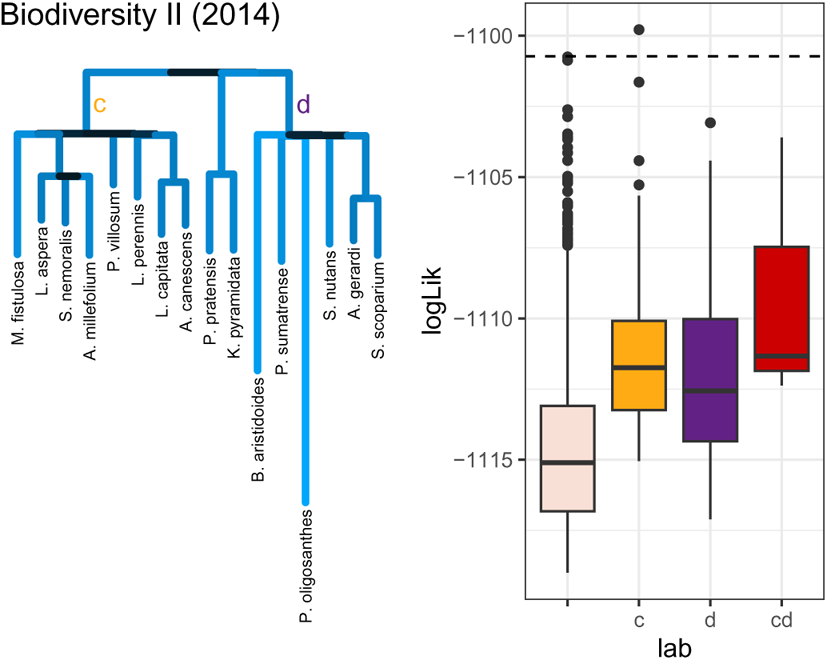
Resulting trees for the Biodiversity II experiments in Cedar Creek, fit for phylogenetic tree from model 1 and boxplots of likelihoods for randomizations that kept the selected clades highlighted in the trees. Dashed line represents likelihood of the original tree

**Figure 21:**
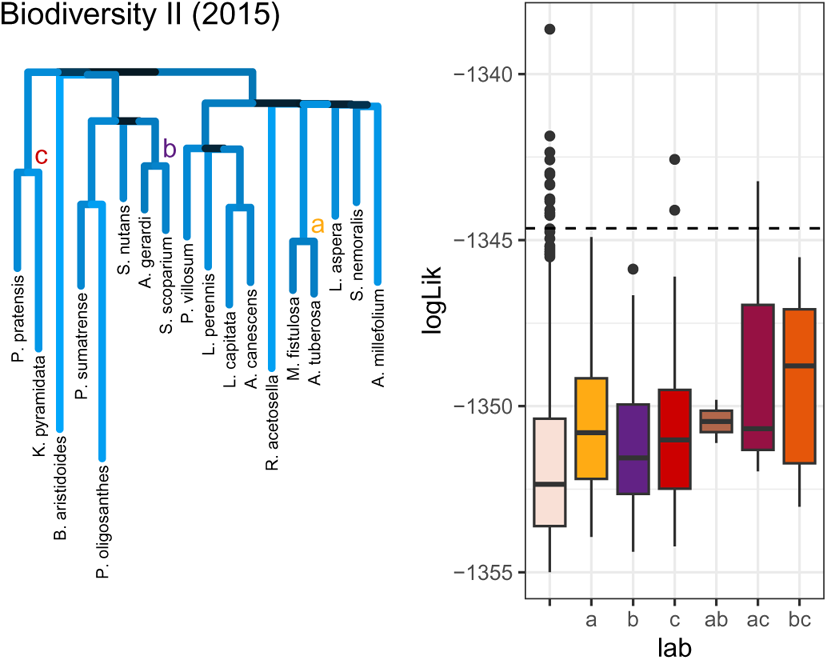
Resulting trees for the Biodiversity II experiments in Cedar Creek, fit for phylogenetic tree from model 1 and boxplots of likelihoods for randomizations that kept the selected clades highlighted in the trees. Dashed line represents likelihood of the original tree

**Figure 22:**
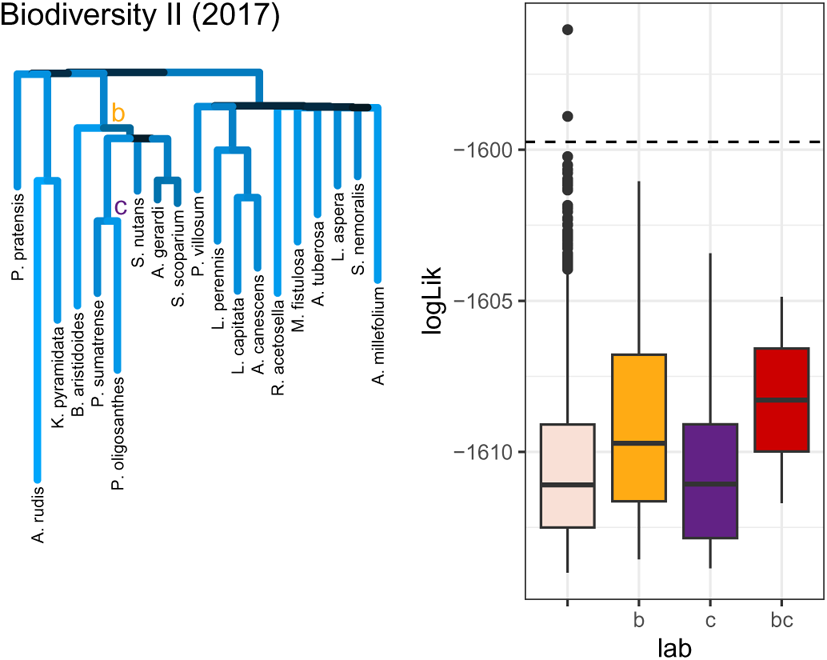
Resulting trees for the Biodiversity II experiments in Cedar Creek, fit for phylogenetic tree from model 1 and boxplots of likelihoods for randomizations that kept the selected clades highlighted in the trees. Dashed line represents likelihood of the original tree

**Figure 23:**
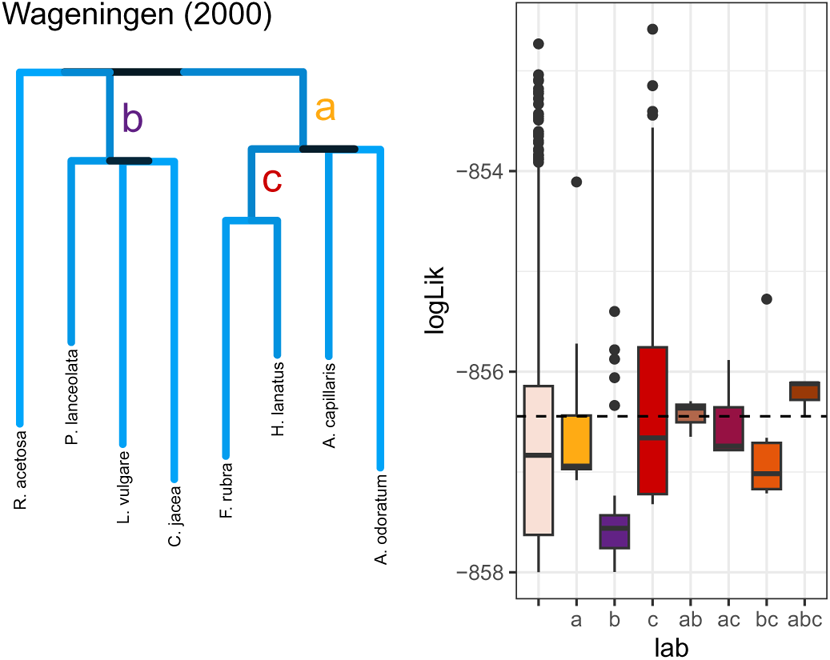
Resulting trees for the Wageningen experiments, fit for phylogenetic tree from model 1 and boxplots of likelihoods for randomizations that kept the selected clades highlighted in the trees. Dashed line represents likelihood of the original tree

**Figure 24:**
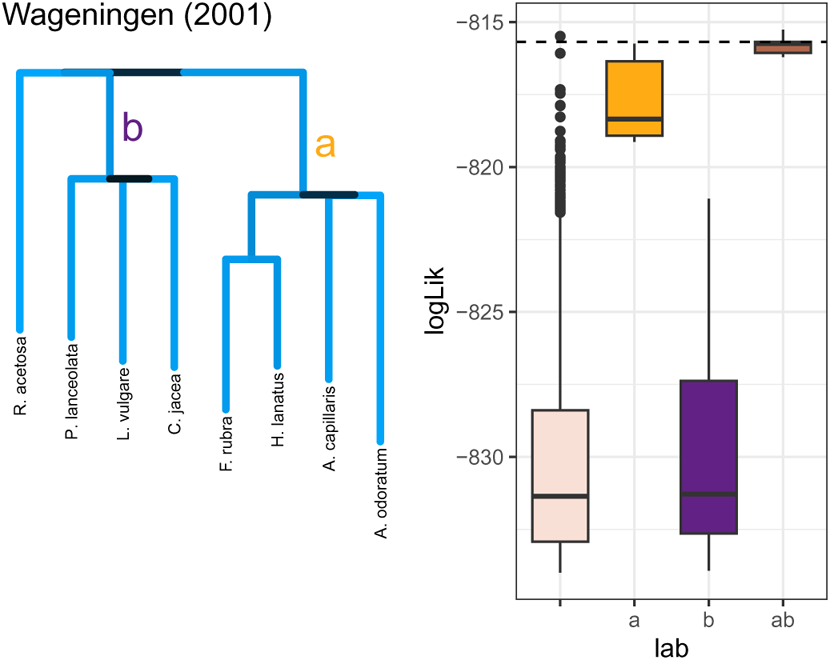
Resulting trees for the Wageningen experiments, fit for phylogenetic tree from model 1 and boxplots of likelihoods for randomizations that kept the selected clades highlighted in the trees. Dashed line represents likelihood of the original tree

**Figure 25:**
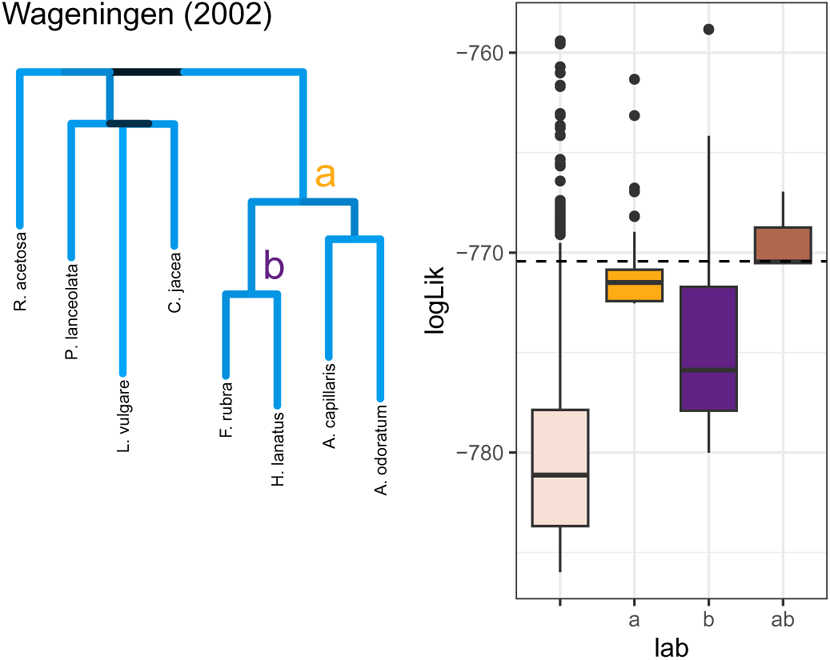
Resulting trees for the Wageningen experiments, fit for phylogenetic tree from model 1 and boxplots of likelihoods for randomizations that kept the selected clades highlighted in the trees. Dashed line represents likelihood of the original tree

**Figure 26:**
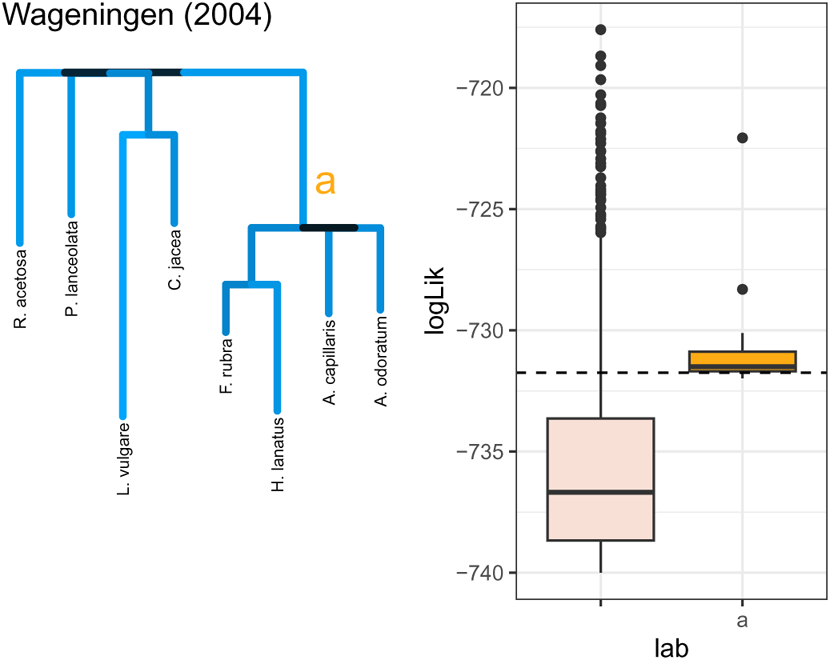
Resulting trees for the Wageningen experiments, fit for phylogenetic tree from model 1 and boxplots of likelihoods for randomizations that kept the selected clades highlighted in the trees. Dashed line represents likelihood of the original tree

**Figure 27:**
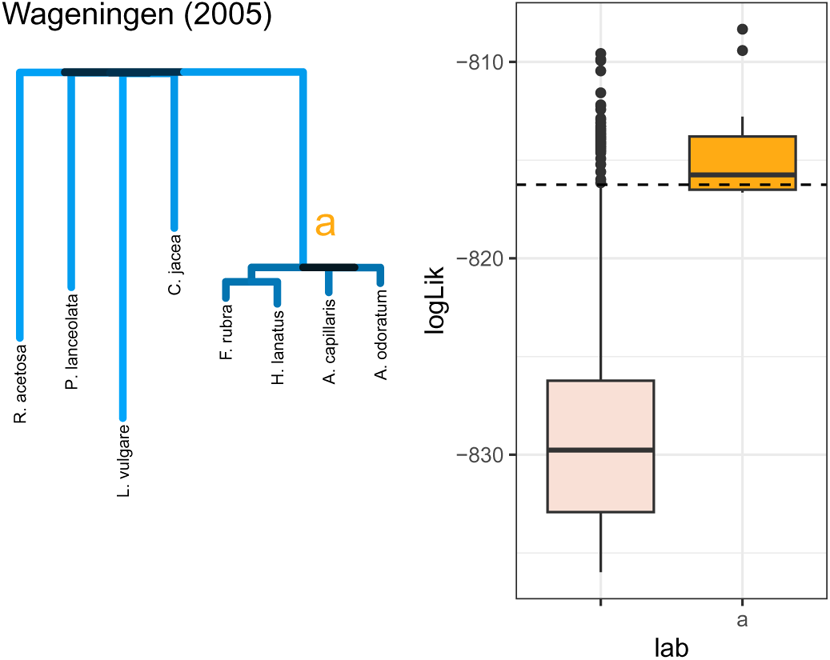
Resulting trees for the Wageningen experiments, fit for phylogenetic tree from model 1 and boxplots of likelihoods for randomizations that kept the selected clades highlighted in the trees. Dashed line represents likelihood of the original tree

**Figure 28:**
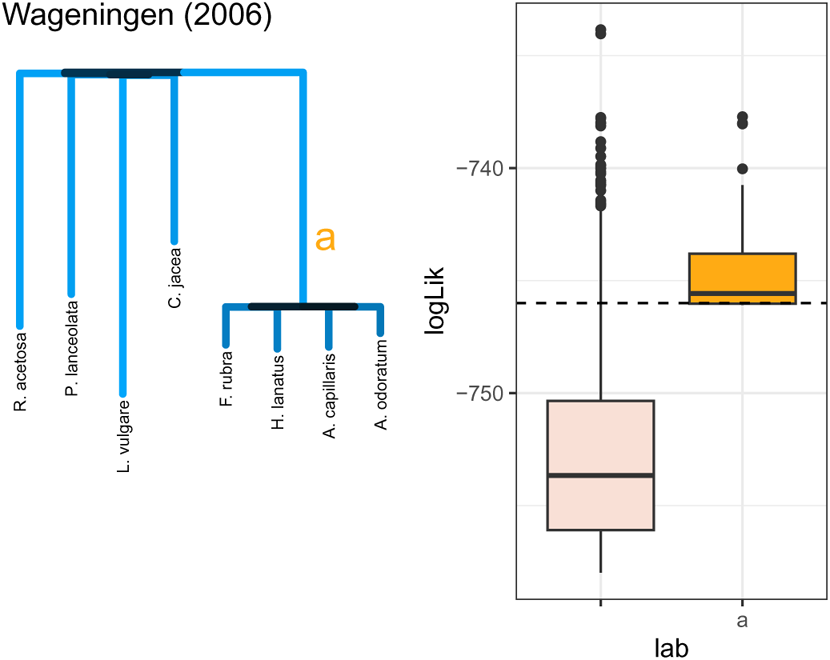
Resulting trees for the Wageningen experiments, fit for phylogenetic tree from model 1 and boxplots of likelihoods for randomizations that kept the selected clades highlighted in the trees. Dashed line represents likelihood of the original tree

**Figure 29:**
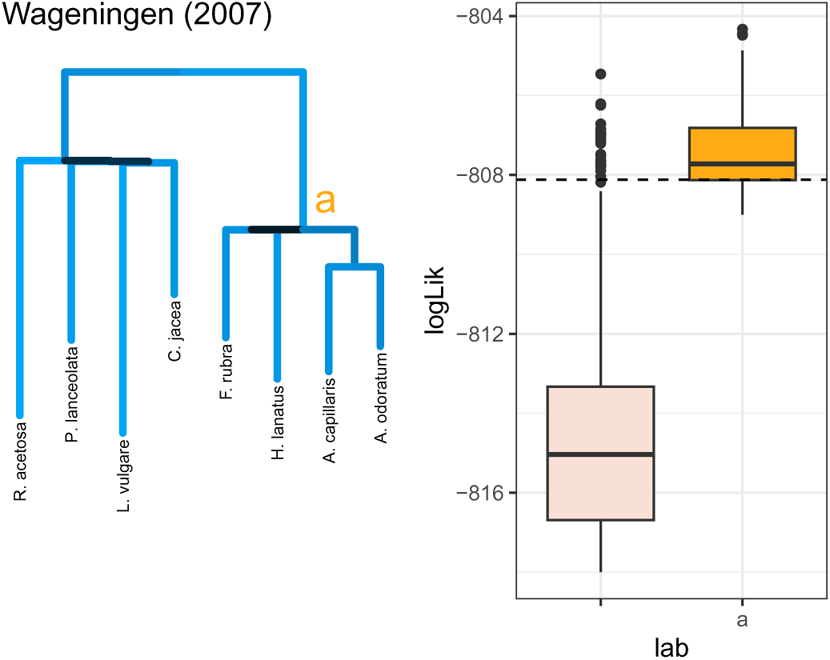
Resulting trees for the Wageningen experiments, fit for phylogenetic tree from model 1 and boxplots of likelihoods for randomizations that kept the selected clades highlighted in the trees. Dashed line represents likelihood of the original tree

**Figure 30:**
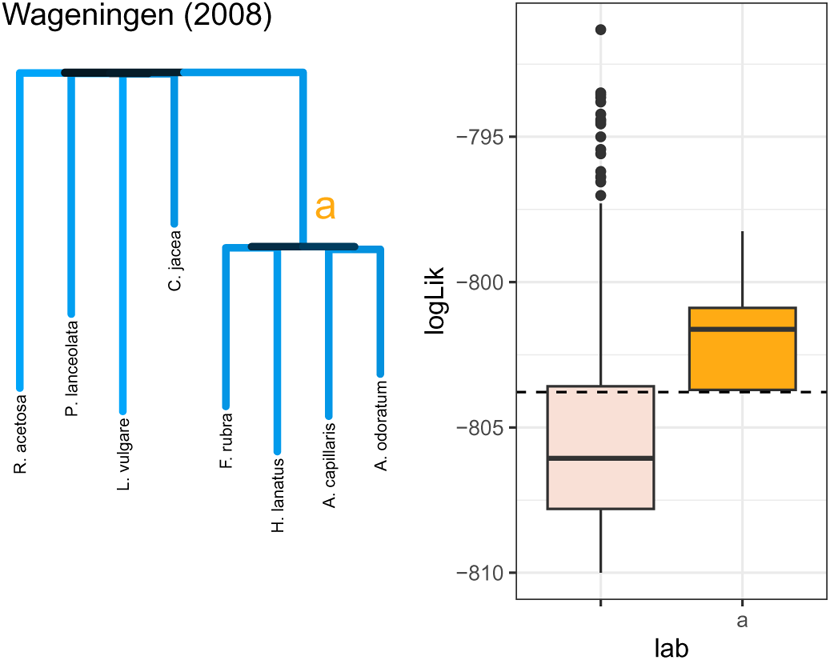
Resulting trees for the Wageningen experiments, fit for phylogenetic tree from model 1 and boxplots of likelihoods for randomizations that kept the selected clades highlighted in the trees. Dashed line represents likelihood of the original tree

**Figure 31:**
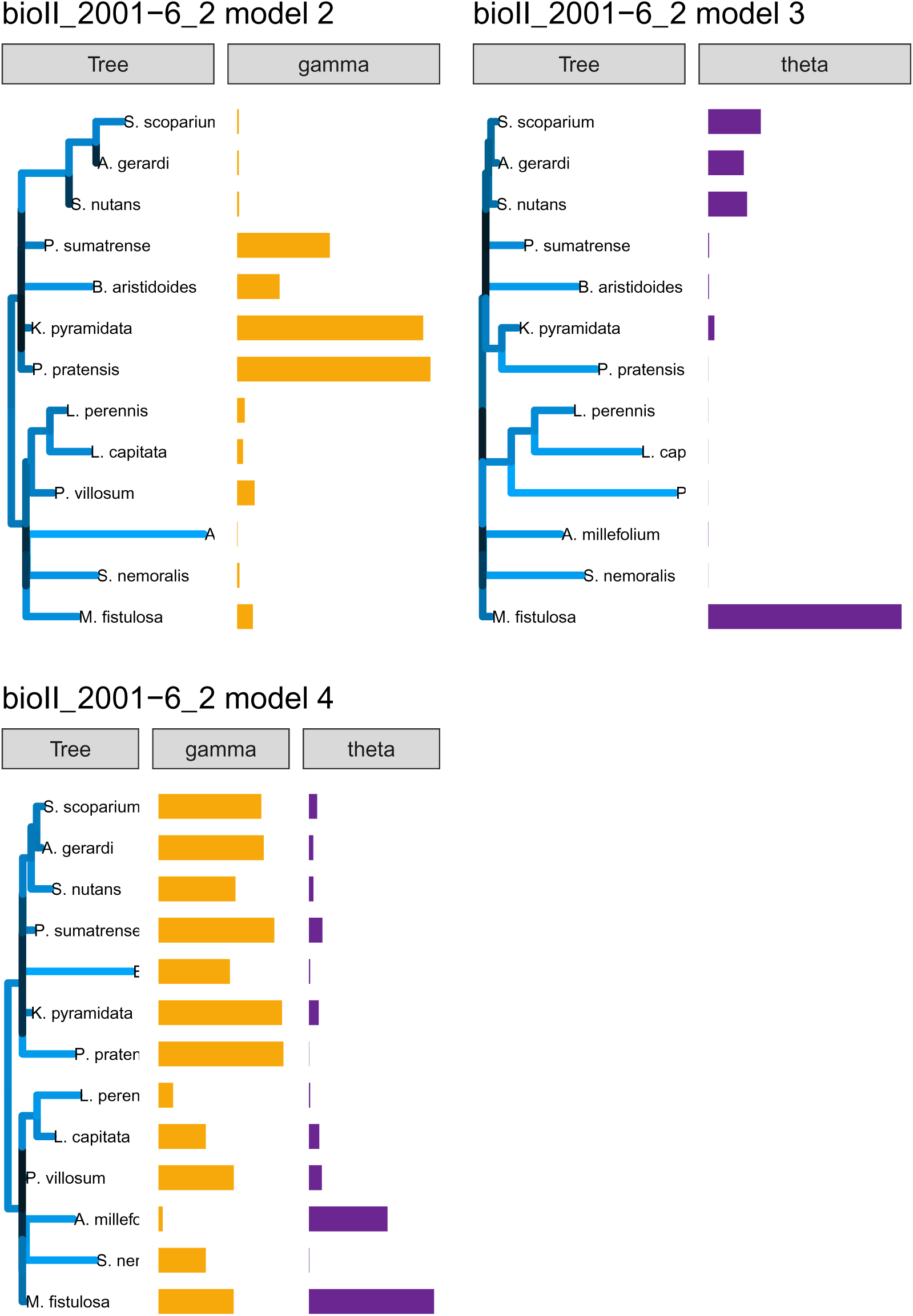
Fit for the phylogenetic tree and associated parameter vectors (Gamma and Theta) for models 2-4. The length and color of the branches correspond to increased strength in competition between the species in the clade and the size of the vector corresponds to the fitted value for the corresponding species in the tree. bioII refers to the Biodiversity II experiment (Tilman et al. (2001)); cadotte is from Cadotte (2013); and vr refers to the Wageningen experiment from Van Ruijven and Berendse (2010).

**Figure 32:**
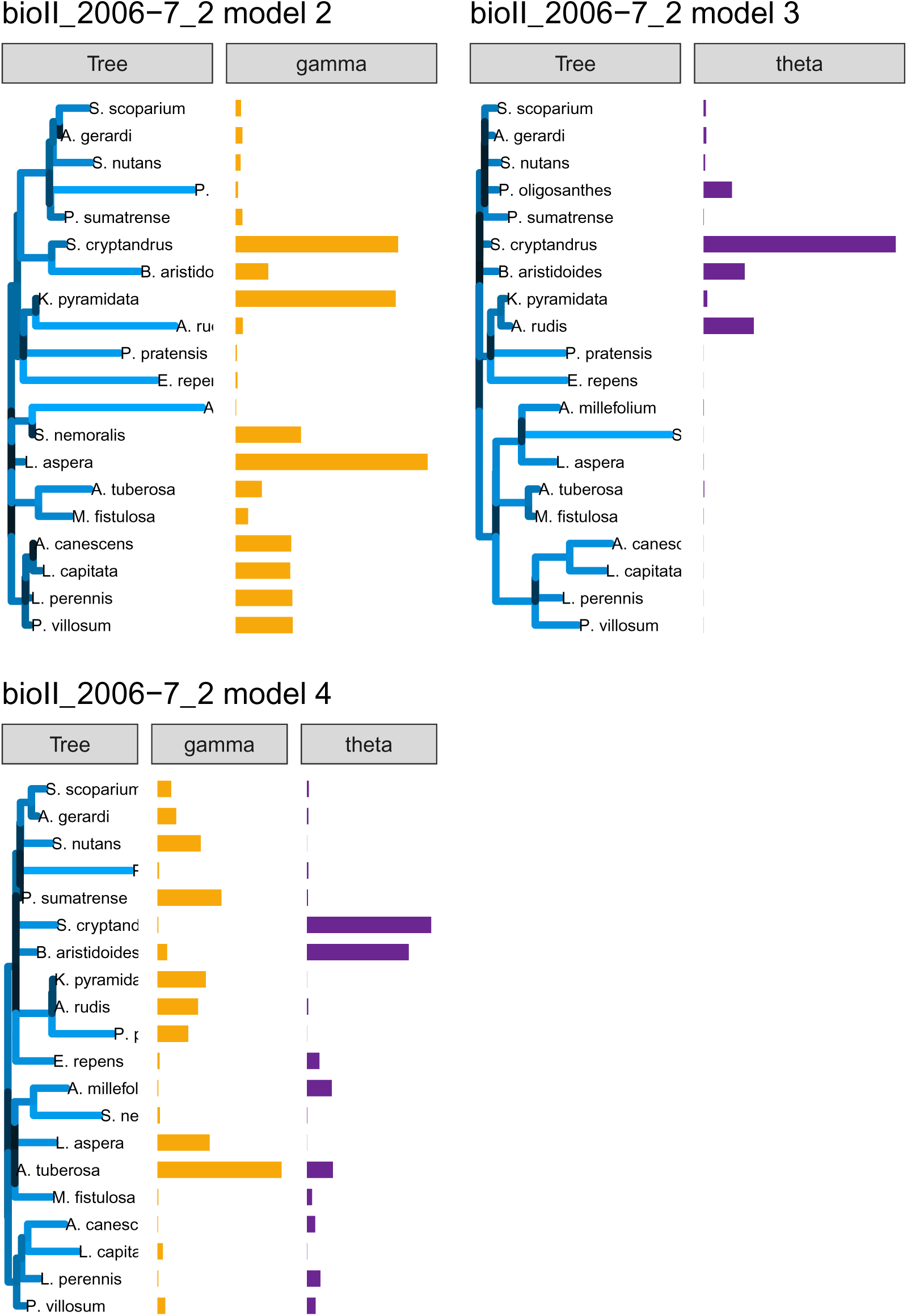
Fit for the phylogenetic tree and associated parameter vectors (Gamma and Theta) for models 2-4. The length and color of the branches correspond to increased strength in competition between the species in the clade and the size of the vector corresponds to the fitted value for the corresponding species in the tree. bioII refers to the Biodiversity II experiment (Tilman et al. (2001)); cadotte is from Cadotte (2013); and vr refers to the Wageningen experiment from Van Ruijven and Berendse (2010).

**Figure 33:**
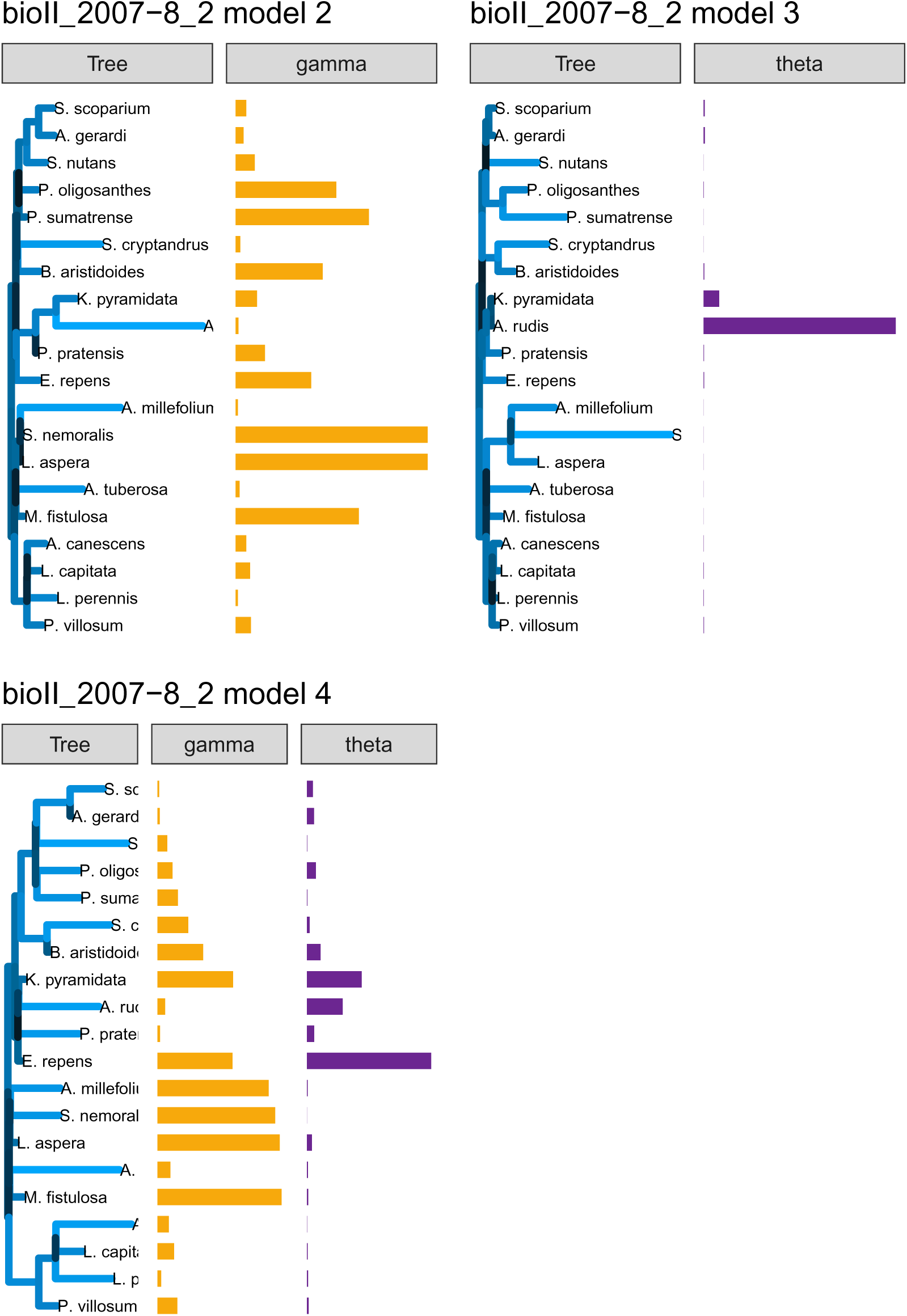
Fit for the phylogenetic tree and associated parameter vectors (Gamma and Theta) for models 2-4. The length and color of the branches correspond to increased strength in competition between the species in the clade and the size of the vector corresponds to the fitted value for the corresponding species in the tree. bioII refers to the Biodiversity II experiment (Tilman et al. (2001)); cadotte is from Cadotte (2013); and vr refers to the Wageningen experiment from Van Ruijven and Berendse (2010).

**Figure 34:**
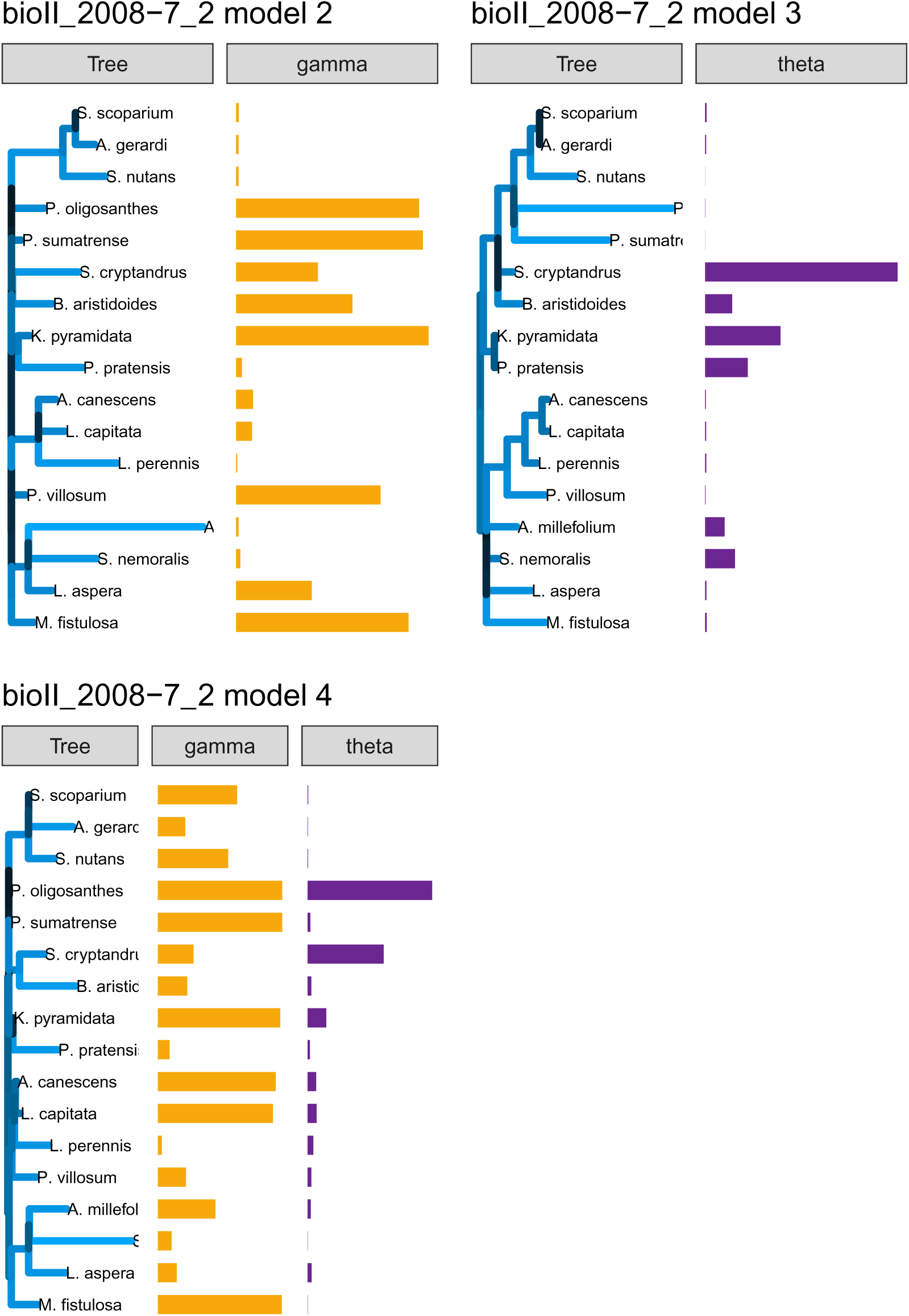
Fit for the phylogenetic tree and associated parameter vectors (Gamma and Theta) for models 2-4. The length and color of the branches correspond to increased strength in competition between the species in the clade and the size of the vector corresponds to the fitted value for the corresponding species in the tree. bioII refers to the Biodiversity II experiment (Tilman et al. (2001)); cadotte is from Cadotte (2013); and vr refers to the Wageningen experiment from Van Ruijven and Berendse (2010).

**Figure 35:**
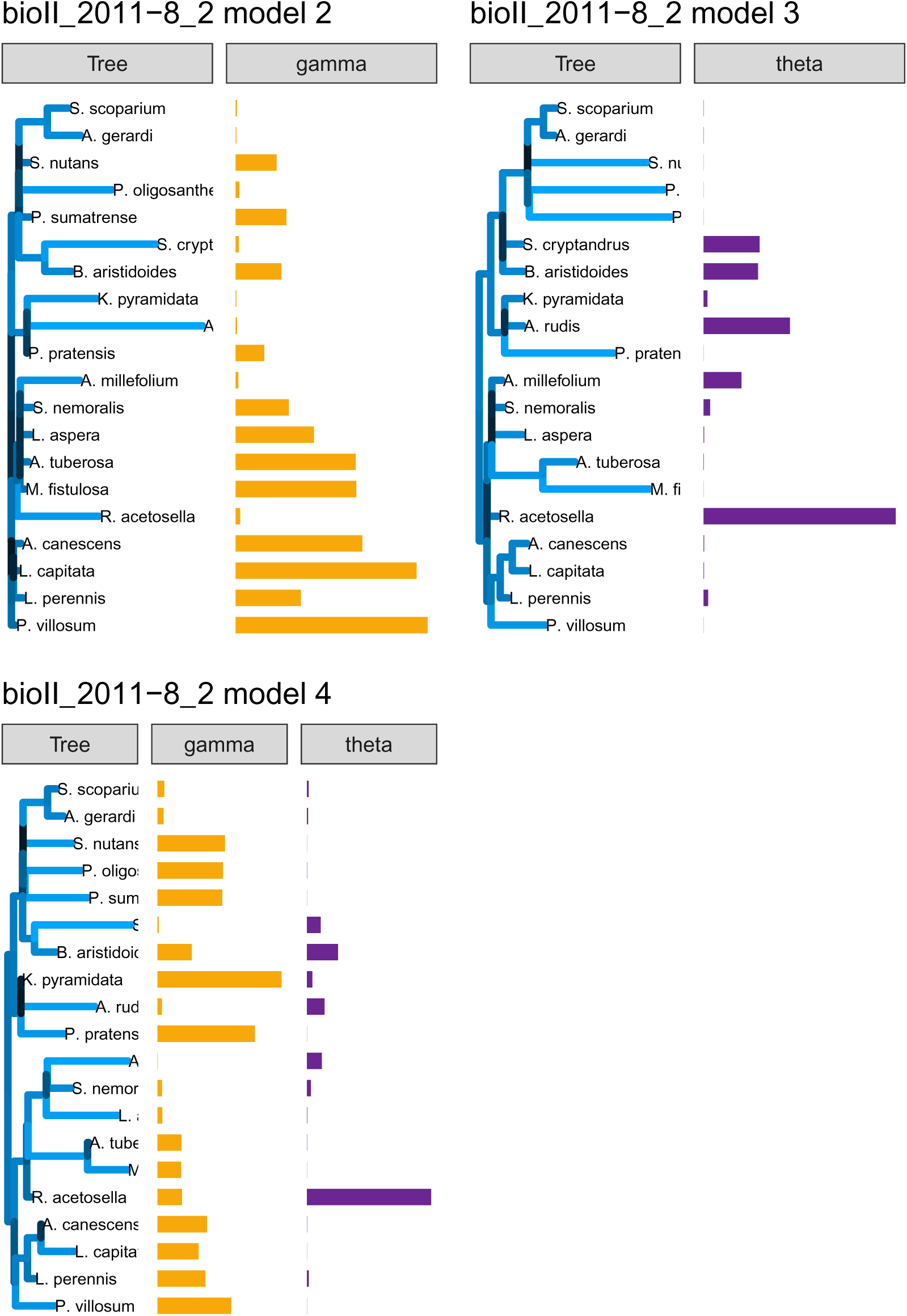
Fit for the phylogenetic tree and associated parameter vectors (Gamma and Theta) for models 2-4. The length and color of the branches correspond to increased strength in competition between the species in the clade and the size of the vector corresponds to the fitted value for the corresponding species in the tree. bioII refers to the Biodiversity II experiment (Tilman et al. (2001)); cadotte is from Cadotte (2013); and vr refers to the Wageningen experiment from Van Ruijven and Berendse (2010).

**Figure 36:**
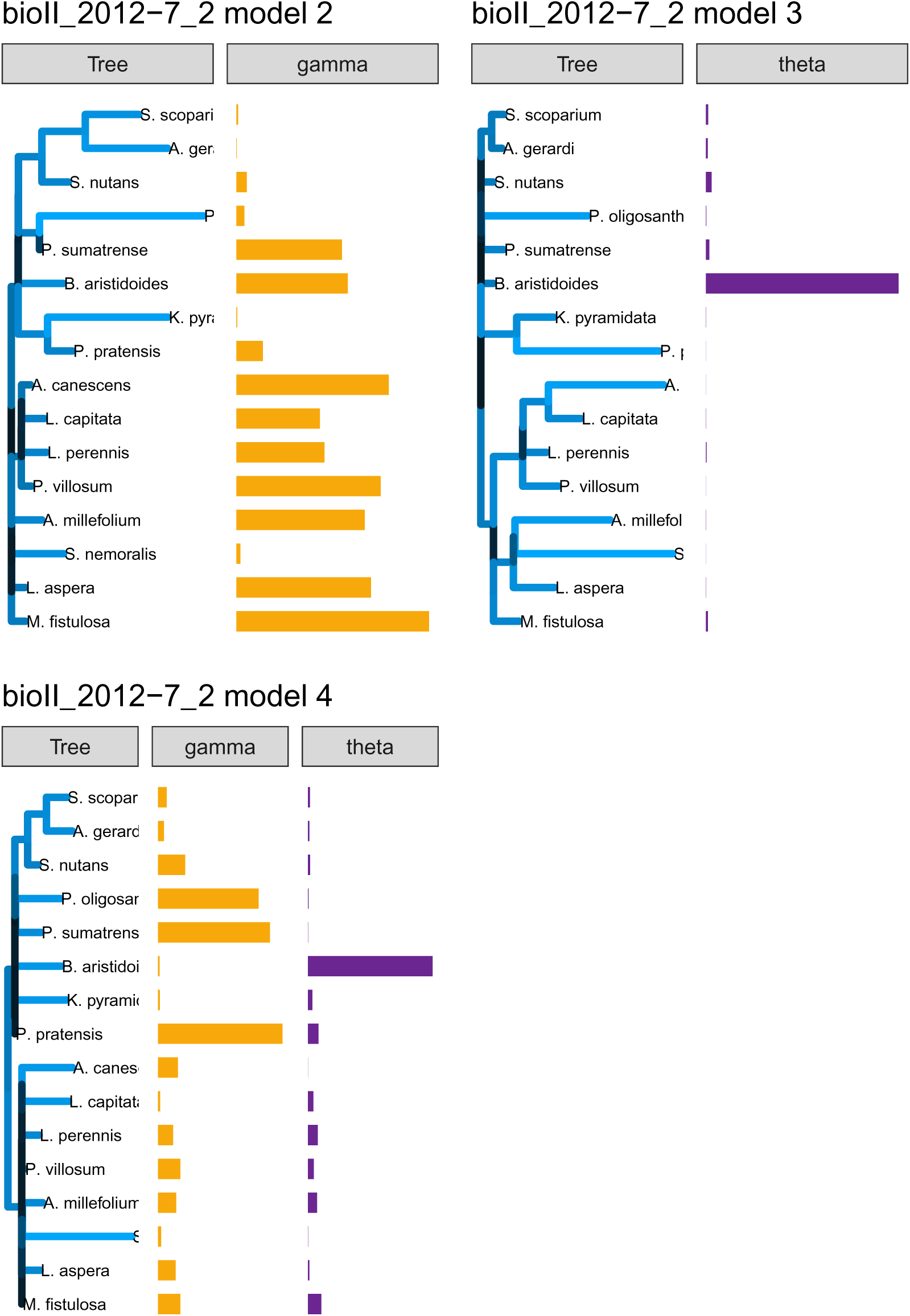
Fit for the phylogenetic tree and associated parameter vectors (Gamma and Theta) for models 2-4. The length and color of the branches correspond to increased strength in competition between the species in the clade and the size of the vector corresponds to the fitted value for the corresponding species in the tree. bioII refers to the Biodiversity II experiment (Tilman et al. (2001)); cadotte is from Cadotte (2013); and vr refers to the Wageningen experiment from Van Ruijven and Berendse (2010).

**Figure 37:**
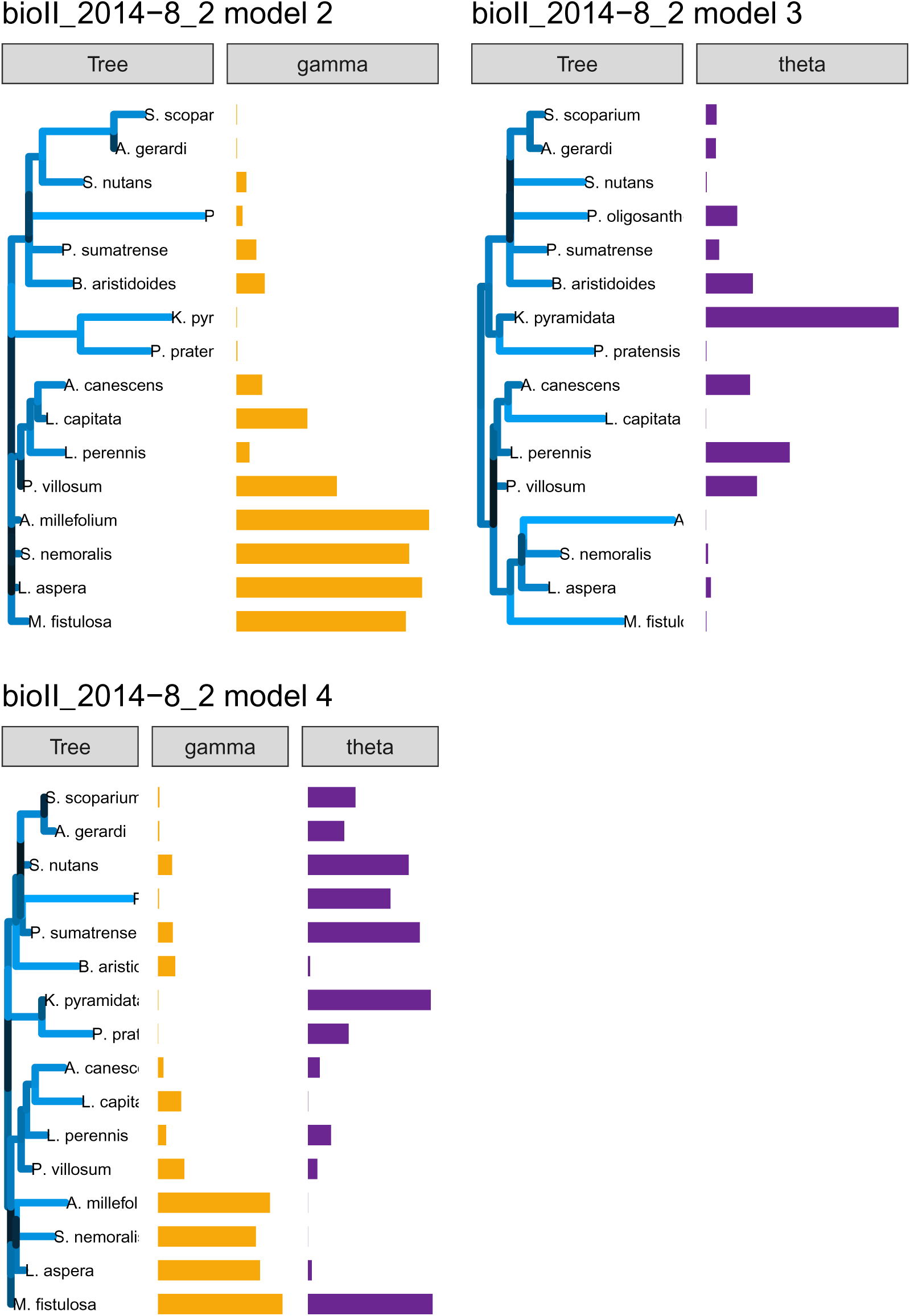
Fit for the phylogenetic tree and associated parameter vectors (Gamma and Theta) for models 2-4. The length and color of the branches correspond to increased strength in competition between the species in the clade and the size of the vector corresponds to the fitted value for the corresponding species in the tree. bioII refers to the Biodiversity II experiment (Tilman et al. (2001)); cadotte is from Cadotte (2013); and vr refers to the Wageningen experiment from Van Ruijven and Berendse (2010).

**Figure 38:**
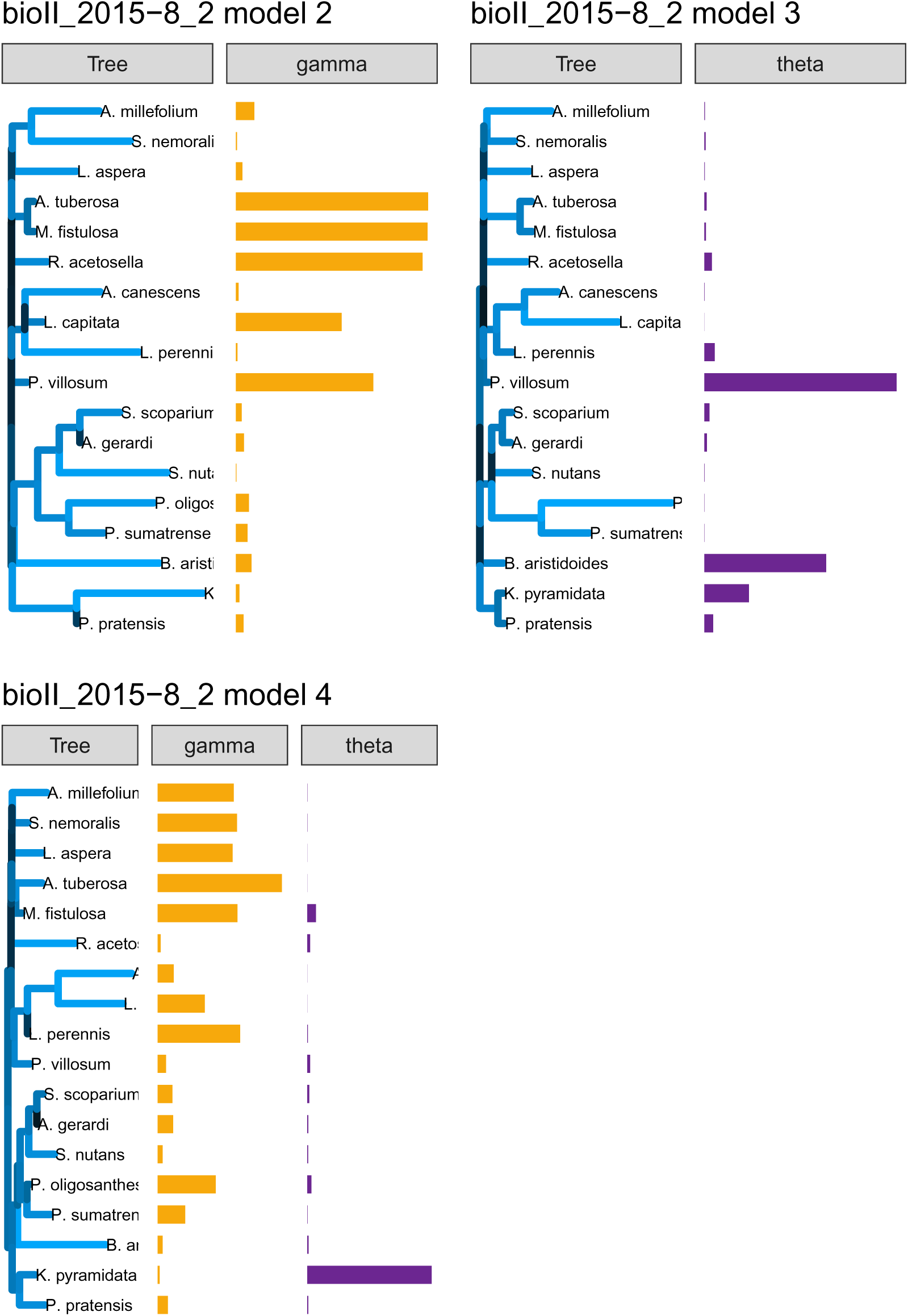
Fit for the phylogenetic tree and associated parameter vectors (Gamma and Theta) for models 2-4. The length and color of the branches correspond to increased strength in competition between the species in the clade and the size of the vector corresponds to the fitted value for the corresponding species in the tree. bioII refers to the Biodiversity II experiment (Tilman et al. (2001)); cadotte is from Cadotte (2013); and vr refers to the Wageningen experiment from Van Ruijven and Berendse (2010).

**Figure 39:**
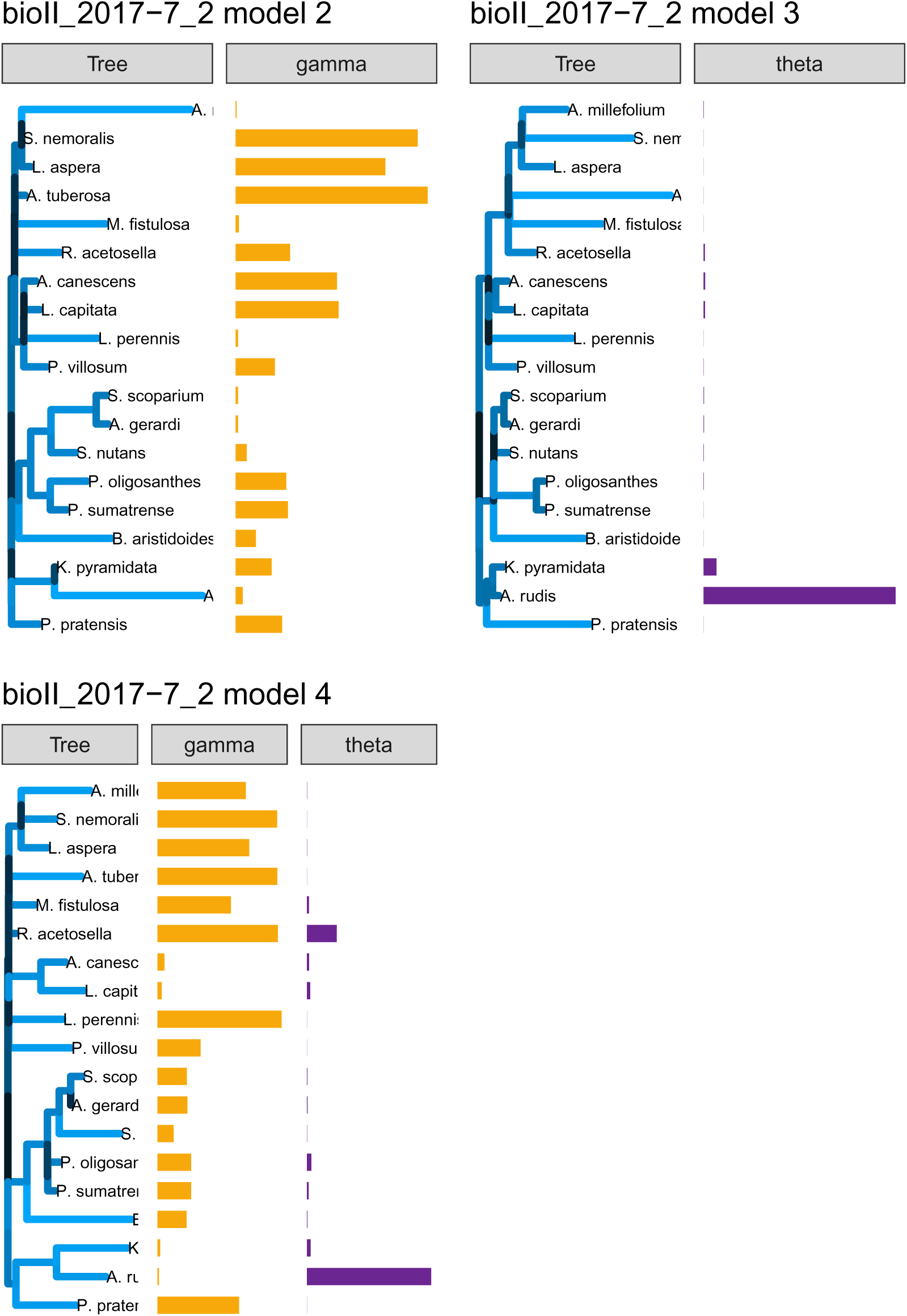
Fit for the phylogenetic tree and associated parameter vectors (Gamma and Theta) for models 2-4. The length and color of the branches correspond to increased strength in competition between the species in the clade and the size of the vector corresponds to the fitted value for the corresponding species in the tree. bioII refers to the Biodiversity II experiment (Tilman et al. (2001)); cadotte is from Cadotte (2013); and vr refers to the Wageningen experiment from Van Ruijven and Berendse (2010).

**Figure 40:**
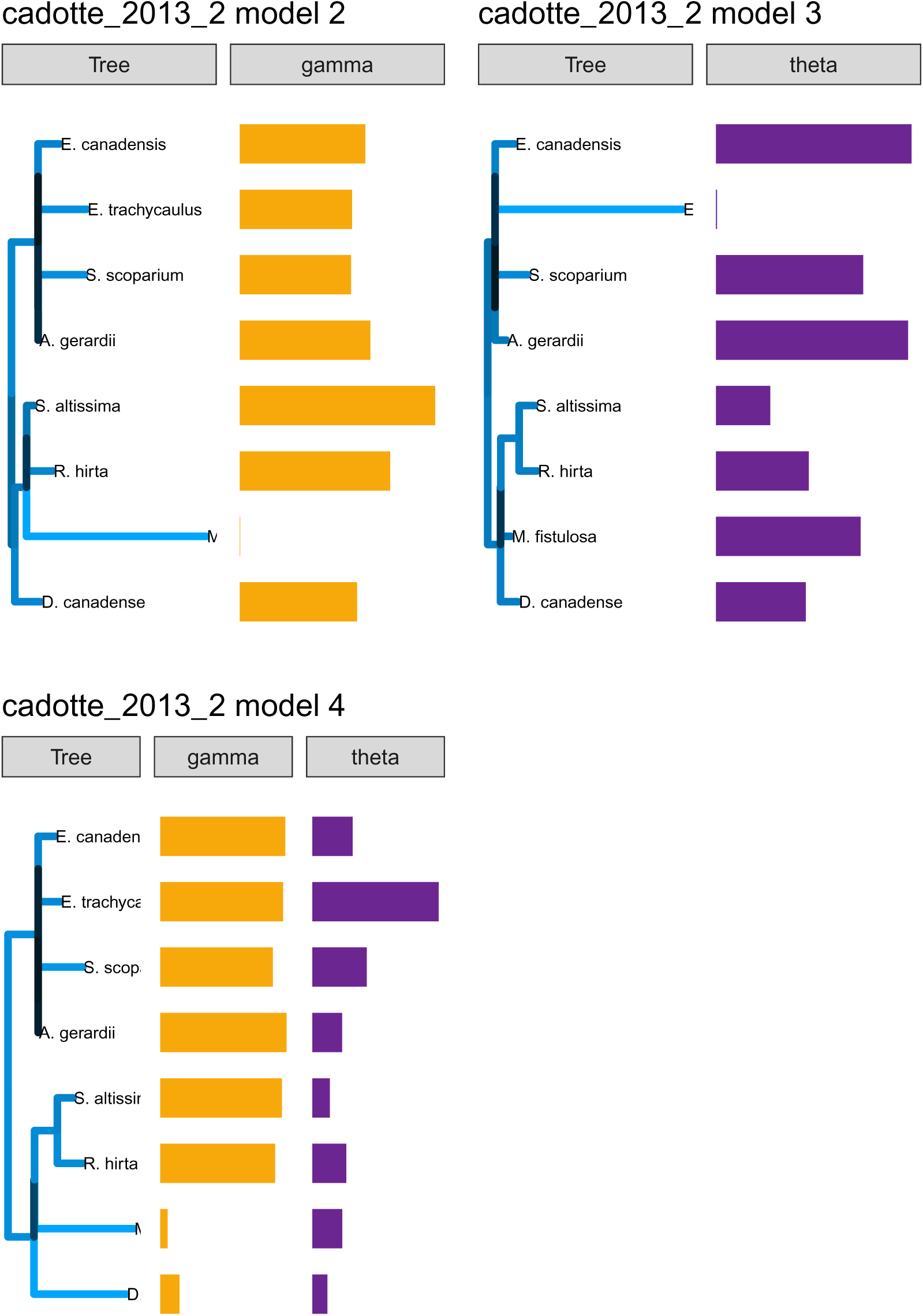
Fit for the phylogenetic tree and associated parameter vectors (Gamma and Theta) for models 2-4. The length and color of the branches correspond to increased strength in competition between the species in the clade and the size of the vector corresponds to the fitted value for the corresponding species in the tree. bioII refers to the Biodiversity II experiment (Tilman et al. (2001)); cadotte is from Cadotte (2013); and vr refers to the Wageningen experiment from Van Ruijven and Berendse (2010).

**Figure 41:**
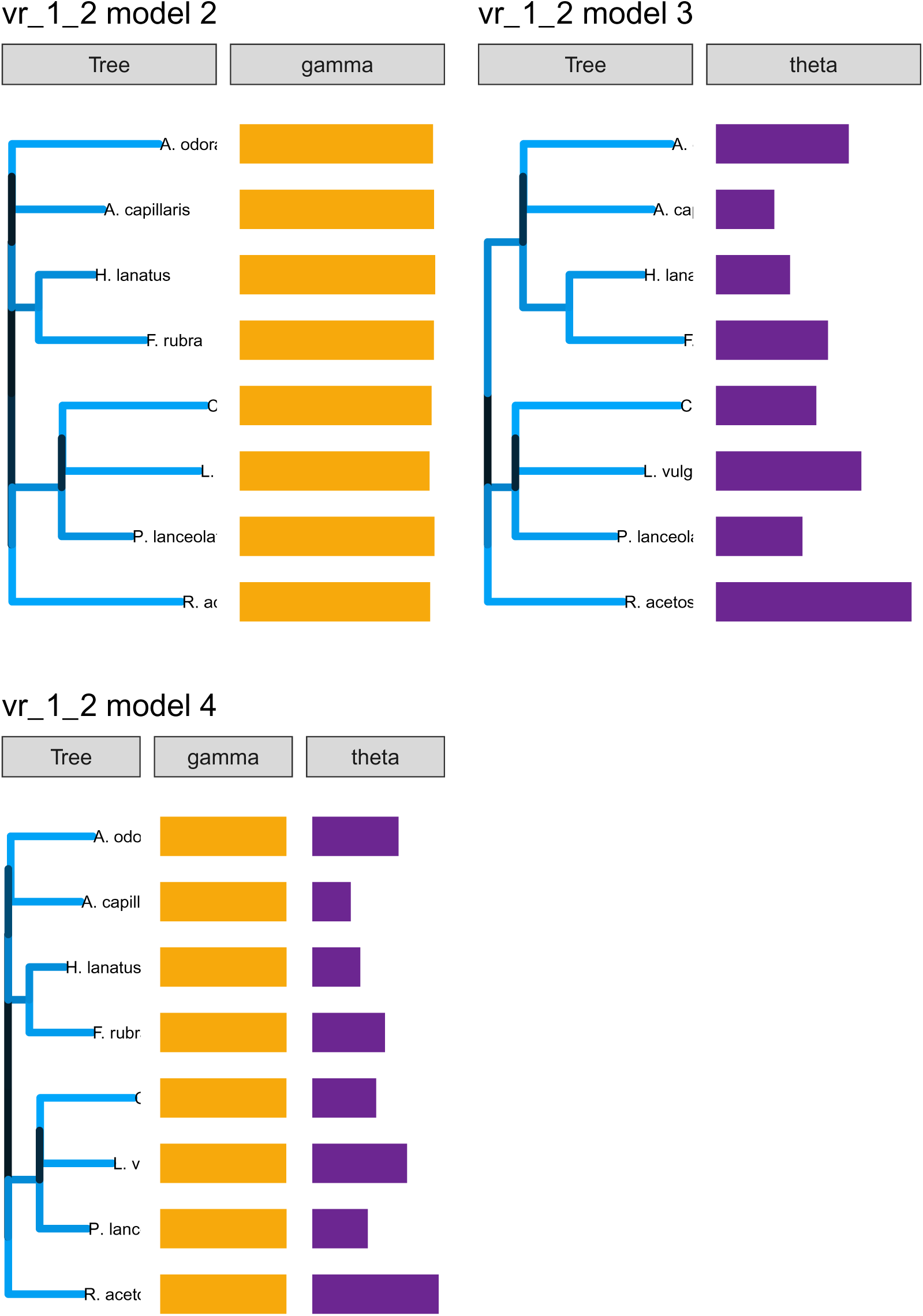
Fit for the phylogenetic tree and associated parameter vectors (Gamma and Theta) for models 2-4. The length and color of the branches correspond to increased strength in competition between the species in the clade and the size of the vector corresponds to the fitted value for the corresponding species in the tree. bioII refers to the Biodiversity II experiment (Tilman et al. (2001)); cadotte is from Cadotte (2013); and vr refers to the Wageningen experiment from Van Ruijven and Berendse (2010).

**Figure 42:**
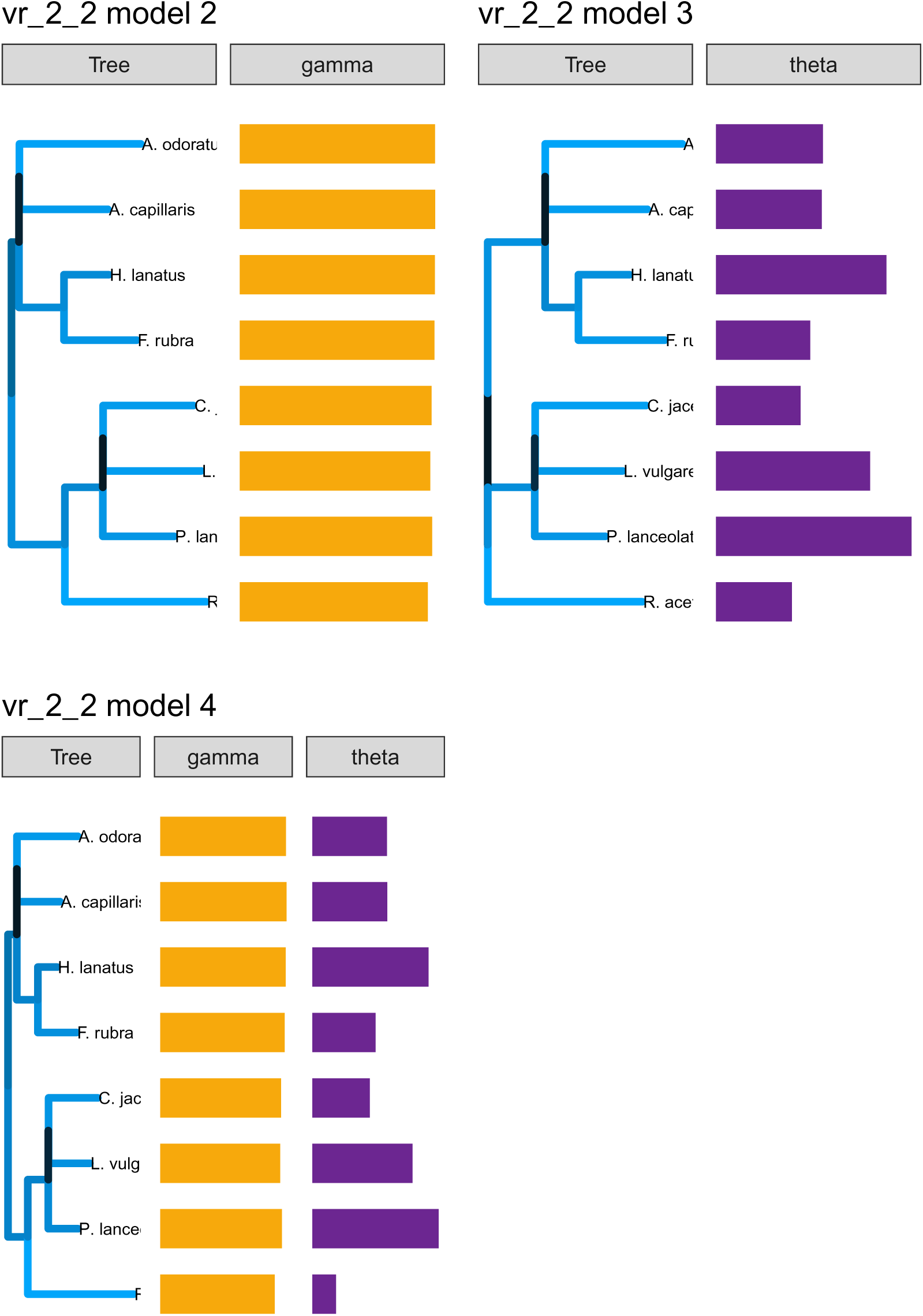
Fit for the phylogenetic tree and associated parameter vectors (Gamma and Theta) for models 2-4. The length and color of the branches correspond to increased strength in competition between the species in the clade and the size of the vector corresponds to the fitted value for the corresponding species in the tree. bioII refers to the Biodiversity II experiment (Tilman et al. (2001)); cadotte is from Cadotte (2013); and vr refers to the Wageningen experiment from Van Ruijven and Berendse (2010).

**Figure 43:**
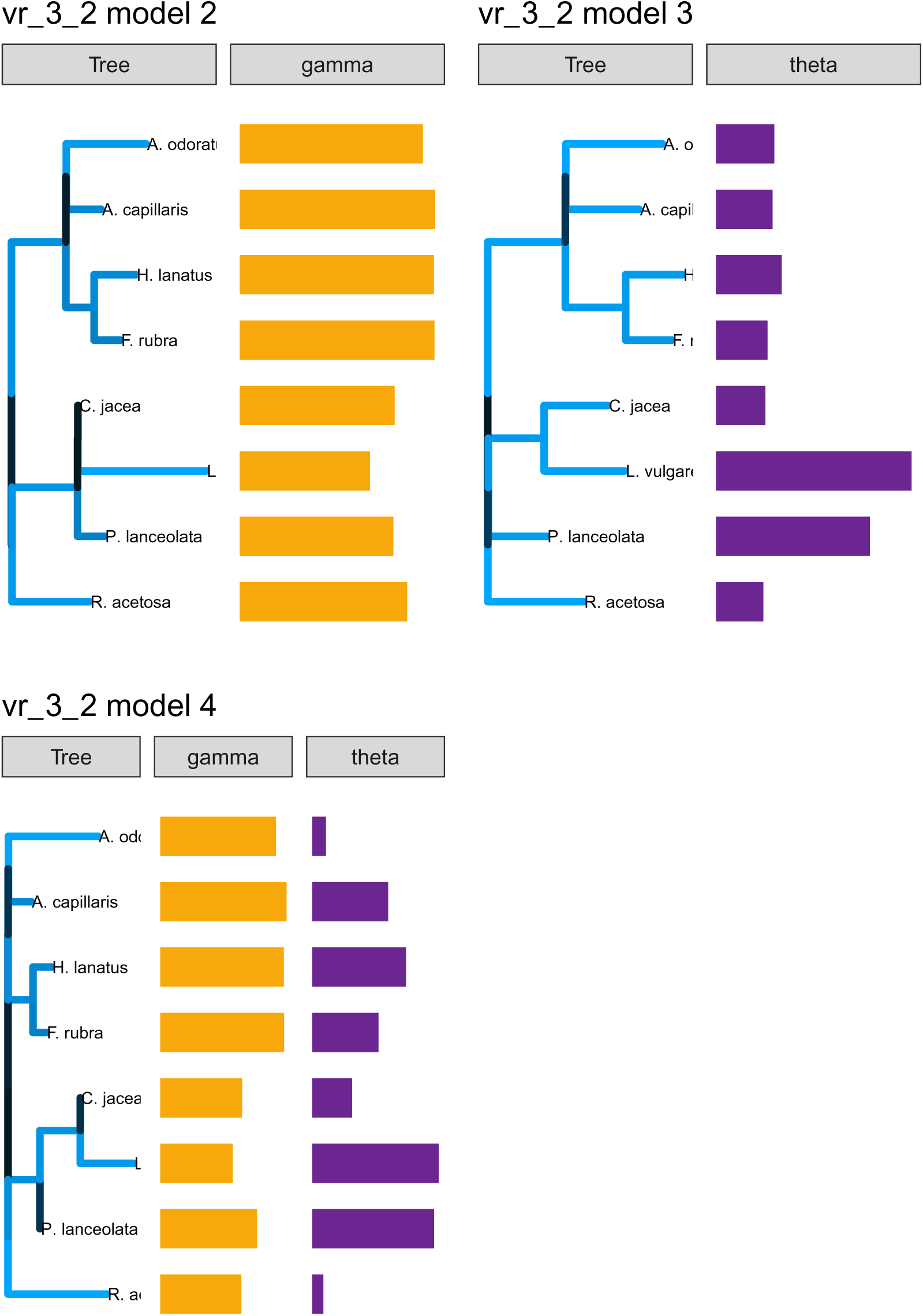
Fit for the phylogenetic tree and associated parameter vectors (Gamma and Theta) for models 2-4. The length and color of the branches correspond to increased strength in competition between the species in the clade and the size of the vector corresponds to the fitted value for the corresponding species in the tree. bioII refers to the Biodiversity II experiment (Tilman et al. (2001)); cadotte is from Cadotte (2013); and vr refers to the Wageningen experiment from Van Ruijven and Berendse (2010).

**Figure 44:**
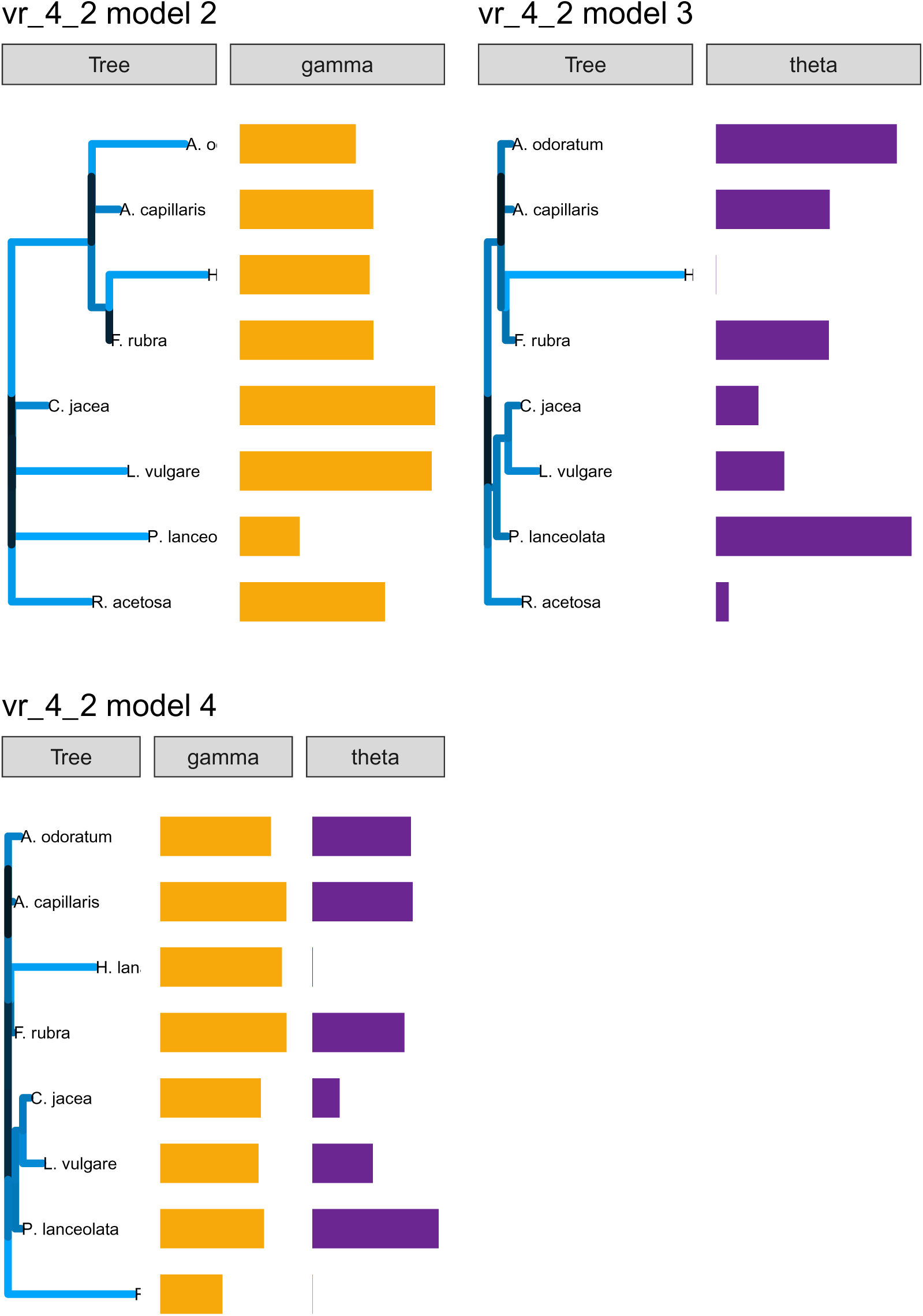
Fit for the phylogenetic tree and associated parameter vectors (Gamma and Theta) for models 2-4. The length and color of the branches correspond to increased strength in competition between the species in the clade and the size of the vector corresponds to the fitted value for the corresponding species in the tree. bioII refers to the Biodiversity II experiment (Tilman et al. (2001)); cadotte is from Cadotte (2013); and vr refers to the Wageningen experiment from Van Ruijven and Berendse (2010).

**Figure 45:**
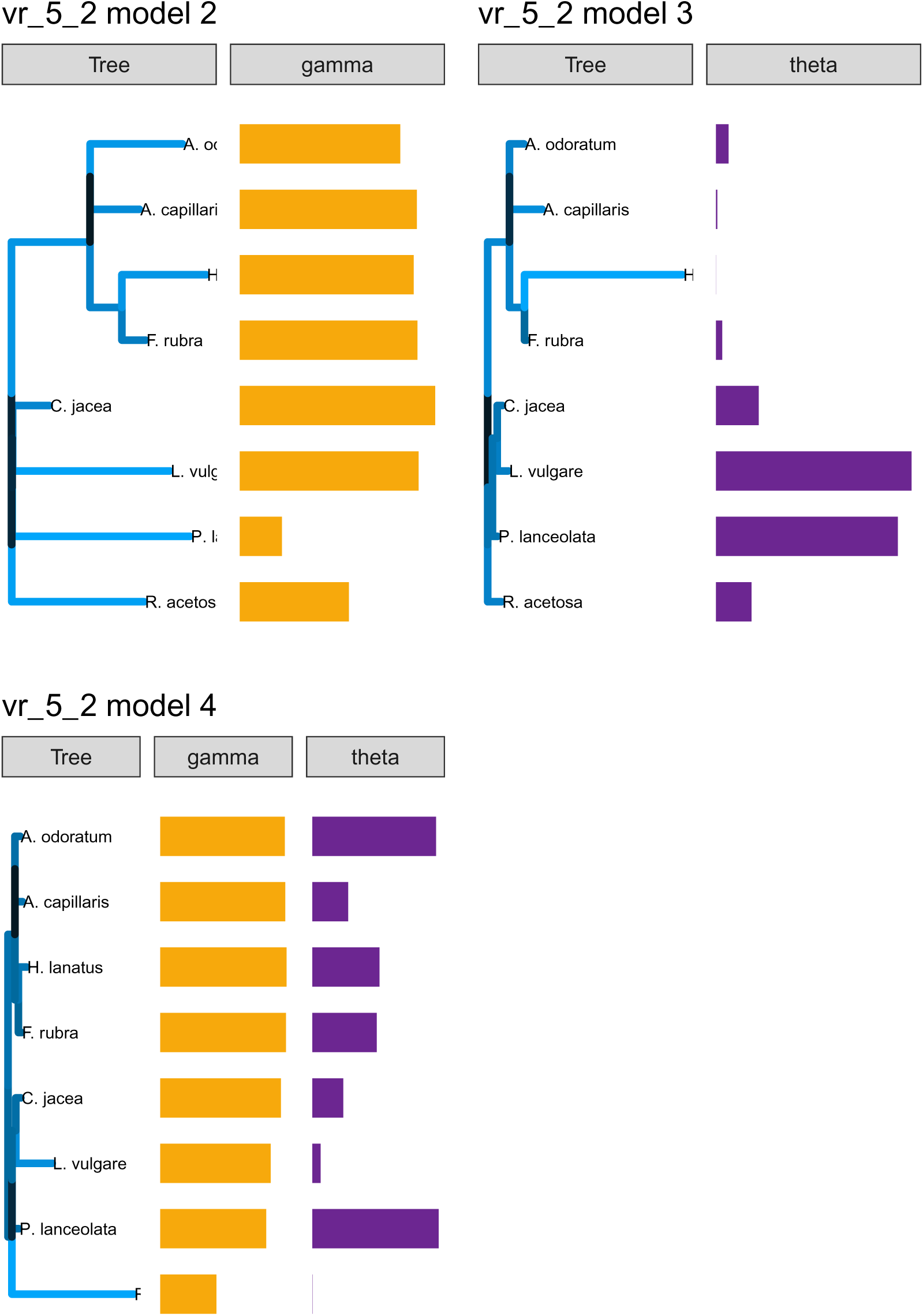
Fit for the phylogenetic tree and associated parameter vectors (Gamma and Theta) for models 2-4. The length and color of the branches correspond to increased strength in competition between the species in the clade and the size of the vector corresponds to the fitted value for the corresponding species in the tree. bioII refers to the Biodiversity II experiment (Tilman et al. (2001)); cadotte is from Cadotte (2013); and vr refers to the Wageningen experiment from Van Ruijven and Berendse (2010).

**Figure 46:**
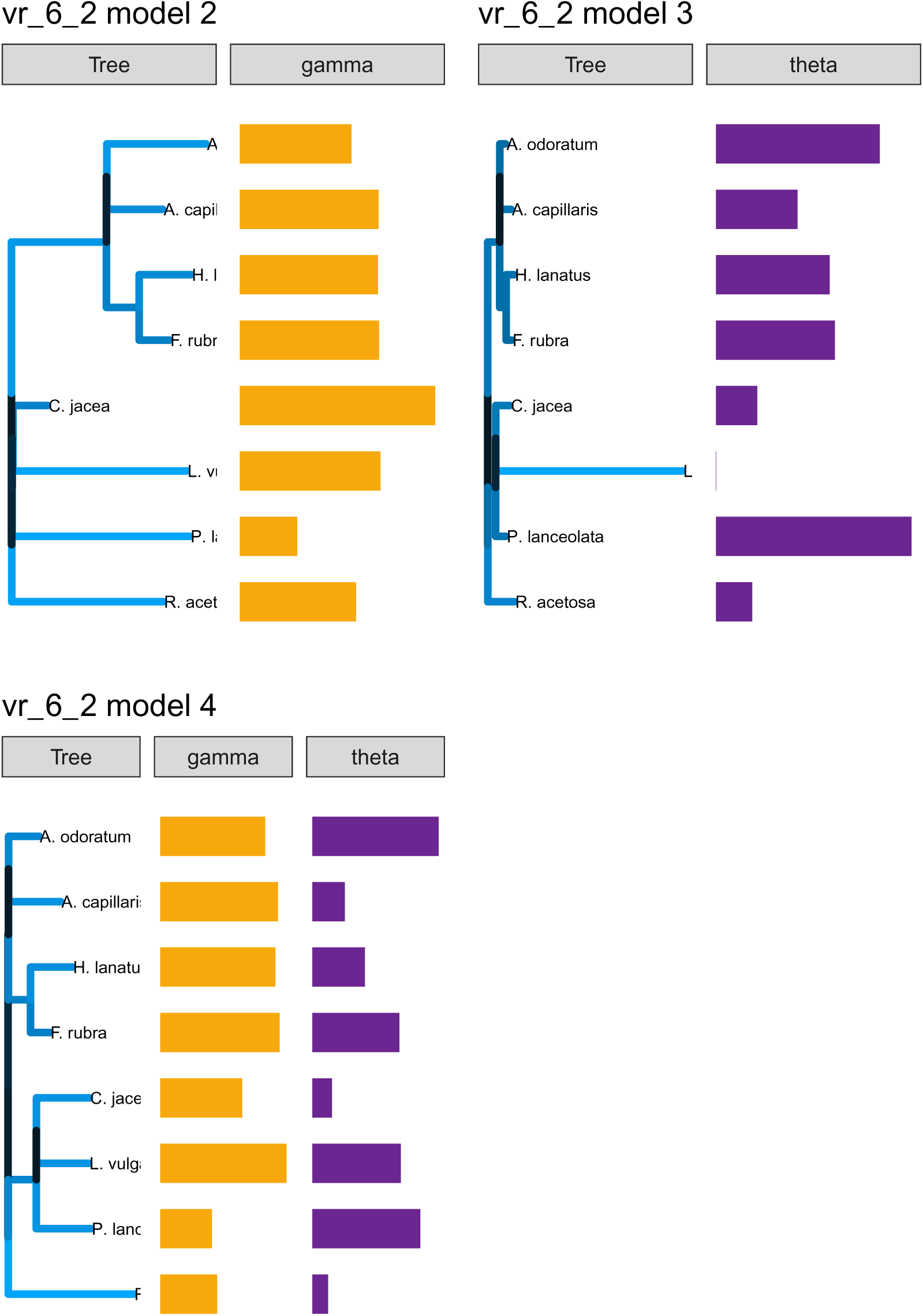
Fit for the phylogenetic tree and associated parameter vectors (Gamma and Theta) for models 2-4. The length and color of the branches correspond to increased strength in competition between the species in the clade and the size of the vector corresponds to the fitted value for the corresponding species in the tree. bioII refers to the Biodiversity II experiment (Tilman et al. (2001)); cadotte is from Cadotte (2013); and vr refers to the Wageningen experiment from Van Ruijven and Berendse (2010).

**Figure 47:**
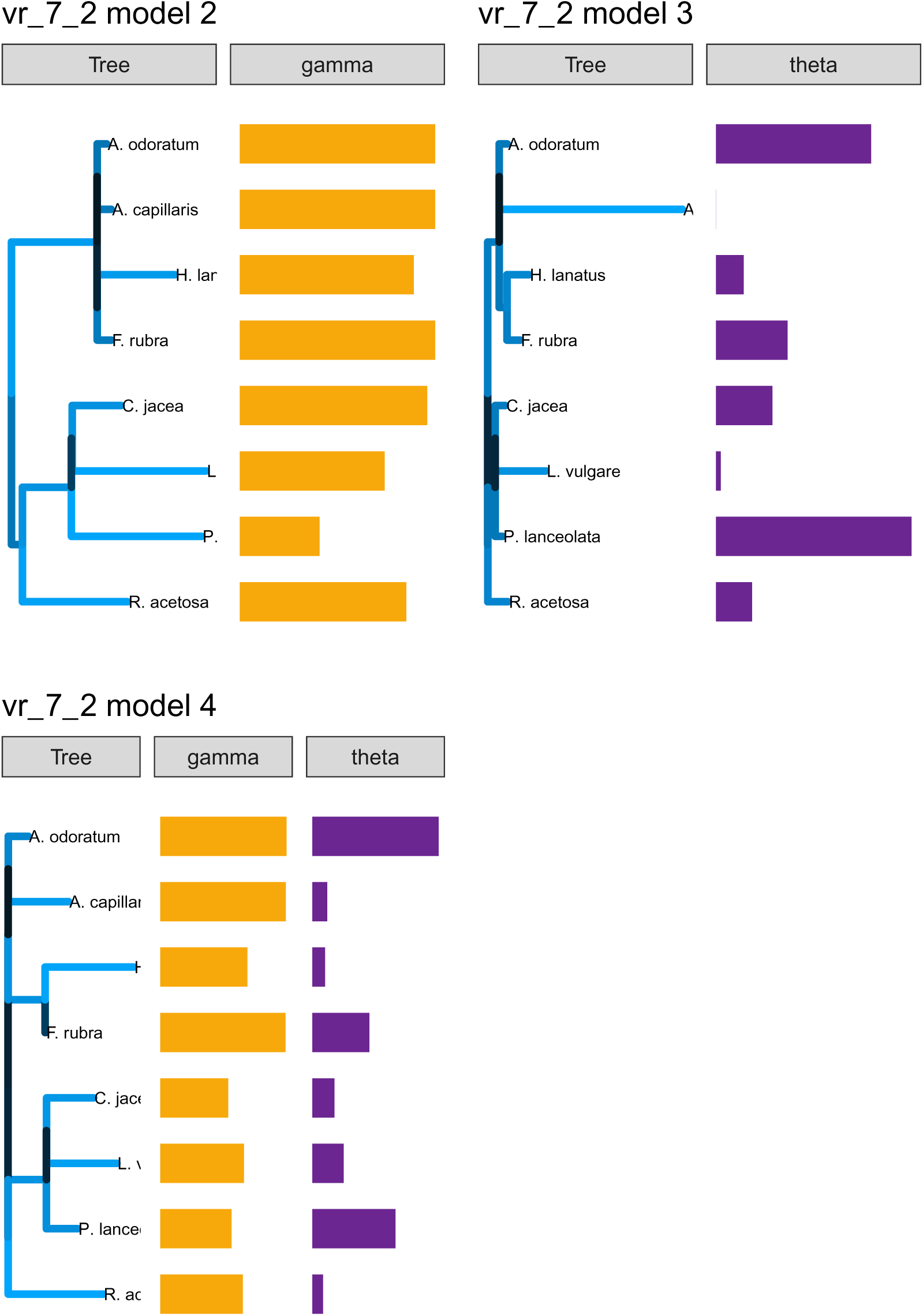
Fit for the phylogenetic tree and associated parameter vectors (Gamma and Theta) for models 2-4. The length and color of the branches correspond to increased strength in competition between the species in the clade and the size of the vector corresponds to the fitted value for the corresponding species in the tree. bioII refers to the Biodiversity II experiment (Tilman et al. (2001)); cadotte is from Cadotte (2013); and vr refers to the Wageningen experiment from Van Ruijven and Berendse (2010).

**Figure 48:**
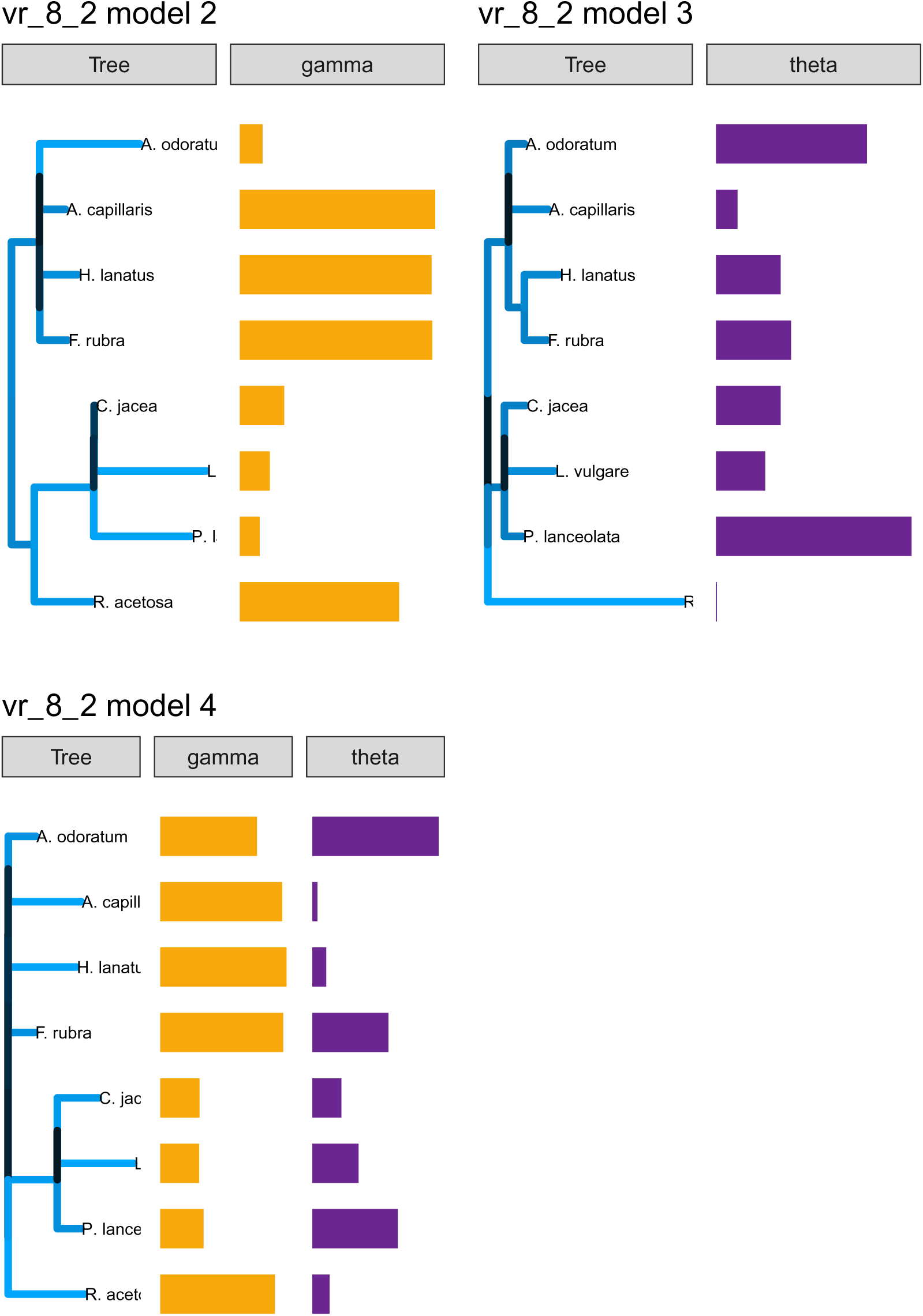
Fit for the phylogenetic tree and associated parameter vectors (Gamma and Theta) for models 2-4. The length and color of the branches correspond to increased strength in competition between the species in the clade and the size of the vector corresponds to the fitted value for the corresponding species in the tree. bioII refers to the Biodiversity II experiment (Tilman et al. (2001)); cadotte is from Cadotte (2013); and vr refers to the Wageningen experiment from Van Ruijven and Berendse (2010).

**Figure 49:**
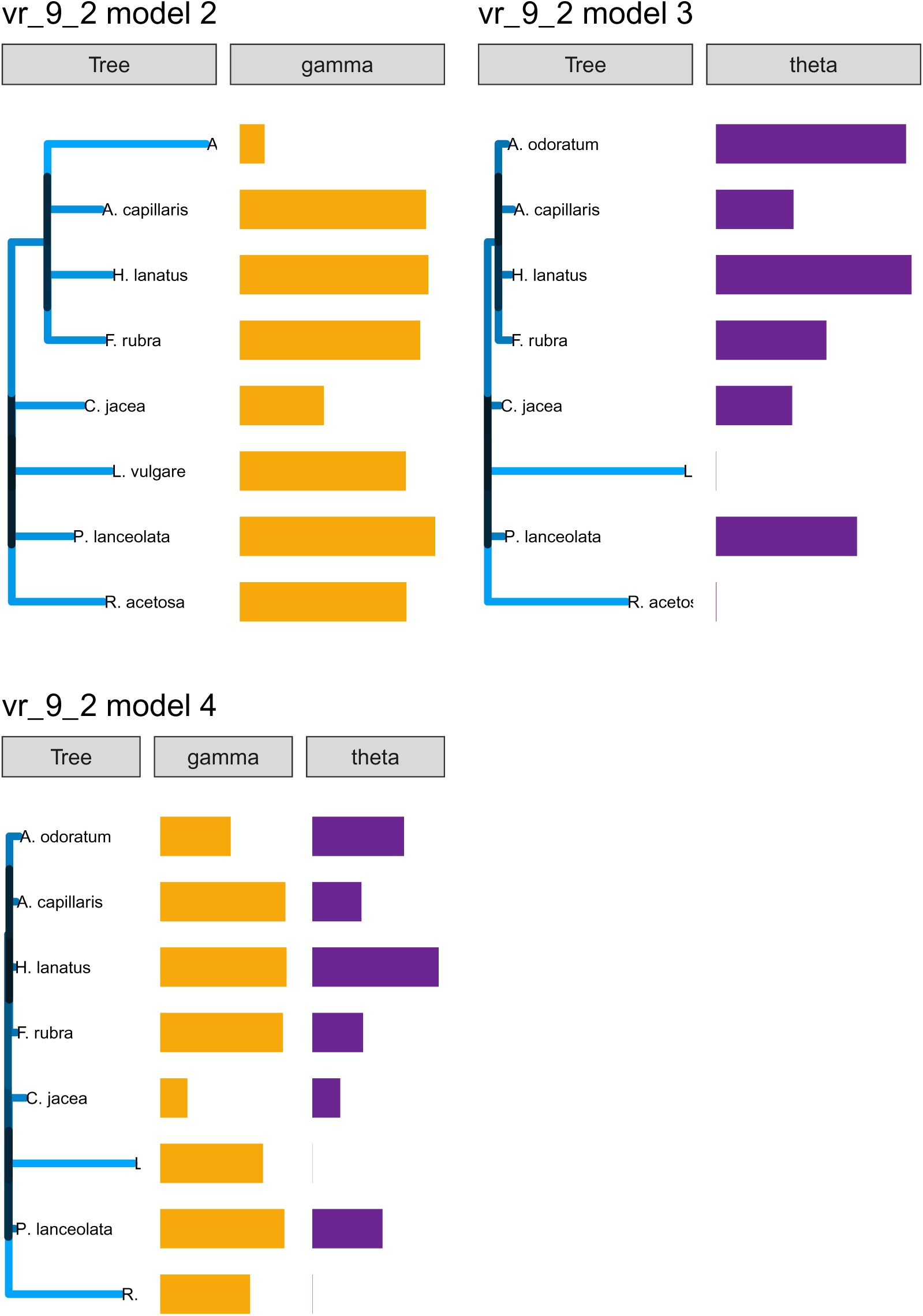
Fit for the phylogenetic tree and associated parameter vectors (Gamma and Theta) for models 2-4. The length and color of the branches correspond to increased strength in competition between the species in the clade and the size of the vector corresponds to the fitted value for the corresponding species in the tree. bioII refers to the Biodiversity II experiment (Tilman et al. (2001)); cadotte is from Cadotte (2013); and vr refers to the Wageningen experiment from Van Ruijven and Berendse (2010).

## Notes

### Competing Interest Statement

The authors have declared no competing interest.

### Summary of Updates

Acknowledgments updated

